# An egg-derived sulfated N-Acetyllactosamine glycan is an antigenic decoy of influenza virus vaccines

**DOI:** 10.1101/2021.03.16.435673

**Authors:** Jenna J. Guthmiller, Henry A. Utset, Carole Henry, Lei Li, Nai-Ying Zheng, Weina Sun, Marcos Costa Vieira, Seth Zost, Min Huang, Scott E. Hensley, Sarah Cobey, Peter Palese, Patrick C. Wilson

**Affiliations:** Department of Medicine, Section of Rheumatology, University of Chicago, Chicago, IL 60637, USA; Department of Microbiology, Icahn School of Medicine at Mount Sinai, New York, NY 10029, USA; Department of Ecology and Evolution, University of Chicago, Chicago, IL 60637, USA; Department of Microbiology, Perelman School of Medicine, University of Pennsylvania, Philadelphia, PA 19104, USA

**Author notes:** Address correspondence to: Jenna J. Guthmiller, 924 E. 57^th^ St. BSLC R422, Chicago, IL 60637., Patrick C. Wilson, 924 E. 57^th^ St. BSLC R420, Chicago, IL 60637. Moderna Inc., Cambridge, MA 02139, USA.

## Abstract

Influenza viruses grown in eggs for the purposes of vaccine generation often acquire mutations during egg adaptation or possess differential glycosylation patterns than viruses circulating amongst humans. Here, we report that seasonal influenza virus vaccines possess an egg-derived sulfated N-acetyllactosamine (LacNAc) that is an antigenic decoy. Half of subjects that received an egg-grown vaccine mounted an antibody response against this egg-derived antigen. Egg-binding monoclonal antibodies specifically bind viruses grown in eggs, but not viruses grown in other chicken derived cells, suggesting only egg-grown vaccines can induce anti-LacNAc antibodies. Notably, antibodies against the sulfated LacNAc utilized a restricted antibody repertoire and possessed features of natural antibodies, as most antibodies were IgM and have simple heavy chain complementarity determining region 3. By analyzing a public dataset of influenza virus vaccine induced plasmablasts, we discovered egg-binding public clonotypes that were shared across studies. Together, this study shows that egg-grown vaccines can induce antibodies against an egg-associated glycan, which may divert the host immune response away from protective epitopes.

## Introduction

Influenza viruses have been historically grown in embryonated chicken eggs as a way to culture large quantities of virus, and as a result, most influenza virus vaccines are still generated using viruses grown in eggs. However, this process has the potential to change the immunogenicity of the virus, as the viruses may mutate their major surface glycoproteins hemagglutinin (HA) and neuraminidase (NA) to increase infectivity in eggs (1–5). Moreover, influenza viruses grown in eggs are often less immunogenic than viruses grown in mammalian cells (6–8) and have been shown to be less effective than mammalian cell-based influenza vaccines and recombinantly expressed HA vaccines (9, 10). Due to the inherent difference in avian versus mammalian glycosylation patterns, egg-grown vaccines may lack certain glycans that would be expressed on influenza viruses transmitted among humans. Notably, vaccine effectiveness against recent H3N2 viruses may be reduced due to the lack of a glycan on HA of H3N2 viruses grown in eggs (1).

However, slight differences in viral sequences and glycosylation patterns of HA do not fully explain why vaccine effectiveness is low, as serum from vaccinated subjects can have similar antibody titers against egg adapted strains and viruses circulating in the population (11). Poor immunogenicity against HA may explain reductions in vaccine effectiveness rather than egg-adapted mutations (11). It is possible that egg-grown vaccines are preferentially inducing antibodies against non-protective viral antigens, therefore reducing vaccine effectiveness and seroconversion against protective epitopes on the HA head domain. Similar to subjects receiving egg-grown vaccines (12, 13), HA-reactive antibodies induced by vaccines grown in mammalian cells and insect cells largely induced antibodies mostly targeting the head domain of HA (14). However, whether different vaccine platforms induced antibodies against distinct influenza virus antigens, other than HA, is not known. As internal antigens, such as the nucleoprotein (NP), were shown to provide limited protection against infection (12), it remains to be determined how different vaccine formulations drive antibodies against distinct protective and non-protective antigens.

To address whether these egg-grown influenza virus vaccines induced antibodies against potentially non-protective antigens, we cloned monoclonal antibodies (mAbs) from plasmablasts (PBs), a transient antibody secreting cell population, isolated from subjects following vaccination with egg-grown influenza virus vaccines. We show that 50% of subjects generated a PB response against an egg-derived antigen present in the vaccine. Subjects that mounted a response against the egg-associated antigen seroconverted against HA to similar levels as subjects that did not mount an anti-egg response, indicating egg-grown vaccines did not reduce overall secreted antibody responses against HA. We identified that the egg-derived antigen was a sulfated N-acetyllactosamine (LacNAc) glycan and was only present in viruses grown in the allantois of eggs, but not in viruses grown in chicken embryo cell line or primary chicken fibroblasts. Antibodies binding the egg-derived gylcan utilized a restricted repertoire and resembled natural antibodies, as antibodies were largely IgM and had short heavy chain complementarity determining region 3 (H-CDR3). Moreover, we identified that egg-binding antibodies identified in our study were public clonotypes, indicating the same antibodies were found across individual subjects that had been vaccinated with an egg-grown vaccine. Together, our study shows that egg-grown vaccines can induce antibodies against an egg related glycan and that these glycan-binding mAbs resemble those produced by innate-like B cells.

## Results

### Influenza virus vaccination induces antibodies against an egg-associated antigen

To address the antigen specificity of memory B cells recalled by egg-grown influenza virus vaccines, we generated mAbs from sorted PBs 7 days following vaccination. The transient PB population are highly specific to the components of the vaccine and are recalled from pre-existing memory B cells (15–17). We focused our studies on mAbs generated from subjects following vaccination with the 2009 monovalent pandemic H1N1 inactivated influenza virus vaccine (MIV) and the 2010 trivalent inactivated influenza virus vaccine (TIV). From the vaccine induced PBs, we found that 75% of mAbs bound HA (Figure S1A and B). Notably, 27 mAbs generated from multiple subjects receiving either of the egg-grown vaccines bound all influenza virus strains tested (Figure 1A; Table S1). To confirm these mAbs were specific to influenza viruses and not an artifact of vaccine preparation in eggs, we tested mAb binding to A/California/7/2009 H1N1 virus grown in eggs or in mammalian Madin-Darby Canine Kidney (MCDK) cells. Strikingly, these mAbs only bound to A/California/7/2009 grown in eggs, but not virus propagated in mammalian cells (Figure 1B and C), indicating these broadly reactive mAbs were binding to an egg-related antigen. Moreover, these broadly reactive mAbs bound to allantoic fluid from both uninfected eggs and A/California/7/2009 H1N1 infected eggs (Figure 1D-E), indicating these mAbs were specific to an egg associated antigen.

**Figure 1:**
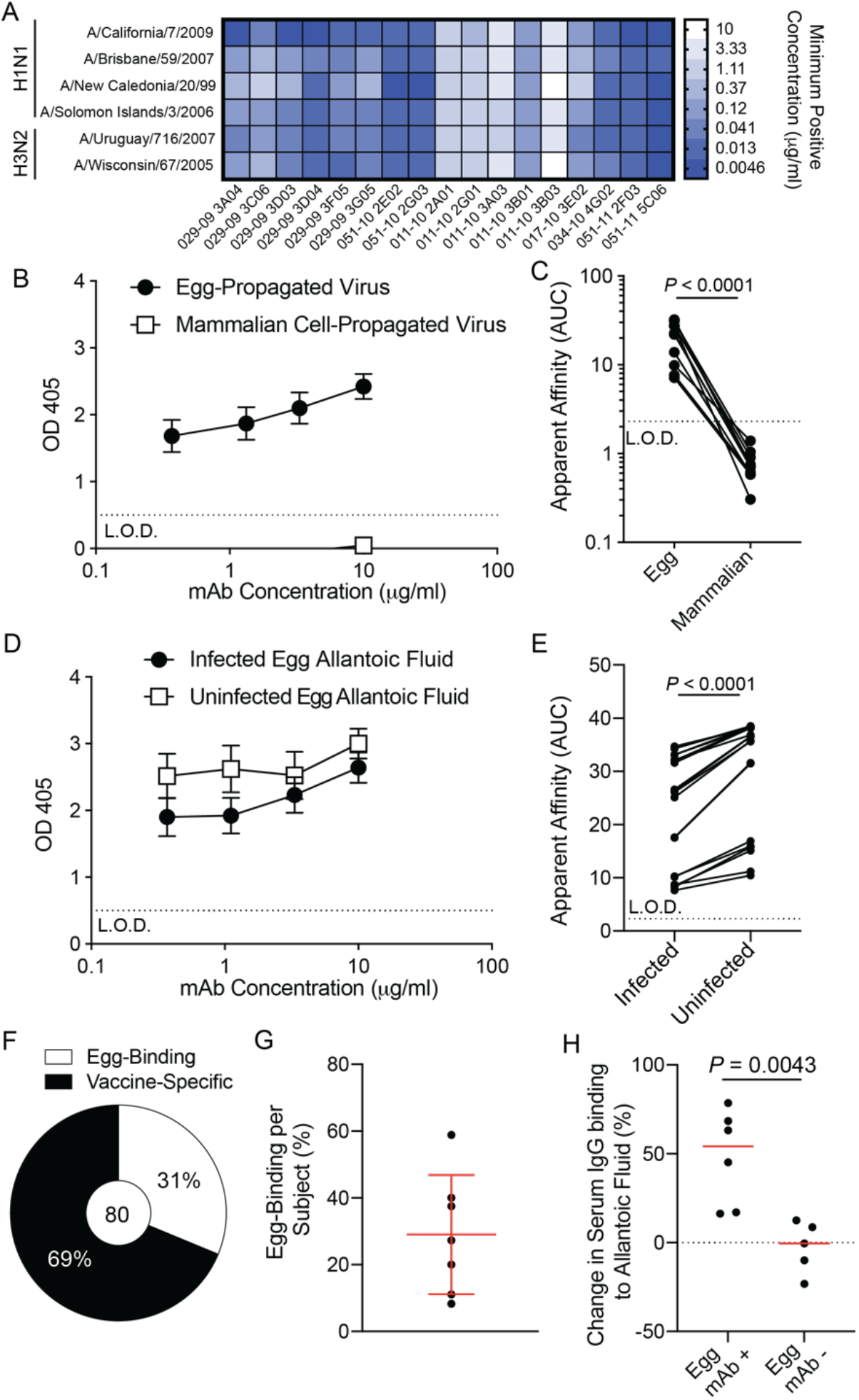
Identification of mAbs binding an egg-specific antigen. **A**, Heatmap of selected mAbs binding all influenza virus strains tested. Data are representative of all 27 identified antibodies. **B** and **C**, Broadly reactive mAbs (n=22) binding to egg-propagated and MDCK cell (mammalian) propagated A/California/7/2009 H1N1 (**B**) and apparent affinity, as calculated as area under the curve (AUC), of mAb binding (**C**; n=11 mAbs). **D** and **E**, Broadly reactive mAb binding to A/California/7/2009 H1N1 infected allantoic fluid and uninfected allantoic fluid (**D**) and AUC of mAb binding (**E**; n=21). **F**, Proportion of mAbs from subjects with egg mAbs that are egg-binding or are specific to the vaccine. **G**, Proportion of total mAbs that are egg-binding per subject. **H**, Serum was isolated from subjects with or without isolated egg-binding mAbs before and 14-21 days after vaccination. Relative change in serum IgG binding to uninfected allantoic fluid represented as a percentage. Red line represented median. Data in **B**, **D**, and **G** are mean ± S.D. Data in **C** and **E** were analyzed using a two-tailed Wilcoxon matched-pairs signed rank test. Data in **h** were analyzed using a two-tailed Mann-Whitney test.

50% of all subjects analyzed generated egg-specific antibody responses (Figure S1C), with 1 out of 5 of the subjects receiving the MIV and 6 out of 9 of subjects receiving the TIV generating an egg-specific antibody response (Table S1). Within the subjects that mounted an antibody response against this egg-derived antigen, 31% of mAbs generated specifically bound the egg-derived antigen (Figure 1F). The range of antibodies per subject ranged from 8% to 58% of isolated vaccine induced PBs (Figure 1G). Additionally, we found that subjects that had egg-reactive mAbs had a larger fold increase in serum IgG responses against uninfected allantoic fluid relative to subjects who did not have detectable mAbs against the egg-associated antigen (Figure 1H). Despite this, subjects that mounted an antibody response against the egg-associated antigen had a similar fold increase in serum IgG titers against A/California/7/2009 recombinant HA and hemagglutination inhibition (HAI) titers against A/California/7/2009 H1N1, relative to subjects that did not generate a PB response against the egg antigen (Figure S1D-F). Together, these data indicate that some subjects following influenza virus vaccination generate an antibody response against an egg-derived antigen.

### Viruses grown in allantoic fluid, but not other parts of the egg, possess the egg antigen

Starting in 2018, the United States Center for Disease Control began recommending that people with egg allergies could receive egg-grown influenza virus vaccines, suggesting the major egg allergens were removed from the vaccine (https://www.cdc.gov/flu/prevent/egg-allergies.htm). Although subjects within our cohorts had not experienced an allergic response to influenza virus vaccination or reported a history of egg allergies, we next tested whether the identified egg-specific mAbs could bind to more recent inactivated influenza virus vaccines grown in eggs that lack the egg allergens. Notably, the egg-specific mAbs could bind old TIVs and quadrivalent inactivated influenza virus vaccines (QIVs) to a similar degree as recent egg-grown QIVs, including the 2020 Fluarix QIV (Figure 2A), indicating the egg-specific antigen identified still persists in recent egg-grown vaccines. Additionally, some QIVs are also made from viruses grown in mammalian MDCK cells (Flucelvax) and recombinant HA generated in insect cells (Flublok). The egg-binding mAbs specifically bound to QIV viruses grown in eggs, but not viruses grown in MDCK or recombinant HA grown in insect cells (Figure 2B). We also confirmed that these egg-binding mAbs could bind other viruses grown in the allantoic fluid of eggs, as these mAbs bound as strongly to Newcastle Disease Virus grown in eggs as it did to A/California/7/2009 H1N1 grown in eggs (Figure 2C). To understand how ubiquitous the egg antigen was across chicken cell- and egg-grown vaccines, we next tested for mAb binding to other vaccines grown in chicken cells. Notably, the mumps and measles viruses of the Measles/Mumps/Rubella vaccine (MMR) are both grown in a chicken embryo cell line and the rabies virus in Rabavert is grown in primary chicken fibroblasts. As a control, we also tested the egg binding mAbs for binding to non-egg or chicken grown vaccines including the Japanese Encephalitis Virus Vaccine (Ixiaro) grown in Vero cells and the Pneumovax 23 vaccine that contains purified capsular polysaccharides from 23 distinct *Streptococcus pneumoniae* serotypes. The egg-binding mAbs only bound to vaccines grown in eggs, but not to vaccines grown in a chicken embryo cell line (MMR), primary chicken fibroblasts (Rabavert), or vaccines not produced in eggs (Ixiaro and Pneumovax 23; Figure 2D). Together, these data reveal that the egg-associated antigen is only present in viruses grown in allantoic membrane, but not in those grown in cells isolated from chicken embryos or chicken fibroblasts.

**Figure 2:**
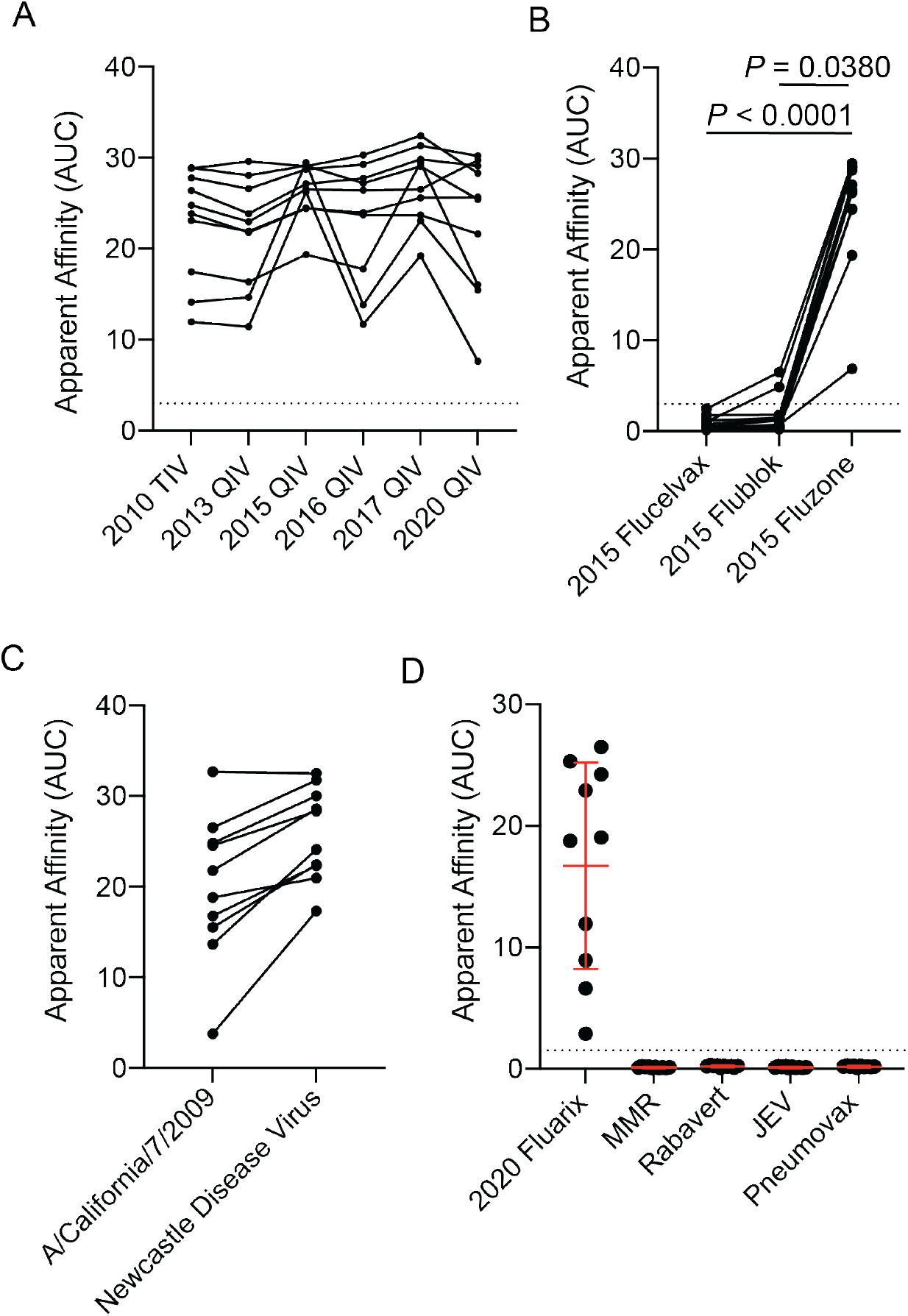
Binding is specific to viruses grown in eggs, but not chicken cells. **A**, Apparent affinity (AUC) of egg-specific mAbs binding to multiple years of influenza virus vaccines. Each line connects the same mAb binding a different vaccine (n=10 mAbs). **B**, Apparent affinity of mAbs binding to the mammalian cell grown vaccine (Flucelvax), insect cell grown vaccine (Flublok), and egg-grown vaccine (Fluzone). Each line connects the same mAb binding a different vaccine (n=11 mAbs). **C**, Egg-specific mAb binding to Newcastle Disease Virus grown in eggs and egg-grown A/California/7/2009 H1N1 virus (n=10). Each line connects the same mAb. **D**, Egg-specific mAbs (n=10) binding to the 2020/2021 Fluarix vaccine (egg-grown), measles/mumps/rubella vaccine (MMR; chicken cell line grown), rabavert (Rabies vaccine; chicken cell grown), Japanese Encephalitis Virus Vaccine (JEV, Ixiaro, Vero-cell grown), and pneumovax-23 vaccine (polysaccharides from bacteria). Data in **D** are mean ± S.D. Data in **B** were analyzed using a non-parametric Friedman Test. Data in **C** were analyzed using a two-tailed Wilcoxon matched-pairs signed rank test.

### Sulfated LacNAc is the egg-derived antigen

Egg allergies are typically caused by antibody responses against ovalbumin and ovomucoid (18), which are present in both the egg white and allantoic fluid (19, 20). However, the egg-specific mAbs did not bind to ovalbumin or ovomucoid purified from egg whites (data not shown), further indicating these mAbs were likely not specific to known egg allergens. As the antigen did not seem to be protein in nature, we next tested whether egg-binding mAbs were binding an egg-associated glycan. To test this, we degylcosylated the 2020 Fluarix QIV with a degylcosylating enzyme that removes N-linked glycans. MAbs had reduced binding to deglycosylated vaccine relative to untreated vaccine (Figure 3A). To investigate the particular glycan these mAbs were binding, we tested two mAbs (029-09 3A04 and 034-10 4G02) for binding to a glycan microarray that included 585 distinct glycans (Table S2). Both mAbs specifically bound to two sulfated LacNAc antigens, (6S)(4S)Glaβ1-4GlcNAcβ and (4S)Glaβ1-4GlcNAcβ (Figure 3 B and C). Treatment of purified egg-grown A/Hong Kong/485197/2014 H3N2 with a sulfate ester sulfatase significantly reduced egg-specific mAb binding (Figure 3D). Moreover, both mAbs only bound to LacNAc with a sulfate group on the hydroxyl group of 4C’ of galactose, and not the hydroxyl group on 6C’ of the galactose of LacNAc only, or an unsulfated LacNAc (Figure 3E and F; Table S3). Together, these data reveal that B cells induced by influenza viruses grown in eggs are binding a sulfated LacNAc.

**Figure 3:**
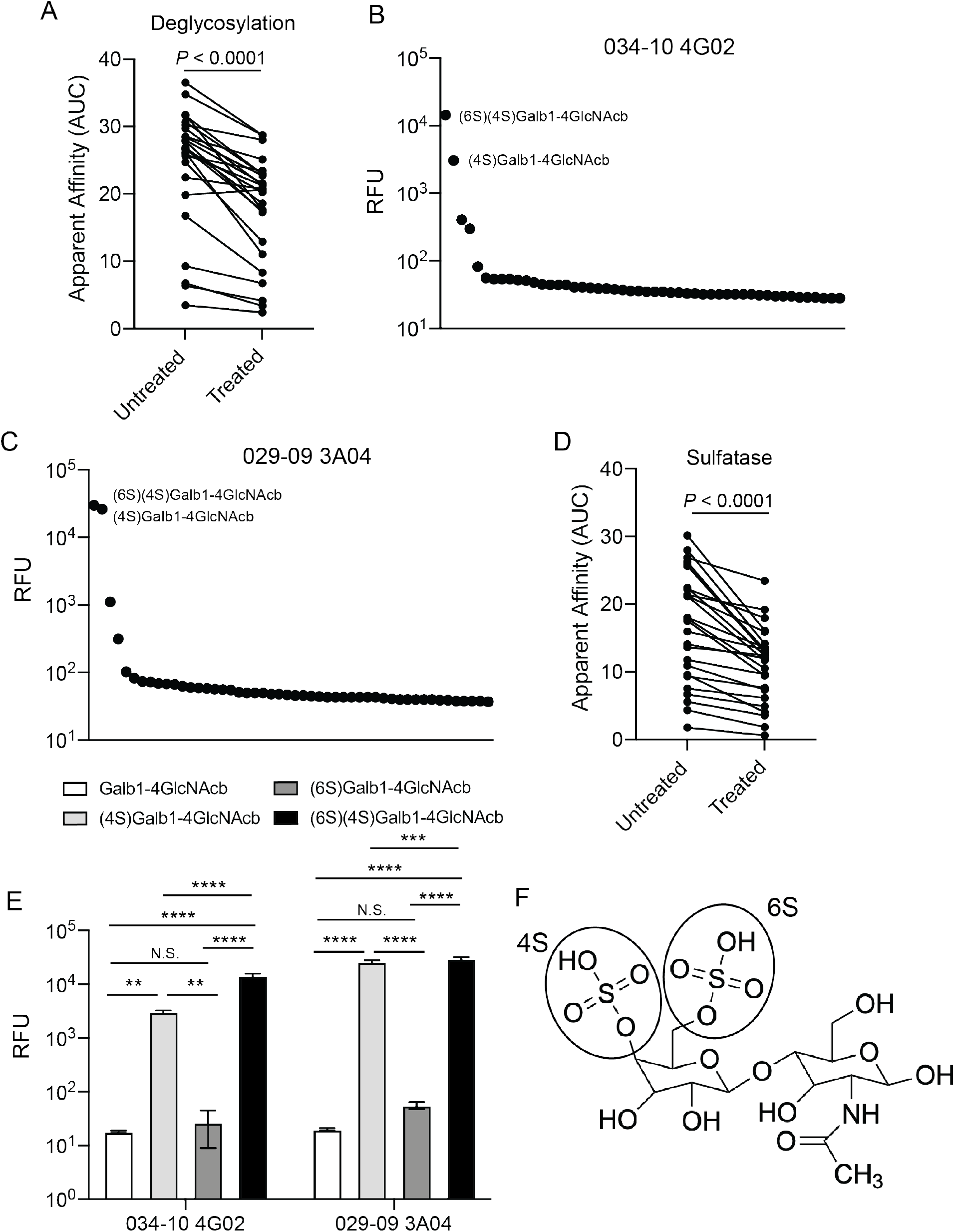
Egg-specific mAbs are binding to a sulfated LacNAc. **A**, Egg-specific mAb (n=24) binding to deglycosylated or untreated 2020 Fluarix QIV. **B** and **C**, 034-10 4G02 (**B**) and 029-09 3A04 (**C**) were tested for binding to glycans on a microarray. Data represent the top 50 glycan hits. **D**, Egg-specific mAb (n=26) binding to sulfatase treated or untreated A/Hong Kong/485197/2014 H3N2. **E**, 034-10 4G02 and 029-09 3A04 binding to non-sulfated and sulfated LacNAc glycans in glycan microarray. **F**, Structure of (6S)(4S)Galb1-4GlcNacb (LacNAc). Data in **B**, **C**, and **E** are averaged RFU of 4 replicates. Each line in **A** and **D** connects the same mAb. Data in **E** are mean ± S.D. Data in **A** and **D** were analyzed using a two-tailed Wilcoxon matched-pairs signed rank test and data in **E** were analyzed using an ordinary two-way ANOVA.

### Egg binding antibodies utilize a restricted repertoire and resemble natural antibodies

Of the egg-binding mAbs, we identified strong repertoire biases on the usage of particular heavy and light chain variable genes, with the vast majority of mAbs using VH3-07 and VL1-44 (Figure 4A and B). However, there was a lot of diversity in the H-CDR3 sequences, with no consensus on DH gene usage (Figure S2A). H-CDR3s and light chain CDR3s (L-CDR3) of egg-binding mAbs preferentially used JH4 and JL3, suggesting there was some selection for certain J genes (Figure S2B and C). Moreover, H-CDR3s of egg-binding mAbs were significantly shorter than those of vaccine-specific mAbs, whereas the L-CDR3 were significantly longer than vaccine-specific mAbs (Figure 4C). Concordantly, H-CDR3s of egg-binding mAbs had fewer non-templated DNA insertions, or N-nucleotides, relative to vaccinespecific mAbs (Figure 4D; Figure S2D).

**Figure 4:**
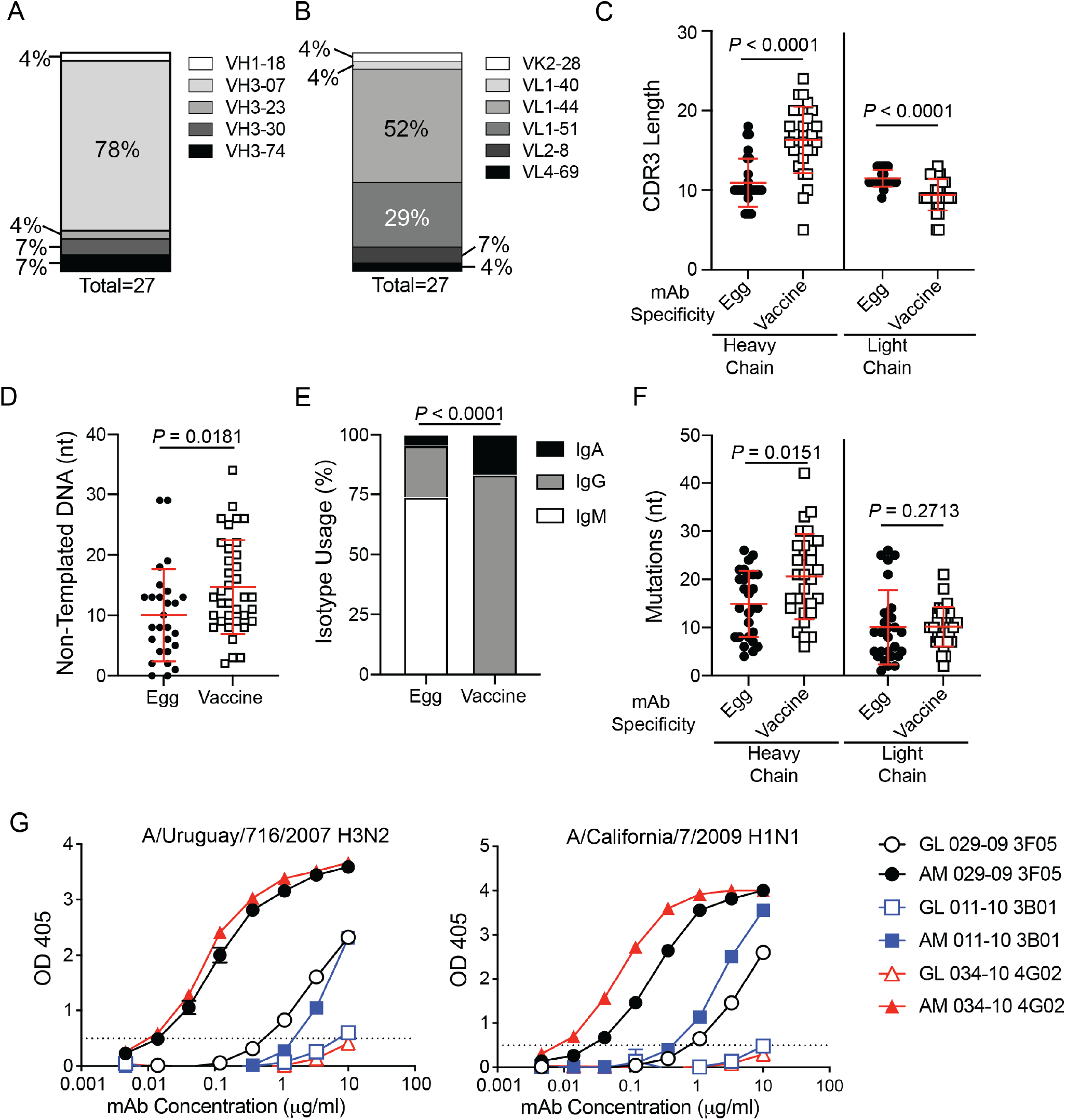
Egg-binding mAbs resemble natural antibodies. **A** and **B**, VH (**A**) and VK/VL (**B**) gene usage of egg-binding mAbs. **C** and **D**, heavy chain and light chain CDR3 lengths (**C**) and non-templated DNA (N-nucleotides; **D**) of egg-binding and vaccine-specific mAb heavy chains. **E** and **F**, isotype usage (**E**) and somatic hypermutations (nucleotide mutations; nt; **F**) of egg-binding and vaccine-specific mAbs. **G**, affinity-matured (AM) 029-09 3F05, 011-10 3B01, and 034-10 4G02 were reverted back to germline (GL) and were tested for binding to egg-grown A/Uruguay/716/2007 H3N2 and A/California/7/2009 H1N1 relative to their affinity-matured (AM) counterparts. Data in **C**, **D**, **F**, and **G** are mean ± S.D. Data in **C**, **D**, and **F** were analyzed using two-tailed Mann-Whitney tests. Data in **E** were analyzed using chi-squared test.

Analysis of clonal expansions revealed that the light chain of egg-binding mAbs across individual mAbs were highly clonal, with one light chain clone occupying over one-third of all egg-binding mAbs (Figure S2E; Table S2 and S5). Additionally, we identified 4 distinct clonal expansions, accounting for one-third of all antibodies identified (Figure S2E). However, these paired heavy and light chain clones were private, with individual clones only identified in one subject each (Table S2). PBs induced by vaccination derive from memory B cells and therefore are usually class-switched to IgG and highly mutated (17). However, egg-binding mAbs were largely IgM and had fewer mutations in the heavy chain relative to vaccine-specific mAbs (Figure 4E and F). In combination, egg-binding mAbs resemble natural antibodies produced by innate-B cells, as they express simple and short H-CDR3s, do not commonly class-switch, and have fewer mutations than vaccine-specific antibodies (21, 22). In addition, natural antibodies commonly target glycans (23) and are polyreactive (24, 25). However, egg-binding mAbs were not enriched for polyreactivity relative to vaccine-induced antibodies (Figure S2F). Despite having fewer mutations than vaccine-induced antibodies, germline (GL) reverted egg-binding mAbs had reduced binding affinity for influenza viruses grown in eggs relative to the affinity-matured (AM) mAbs generated from PBs (Figure 4G). Furthermore, we identified a clonal expansion from one subject over two influenza virus vaccine seasons (2010 TIV and 2011 TIV), with the mAbs from 2011 having higher affinity for influenza virus strains relative to the mAb from 2010 (Figure S2G-I). With the highly restricted VH/VL repertoire, short H-CDR3 sequences and reduced N-nucleotide additions, lack of class-switching, and fewer mutations, mAbs targeting sulfated LacNAc resemble natural antibodies produced by innate-like B cells (26).

### Influenza virus vaccines commonly induce PBs with repertoire features of egg-binding mAbs

We next addressed whether antibodies with repertoire features of egg-binding antibodies are commonly induced after influenza virus vaccination. To dissect this question, we utilized a dataset of 7,777 B cell receptor sequences from influenza virus vaccine induced PBs from subjects that received the egg-grown 2016-2017 Fluzone QIV (27). From this dataset, we identified that 2% (175 total) of all IgG^+^ PBs expressed VH3-7 with a H-CDR3 length of equal to or less than 12 amino acids that was paired with VL1 – 44 or VL1-51 (potential egg-binding mAbs; Figure 5A). Notably, 13 out of 17 total subjects had PBs with these repertoire features, occupying 0.2-17.4% of the PB response per subject (Figure 5B). Notably, this dataset was specifically generated from IgG^+^ PBs. As we identified most egg-binding mAbs were IgM (Figure 4E), the true number of potential egg-specific PBs induced within these subjects may be substantially higher. Of the 175 heavy and light chain pairings identified, we discovered 6 public clonotypes (Figure 5C; Table S5), which comprised 66.9% of the total response (117/175 paired sequences). Strikingly, 3 of the public clones were shared between our study and the dataset (Figure 5C; Table S5), suggesting the PBs induced in subjects in the Forgacs et al. study are specific to the egg glycan. Moreover, we identified that egg-binding mAbs from our study shared at least a heavy chain or light chain clone with the potential egg-binding PBs from the IgG^+^ PB dataset (Figure 5D; Table S5). Together, these data suggest that PBs targeting an egg-associated glycan are commonly induced by influenza virus vaccines grown in eggs.

**Figure 5:**
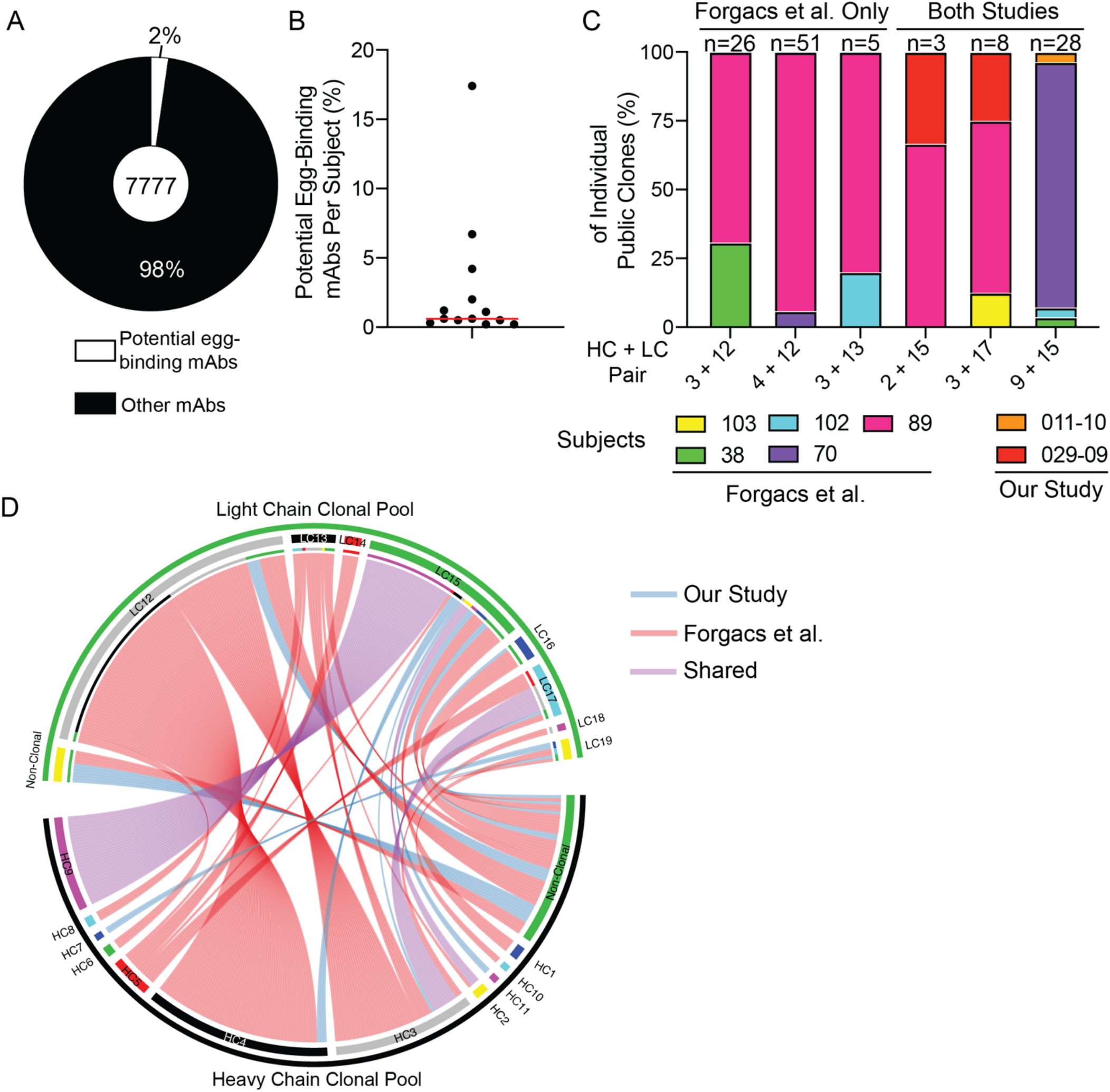
PBs with repertoire features of egg-binding antibodies are common after vaccination. B cell receptor sequences from 7,777 IgG^+^ PBs induced by influenza virus vaccination were analyzed for repertoire features of egg-binding mAbs (VH3-7 with H-CDR3 of ≤ 12 amino acids paired with VL1-44 or VL1-51). **A** and **B**, proportion of sequences with egg-binding mAb repertoire features out of all sequences (**A**) and by subject (**B**). Red line in **B** represents the median. **C**, subject makeup (%) of public clones, including public clones specific to Forgacs et al. and shared across studies. The number above each column represents the number of clonal members per clone. **D,** circos plot of heavy and light chain clones distinct to each study (our study: blue; Forgacs et al. red) or shared across studies (purple).

## Discussion

In this report, we identified that egg-grown viruses possess an egg-associated glycan that is immunogenic in humans. Egg-specific mAbs from PBs had evidence of prior affinity-maturation and likely derived from memory B cells that were primed by earlier vaccination with egg-grown vaccines. Moreover, most subjects within the 2010 TIV cohort had previously been vaccinated with the 2009 MIV, suggesting prior exposure to the egg antigen from the 2009 MIV generated memory B cells against this antigen. Despite some subjects mounting a response against the egg antigen, the same subjects seroconverted against the H1N1 component of the vaccine to similar levels as subjects that did not mount a response against the egg-associated antigen. Moreover, 13 out of 17 subjects from Forgacs et al. had detectable PBs with repertoire features of egg-binding mAbs. Notably, this study specifically recruited subjects that had not been vaccinated in the prior 3 seasons (27). Prior research has indicated that pre-existing serum antibodies against the strains included in the vaccine inversely correlate with the induction of PBs by vaccination (28). Therefore, subjects that mounted an anti-egg response perhaps had more B cell activation relative to subjects that did not and as a result has similar anti-HA antibody responses.

Despite no differences in serum antibodies against the H1N1 component of the vaccine, B cells mounted against the egg antigen may compete within germinal centers with B cells targeting protective epitopes of HA and NA, which could perturb the generation of plasma cells and memory B cells against protective epitopes. In accordance, we identified the same egg-specific clone was recalled over multiple vaccine years, suggesting subjects can repeatedly recall memory B cells against the egg-derived antigen upon repeated vaccination and further indicating egg-specific antibodies are fixed in the memory B cell repertoire. Therefore, our study suggests that repeatedly vaccinated subjects can preferentially recall memory B cells targeting irrelevant antigens associated with egg-based vaccine production, which could come at the cost of affinity-maturation and generation of plasma cells and memory B cells specific for protective epitopes that provide long-lived protection against influenza viruses. However, the precise role of egg-specific B cells competing with virus-specific B cells remains to be determined.

The egg-binding mAbs demonstrated features of natural antibodies produced by innate-like B cells, including their glycan specificity and repertoire features. Natural antibodies are typically raised against self-antigens and evolutionarily conserved antigens such as lipids and glycans (29). In humans, natural antibodies are largely elicited against glycans, including the blood group A and B antigens and xenoglycans from other mammals, including the “a-gal” epitope of galactose-α-1,3-galactose expressed by most non-primate mammals (30, 31). Our studies reveals that the egg-binding antibodies were specifically targeting a secondary sulfate structure of LacNAc, a common glycan found across all life forms. Although the egg-binding mAbs identified in this study share key characteristics of natural antibodies, the precise cellular origins of these antibodies require further analysis.

LacNAcs are a critical component of glycosaminoglycans (GAG), including keratan sulfate and the Lewis blood group determinants (32, 33). LacNAcs are also the main ligand for galectins and mediate a variety of cellular functions including cell adhesion, migration, proliferation, and apoptosis (34–36). However, most mammals do not commonly sulfonate the galactose 4C’ of LacNAc, and instead commonly sulfate the 6C’ of galactose of LacNAc and 6C’ of N-acetylglucosamine of LacNAC (37). Therefore, humans may mount a response specifically against the sulfated 4C’ of LacNAc as it is not normally sulfated in humans. However, inflammation is associated with aberrant glycosylation patterns, including during cancer and autoimmunity (38). Moreover, antibodies against a sulfated 4C’ LacNAc, the same antigen identified in this study, were elevated in subjects with systemic sclerosis and was associated with a higher prevalence of pulmonary hypertension (39). Despite these findings, the precise role and function of anti-sulfated-LacNAc antibodies in the development and severity of systemic sclerosis remain unknown. Furthermore, it is unknown if the antibodies induced by egg-grown influenza virus vaccines and those observed during systemic sclerosis share similar repertoire features and could be derived from the same B cell precursors. Lastly, it remains unknown how the 4C’ sulfate LacNAc is conjugated to influenza viruses grown in eggs, as a study of the glycosylation patterns of HA and NA isolated from egg grown viruses did not have this glycan (40). In summary, this study identifies that antibodies with features of natural antibodies can be induced against a sulfated LacNAc glycan present in egg-grown vaccines.

## Methods

### Monoclonal antibody production and sequence analysis

Monoclonal antibodies were generated as previously described (15, 41, 42). Peripheral blood was obtained from each subject approximately 7 days after vaccination or infection. Lymphocytes were isolated and enriched for B cells using RosetteSep. PBs (CD3^-^CD19^+^CD27^hi^CD38^hi^) were single-cell sorted into 96-well plates. Immunoglobulin heavy and light chain genes were amplified by reverse transcriptase polymerase chain reaction (RT-PCR), sequenced, cloned into human IgG1, human kappa chain, or human lambda expression vectors, and co-transfected into HEK293T cells. Secreted mAbs were purified from the supernatant using protein A agarose beads. B cell clones were determined by aligning all the V(D)J sequences sharing identical progenitor sequences, as predicted by IgBLAST using our in-house software, Vgenes. For germline mAbs, germline sequences were synthesized (IdT) and cloned into antibody expression vectors, as described above.

### Antibody sequences and clonal analyses

Previously published IgG^+^ PB sequences (27) were downloaded from NCBI GenBank (KEOV00000000 and KEOU00000000). V(D)J gene usage from our study and Forgacs et al. were analyzed using IgBlast and clones were determined using our in-house software, Vgenes, based on germline sequences. For identification of egg-like mAbs from Forgacs et al. we selected B cell clones that specifically used VH3-7 with a H-CDR3 of 12 or fewer amino acids that was paired with VL1-44 or VL1-51. MAb sequence alignments were made using ClustalOmega (EMBL-EBI). Non-templated nucleotide insertions at the V-D and D-J junctions of the heavy-chain gene were identified using partis v0.15.0, a Hidden Markov Modelbased tool for annotating B cell receptor sequences (43). Custom code was used for processing the output (available at https://github.com/cobeylab/egg_antibodies). Visualization of clones in Figure 5D was performed in R using circlize v0.4.12 (44).

### Viruses, proteins, and vaccines

Influenza viruses used in all assays were grown in-house in specific pathogen free (SPF) eggs, harvested, purified, and titered. Allantoic fluid was harvested from both infected and uninfected eggs. For MDCK cell grown virus, A/California/7/2009 H1N1 was grown in MDCK-SIAT1 cells, concentrated, and chemically inactivated with beta-propiolactone. Newcastle disease virus was grown in eggs and allantoic fluid was harvested and subsequently inactivated with beta-propiolactone, purified, and quantified. Vaccines used for mAb binding assays are outlined in Table S6. Recombinant HA (rHA) from A/California/7/2009 was expressed in HEK293T cells.

### Antigen-Specific ELISA

High protein-binding microtiter plates (Costar) were coated with 8 hemagglutination units (HAU) of virus or allantoic fluid diluted 1:500 in carbonate buffer overnight at 4°C. Plates were coated with recombinant HA from A/California/7/2009 at 1 μg/ml in PBS overnight at 4°C. For testing egg-binding mAb binding to various vaccines, influenza virus vaccines were diluted to 5 μg/ml, rabavert was diluted to 0.05 UI/ml, MMR was diluted 1:100, Ixiaro (JEV) was diluted to 0.05 antigen units per ml, and Pneumovax 23 was diluted to 5 μg/ml. All tested vaccines were diluted in PBS and coated overnight at 4°C. NDV was diluted to 5 μg/ml in carbonate buffer and plates were coated overnight at 4°C.

Plates were washed the next morning with PBS 0.05% Tween and blocked with PBS containing 20% fetal bovine serum (FBS) for 1 hour at 37°C. MAbs were then serially diluted 3-fold starting at 10 μg/ml and incubated for 1.5 hour at 37°C. For serum ELISAs, serum was diluted 1:50 and further diluted 2-fold. Horseradish peroxidase (HRP)-conjugated goat anti-human IgG antibody diluted 1:1000 (Jackson Immuno Research) was used to detect binding of mAbs and serum antibodies, and plates were subsequently developed with Super Aquablue ELISA substrate (eBiosciences). Absorbance was measured at 405 nm on a microplate spectrophotometer (BioRad). To standardize the assays, control antibodies with known binding characteristics were included on each plate, and the plates were developed when the absorbance of the control reached 3.0 OD units. For the other vaccines used in Figure 2D, anti-sera against the various vaccines were used to confirm antigenicity of vaccines. Polyreactivity was determined using a polyreactive ELISA protocol, as previously described (13). Briefly, mAbs were tested for binding to 6 antigens (cardiolipin, dsDNA, flagellin, insulin, KLH, and LPS) starting at 1 μg/ml for 1.5 hours at 37°C. HRP-conjugated goat anti-human IgG antibody (Jackson Immuno Research) diluted 1:2000 in PBS/0.05% Tween/0.1mM EDTA was used to detect binding of mAbs, and plates were subsequently developed with Super Aquablue ELISA substrate (eBiosciences). All mAbs or serum samples were tested in duplicate and all assays were performed 2-3 times. To determine mAb affinity, a non-linear regression was performed on background subtracted ODs and area under the curve (AUC) values were reported. Serum samples used in Figure 1h and Figure S1d-f are listed in Table S7.

### Hemagglutination inhibition assays

For serum HAI assays, 1 part serum was treated with 3 parts Receptor Destroying Enzyme II (Seiken, Hardy Diagnostics) for 18 hours at 37°C, followed by 30 minutes at 56°C. Serum was further diluted to 1:10 with PBS and serially diluted 2-fold in PBS in duplicate in a 96-well round-bottom plate. Serially diluted serum was mixed with an equal volume of A/California/7/2009 virus (4 HAU/25 μl), and subsequently incubated at room temperature for 1 hour. 50 μl of 0.5% turkey red blood cells (Lampire Biological) were added to each well and incubated for 45 minutes at room temperature. HAI titers were determined based on the final dilution of serum for which hemagglutination inhibition was observed. All experiments were performed in duplicate twice. The fold change in HAI serum titers of post-vaccination samples relative to pre-vaccination samples are shown in Figure S1F.

### Virus deglycosylation and sulfatase treatment

To deglycosylate the vaccine, 25 ug of the 2020 QIV was denatured for 10 minutes at 75°C and treated with the Protein Deglycosylation Mix II (New England Biolabs) for 30 minutes at 25°C and 1 hour at 37°C. For sulfatase treatment, A/Hong Kong/485197/2014 H3N2 virus was diluted to 160 HAU in sodium acetate (pH 5.0) and treated with 20 units/ml of sulfatase from abalone entrails (Sigma-Aldrich) for 1 hour at 37°C. As a control, equal quantities were diluted and incubated but did not receive the Deglycosylation Mix II or sulfatase enzymes and are referred to as untreated. After treatment, preparations underwent buffer exchange with PBS to remove freed glycans and sulfate groups. ELISA plates were coated at 1 μg/ml for the 2020 QIV or 8 HAU for the virus.

### Glycan microarray

MAbs 029-09 3A04 and 034-10 4G02 were sent to the Protein-Glycan Interaction Resource of the Center for Functional Glycomics at the Beth Israel Deaconess Medical Center, Harvard Medical School. Printed arrays consist of 585 glycans in replicate of 6. All glycans used in the microarray are listed in Table S3 and S4. MAbs were diluted to 50 μg/ml and run on the array. The highest and lowest replicates from each set of 6 replicates were removed and the average ± S.D. was calculated from the middle 4 replicates. Structure of 6S,4S-LacNAc was made using ChemDraw JS (PerkinElmer).

### Statistics

All statistical analysis was performed using Prism software (Graphpad Version 9.0). *P* values less than or equal to 0.05 were considered significant. * *P* ≤ 0.05, ** *P* ≤ 0.01, *** *P* ≤ 0.001, **** *P* < 0.0001. Specific number of mAbs shown in each Figure are listed in the corresponding figure legends.

### Subjects, cohorts, and study approval

Written informed consent was received from participants prior to inclusion in the study. All studies were performed with the approval of the University of Chicago and Emory University institutional review boards. Subjects were recruited to receive the Sanofi Pasteur 2009 pandemic H1N1 MIV or the 2010 Novartis Fluvirin TIV. Subjects labeled with SFV were recruited and vaccinated at Emory University, as previously described (45). All other subjects were recruited and vaccinated at the University of Chicago. Subject demographics are detailed in Table S1.

## Author contributions

J.J.G. designed the study, characterized mAbs, analyzed the data, and wrote the manuscript. H.A.U. characterized mAbs. C.H. assisted with the vaccine comparison study. L.L. assisted with data analysis. N.-Y.Z. grew and purified influenza viruses. W.S. provided inactivated Newcastle Disease Virus samples. M.C.V. performed non-templated DNA sequence analyses. S.Z. and S.H. provided MDCK cell grown virus. M.H. performed mAb cloning. S.C. and P.P. provided critical feedback and discussion on the manuscript. P.C.W. supervised the work and edited the manuscript. All authors reviewed and edited the manuscript.

## Acknowledgments

We are thankful to all subjects who participated in this study. We thank Sarah Andrews for initiating and processing samples from the 2009 MIV, 2010 TIV, and 2011 TIV cohort studies, and Jens Wrammert and Rafi Ahmed for providing samples from the 2009 MIV cohort. We thank Jamie Heimburg-Molinaro and Richard Cummings for assistance with the glycan microarray. This project was funded in part by the National Institute of Allergy and Infectious Diseases; National Institutes of Health grant numbers U19AI082724 (P.C.W.), U19AI109946 (P.C.W.), U19AI057266 (P.C.W.), P01 AI097092 (P.P.), R01AI145870-01 (P.P.), and the NIAID Centers of Excellence for Influenza Research and Surveillance (CEIRS) grant number HHSN272201400005C (P.C.W. and S.E.H.), and HHSN272201400008C (P.P.). This work was also partially supported by the National Institute of Allergy and Infectious Disease (NIAID) Collaborative Influenza Vaccine Innovation Centers (CIVIC; 75N93019C00051, P.P., P.C.W.). We thank the Protein-Glycan Interaction Resource of the CFG and the National Center for Functional Glycomics (NCFG) at Beth Israel Deaconess Medical Center, Harvard Medical School (R24 GM137763).

