## Supplemental Figures for "An egg-derived sulfated N-Acetyllactosamine glycan is an antigenic decoy of influenza virus vaccines"

### Figure S1

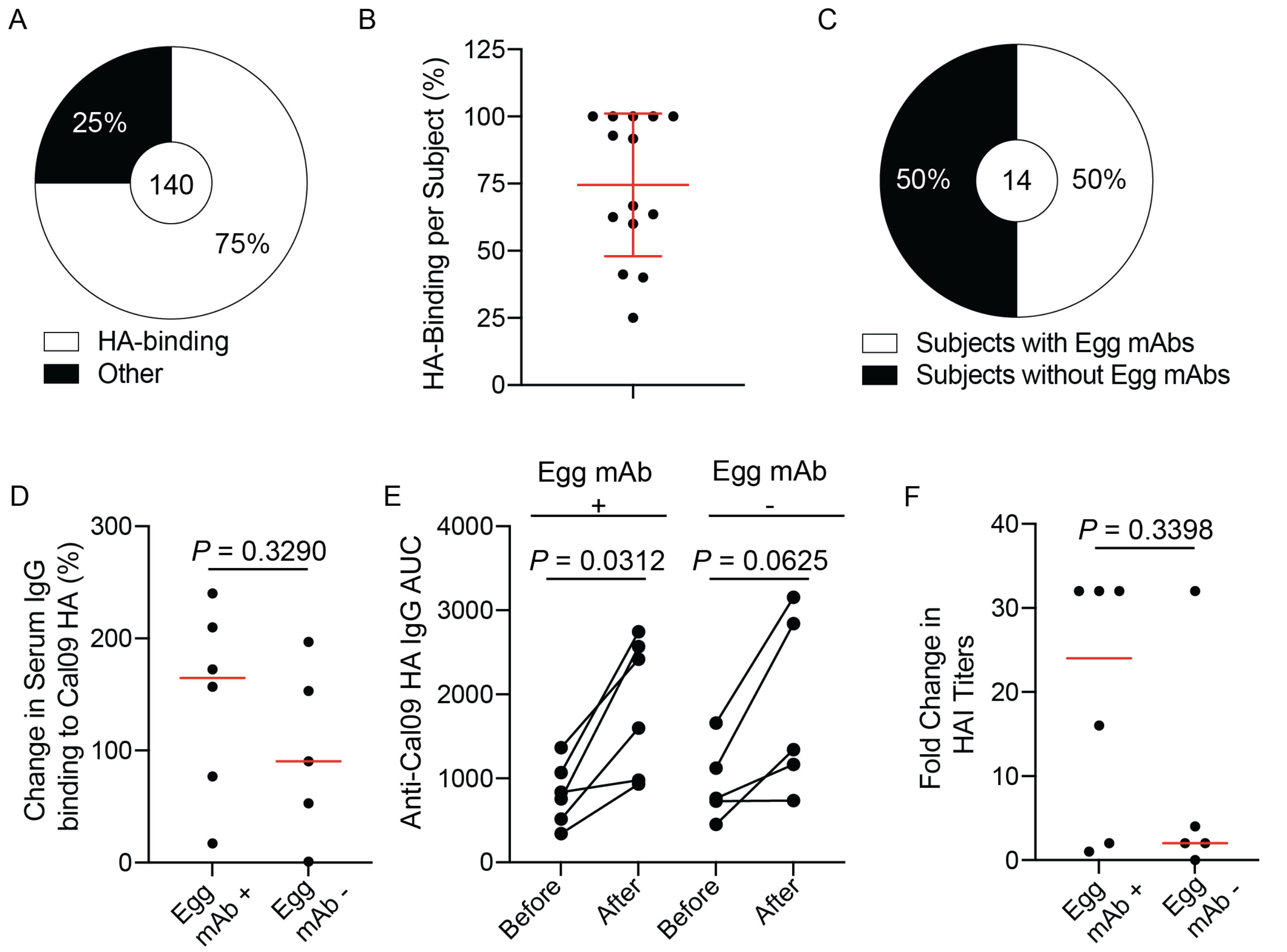


**Fig. S1: MAb binding to HA and egg-derived antigen. A**, Proportion of total mAbs binding to HA after vaccination with an egg-grown vaccine. Number in the center indicates number of mAbs tested. **B**, Proportion of HA-binding mAbs per subjects. Data are mean ± S.D. **C**, Proportion of subjects with detectable egg mAbs. Number in the center indicates number of subjects. **D** and **E**, Serum was isolated from subjects with or without isolated egg-binding mAbs before and 14-21 days after vaccination. **D**, Fold-change (represented as a percentage) in serum IgG binding to rHA from A/California/7/2009 H1N1. **E**, Serum IgG binding to A/California/7/2009 rHA before and after vaccination. Lines connect responses from individual subjects. **F**, Fold change in serum HAI titers against A/California/7/2009 after vaccination. Lines in **D** and **F** are the median. Data in **D** and **F** were analyzed using a two-tailed Mann-Whitney test. Data in **E** were analyzed using a two-tailed Wilcoxon matched-pairs signed rank test.

Figure S2

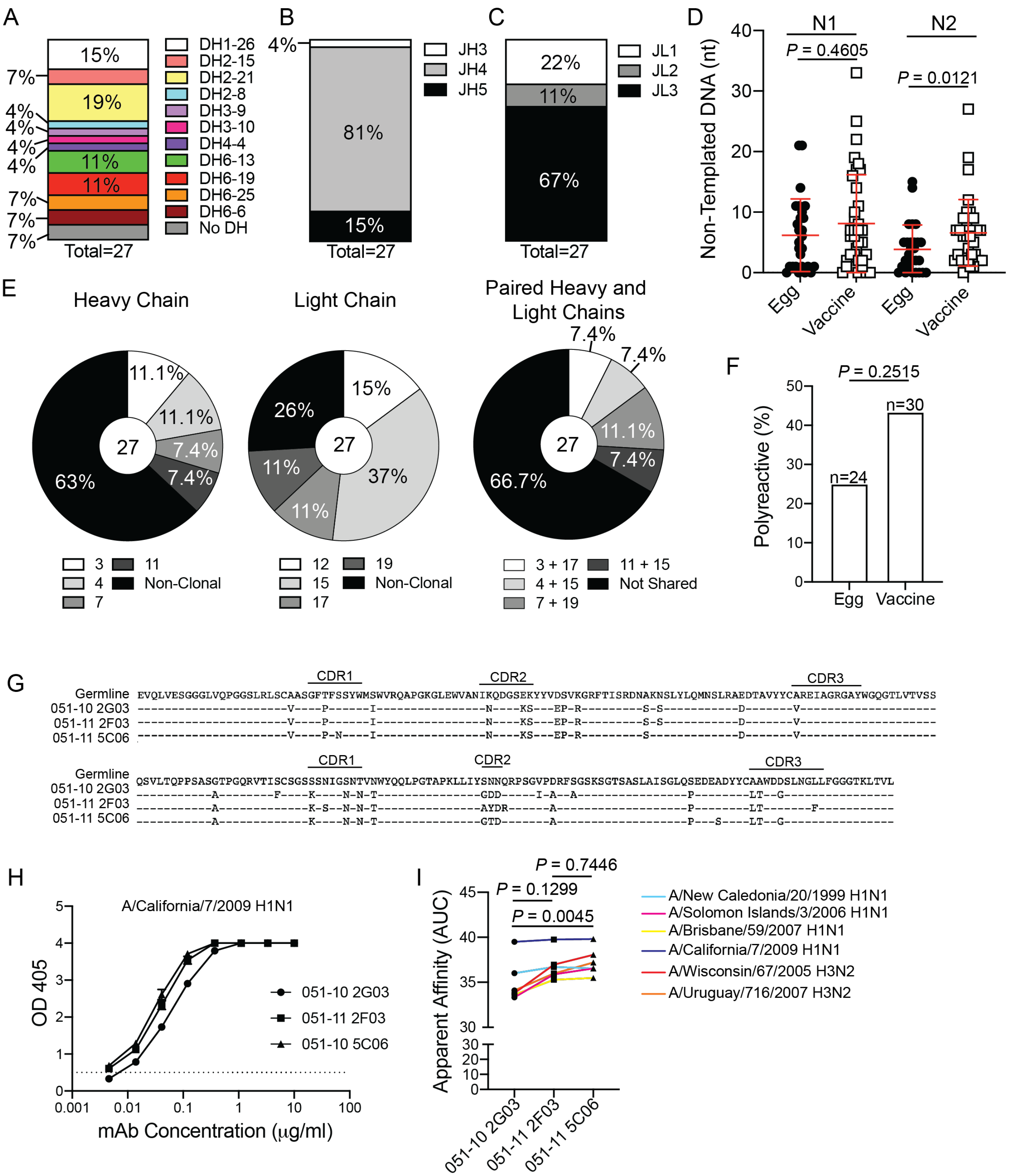


**Fig. S2: Additional repertoire and clonal expansions of egg-binding mAbs. A-C**, DH (**A**), JH (**B**), and JK/JL (**C**) gene usage by egg-binding mAbs. **D**, N1 (V-D) and N2 (D-J) non-templated nucleotide insertions of heavy chain CDR3s of egg- and vaccine-binding mAbs. **E**, clonal relatedness of heavy chain and light chains of egg-binding mAbs. **F**, polyreacitivity of egg-binding mAbs relative to vaccine-specific mAbs. **G-I**, Expansion of an egg-binding clone across multiple years of vaccination. **G**, Alignment of heavy chain and light chain sequences. **H** and **I**, Binding curves of mAbs binding to A/California/7/2009 (**H**) and apparent affinity (AUC; **I**) of mAbs binding to egg-grown influenza viruses. Data in **D** and **H** are mean ± S.D. Data in **D** were analyzed using two-tailed Mann-Whitney tests and data in **F** were analyzed using chi-squared test. Data in **I** were analyzed using non-parametric paired Friedman tests.

| Egg Reactivity | Subject | Vaccine Year/ Formulation | Vaccine/ Manufacturer | Known Prior Vaccination(s) | Age | Sex |
|---|---|---|---|---|---|---|
| Egg mAb Positive | 029-09 | 2009 MIV | Sanofi Pasteur | Unknown | 24 | M |
| | 008-10 | 2010 TIV | Novartis Fluvirin | 2009 TIV, 2009 MIV | 26 | F |
| | 011-10 | 2010 TIV | Novartis Fluvirin | Unknown | 30 | M |
| | 017-10 | 2010 TIV | Novartis Fluvirin | 2009 MIV | 24 | M |
| | 019-10 | 2010 TIV | Novartis Fluvirin | 2009 MIV | 23 | F |
| | 034-10 | 2010 TIV | Novartis Fluvirin | Unknown | 40 | F |
| | 051-10 | 2010 TIV | Novartis Fluvirin | 2009 MIV | 43 | M |
| Egg mAb Negative | 030-09 | 2009 MIV | Sanofi Pasteur | Unknown | 31 | F |
| | SFV018 | 2009 MIV | Sanofi Pasteur | 2009 TIV | 58 | F |
| | SFV019 | 2009 MIV | Sanofi Pasteur | 2009 TIV | 48 | F |
| | SFV020 | 2009 MIV | Sanofi Pasteur | 2009 TIV | 64 | F |
| | 014-10 | 2010 TIV | Novartis Fluvirin | 2009 TIV, 2009 MIV | 27 | M |
| | 028-10 | 2010 TIV | Novartis Fluvirin | Unknown | 32 | M |
| | 039-10 | 2010 TIV | Novartis Fluvirin | 2009 TIV, 2009 MIV | 25 | M |

**Table S1: Study cohort demographics.**

| mAb | VH | DH | JH | VK/VL | JK/JL | H-CDR3 Length | HC Mutations | Original Isotype | Clone (HC/LC) |
|---|---|---|---|---|---|---|---|---|---|
| 029-09 3A01 | VH3-30 | DH2-21 | JH4 | VL2-8 | JL3 | 15 | 25 | IgM | 0/0 |
| 029-09 3A04 | VH3-7 | DH6-19 | JH4 | VL1-51 | JL3 | 10 | 14 | IgG | 7/19 |
| 029-09 3C04 | VH1-18 | DH6-19 | JH4 | VL1-40 | JL3 | 17 | 18 | IgM | 0/0 |
| 029-09 3C06 | VH3-7 | DH6-19 | JH4 | VL1-51 | JL3 | 10 | 14 | IgM | 7/19 |
| 029-09 3D03 | VH3-7 | DH6-6 | JH4 | VL1-51 | JL2 | 10 | 21 | IgM | 3/17 |
| 029-09 3D04 | VH3-7 | DH6-25 | JH5 | VL1-51 | JL2 | 10 | 21 | IgM | 0/17 |
| 029-09 3D06 | VH3-74 | DH3-10 | JH4 | VL2-8 | JL3 | 9 | 21 | IgM | 0/0 |
| 029-09 3F05 | VH3-7 | DH6-25 | JH4 | VL1-44 | JL3 | 10 | 6 | IgM | 2/15 |
| 029-09 3G05 | VH3-7 | DH1-26 | JH4 | VL1-51 | JL2 | 10 | 17 | IgG | 3/17 |
| 029-09 4E04 | VH3-23 | DH6-6 | JH6 | VK2-28 | JK1 | 18 | 26 | IgM | 0/0 |
| 008-10 6D02 | VH3-7 | DH6-13 | JH4 | VL1-51 | JL3 | 10 | 24 | unknown | 0/19 |
| 011-10 2A01 | VH3-7 | DH2-8 | JH5 | VL1-44 | JL3 | 11 | 5 | IgM | 0/15 |
| 011-10 2F01 | VH3-30 | DH2-21 | JH4 | VL1-51 | JL3 | 12 | 8 | IgM | 0/0 |
| 011-10 2G01 | VH3-7 | DH2-21 | JH5 | VL1-44 | JL3 | 14 | 8 | IgM | 11/15 |
| 011-10 3A03 | VH3-7 | DH2-21 | JH5 | VL1-44 | JL3 | 14 | 8 | IgM | 11/15 |
| 011-10 3B01 | VH3-7 | DH3-9 | JH4 | VL1-44 | JL3 | 7 | 18 | IgM | 0/15 |
| 011-10 3B03 | VH3-7 | None | JH4 | VL1-44 | JL3 | 7 | 4 | IgM | 9/15 |
| 017-10 3B06 | VH3-7 | DH2-15 | JH4 | VL1-44 | JL1 | 10 | 13 | IgG | 0/12 |
| 017-10 3D06 | VH3-74 | None | JH4 | VL4-69 | JL3 | 7 | 20 | IgM | 0/0 |
| 017-10 3E02 | VH3-7 | DH2-21 | JH4 | VL1-44 | JL1 | 10 | 6 | IgM | 0/12 |
| 019-10 4A06 | VH3-7 | DH4-4 | JH4 | VL1-44 | JL1 | 10 | 7 | unknown | 0/12 |
| 019-10 4E01 | VH3-7 | DH2-15 | JH4 | VL1-51 | JL1 | 7 | 9 | IgG | 0/16 |
| 034-10 4G02 | VH3-7 | DH6-13 | JH3 | VL1-44 | JL1 | 10 | 11 | IgM | 0/12 |
| 051-10 2E02 | VH3-7 | DH6-13 | JH4 | VL1-44 | JL3 | 10 | 14 | IgA | 3/15 |
| 051-10 2G03 | VH3-7 | DH1-26 | JH4 | VL1-44 | JL3 | 10 | 24 | IgG | 4/15 |
| 051-11 2F03 | VH3-7 | DH1-26 | JH4 | VL1-44 | JL3 | 10 | 22 | unknown | 4/15 |
| 051-11 5C06 | VH3-7 | DH1-26 | JH4 | VL1-44 | JL3 | 10 | 22 | unknown | 4/15 |

**Table S2: Egg-binding mAb information.** Clonal number of 0 indicates a particular chain was non-clonal.

| Chart ID | Glycan | Average RFU | StDev | %CV |
|---|---|---|---|---|
| **39** | (6S)(4S)Galb1-4GlcNAcb-Sp0 | 14423 | 1539 | 11 |
| **40** | (4S)Galb1-4GlcNAcb-Sp8 | 3055 | 223 | 7 |
| 459 | Neu5Aca2-6Galb1-4GlcNAcb1-6(Neu5Aca2-6Galb1-4GlcNAcb1-2)Mana1-6(GlcNAcb1-4)(Neu5Aca2-6Galb1-4GlcNAcb1-4(Neu5Aca2-6Galb1-4GlcNAcb1-2)Mana1-3)Manb1-4GlcNAcb1-4GlcNAcb-Sp21 | 406 | 47 | 11 |
| 285 | Galb1-3GlcNAcb1-3Galb1-3GlcNAcb-Sp0 | 299 | 28 | 9 |
| 562 | Galb1-4GlcNAcb1-3Galb1-4GlcNAcb1-6(Galb1-4GlcNAcb1-3Galb1-4GlcNAb1-2)Mana1-6(Galb1-4GlcNAcb1-3Galb1-4GlcNAcb1-2Mana1-3)Manb1-4GlcNAcb1-4(Fuca1-6)GlcNAcb-Sp24 | 82 | 7 | 8 |
| 88 | GlcNAcb1-3Galb1-3GalNAca-Sp8 | 56 | 12 | 21 |
| 98 | GalNAcb1-4GlcNAcb-Sp0 | 54 | 62 | 114 |
| 86 | GalNAca1-3(Fuca1-2)Galb1-4GlcNAcb-Sp8 | 54 | 19 | 35 |
| 545 | GlcNAcb1-3Galb1-4GlcNAcb1-6(GlcNAcb1-3Galb1-4GlcNAcb1-2)Mana1-6(GlcNAcb1-3Galb1-4GlcNAcb1-2Man a1-3)Manb1-4GlcNAcb1-4GlcNAc-Sp24 | 54 | 8 | 16 |
| 531 | GlcNAcb1-3Galb1-4GlcNAcb1-2Mana1-6(GlcNAcb1-3Galb1-4GlcNAcb1-2Mana1-3)Manb1-4GlcNAcb1-4GlcNAcb-Sp12 | 52 | 3 | 5 |
| 24 | (3S)Galb1-4(Fuca1-3)(6S)Glc-Sp0 | 51 | 20 | 38 |
| 356 | Fuca1-2Galb1-4(Fuca1-3)GlcNAcb1-2Mana1-6(Fuca1-2Galb1-4(Fuca1-3)GlcNAcb1-2Mana1-3)Manb1-4GlcNAcb1-4GlcNAb-Sp20 | 48 | 30 | 62 |
| 336 | GlcNAca1-4Galb1-4GlcNAcb1-3Galb1-4(Fuca1-3)GlcNAcb1-3Galb1-4(Fuca1-3)GlcNAcb-Sp0 | 45 | 2 | 5 |
| 22 | 6S(3S)Galb1-4(6S)GlcNAcb-Sp0 | 44 | 14 | 31 |
| 534 | Fuca1-2Galb1-4GlcNAcb1-3Galb1-4GlcNAcb1-2Mana1-6(Fuca1-2Galb1-4GlcNAcb1-3Galb1-4GlcNAcb1-2Mana1-3)Manb1-4GlcNAcb1-4GlcNAcb-Sp24 | 44 | 2 | 5 |
| 355 | Fuca1-2Galb1-4GlcNAcb1-2Mana1-6(Fuca1-2Galb1-4GlcNAcb1-2Mana1-3)Manb1-4GlcNAcb1-4GlcNAcb-Sp20 | 44 | 15 | 35 |
| 502 | Galb1-3(6S)GlcNAcb-Sp8 | 41 | 4 | 9 |
| 97 | GalNAcb1-4(Fuca1-3)GlcNAcb-Sp0 | 41 | 11 | 27 |
| 536 | GlcNAcb1-3Galb1-4GlcNAcb1-3Galb1-4GlcNAcb1-2Mana1-6(GlcNAcb1-3Galb1-4GlcNAcb1-3Galb1-4GlcNAcb1-2Mana1-3)Manb1-4GlcNAcb1-4GlcNAcb-Sp25 | 40 | 4 | 11 |
| 582 | Neu5Aca2-3Galb1-4GlcNAcb1-3Galb1-4GlcNAcb1-3Galb1-4GlcNAcb1-2Mana1-6(Neu5Aca2-3Galb1-4GlcNAcb1-3Galb1-4GlcNAcb1-3Galb1-4GlcNAcb1-2Mana1-3)Manb1-4GlcNAcb1-4GlcNAcb-Sp12 | 39 | 2 | 4 |
| 465 | Fuca1-2Galb1-4(Fuca1-3)GlcNAcb1-2Mana1-6(Fuca1-2Galb1-4(Fuca1-3)GlcNAcb1-2Mana1-3)Manb1-4GlcNAcb1-4(Fuca1-6)GlcNAcb-Sp24 | 39 | 3 | 6 |
| 389 | Gala1-3Galb1-3(Fuca1-4)GlcNAcb1-2Mana1-6(Gala1-3Galb1-3(Fuca1-4)GlcNAcb1-2Mana1-3)Manb1-4GlcNAcb1-4GlcNAc-Sp19 | 38 | 8 | 21 |
| 262 | Neu5Aca2-6GalNAcb1-4GlcNAcb-Sp0 | 37 | 25 | 69 |
| 354 | Fuca1-2Galb1-3GlcNAcb1-2Mana1-6(Fuca1-2Galb1-3GlcNAcb1-2Mana1-3)Manb1-4GlcNAcb1-4GlcNAcb-Sp20 | 36 | 5 | 13 |
| 217 | (3S)Galb1-4(Fuca1-3)(6S)GlcNAcb-Sp8 | 36 | 2 | 7 |
| 557 | Galb1-3GlcNAcb1-3Galb1-4GlcNAcb1-3Galb1-4GlcNAcb1-6(Galb1-3GlcNAcb1-3Galb1-4GlcNAcb1-3Galb1-4GlcNAb1-2)Mana1-6(Galb1-3GlcNAcb1-3Galb1-4GlcNAcb1-3Galb1-4GlcNAcb1-2Mana1-3)Manb1-4GlcNAcb1-4(Fuca1-6)GlcNAcb-Sp24 | 35 | 4 | 13 |
| 366 | Gala1-3Galb1-4(Fuca1-3)GlcNAcb1-2Mana1-6(Gala1-3Galb1-4(Fuca1-3)GlcNAcb1-2Mana1-3)Manb1-4GlcNAcb1-4GlcNAcb-Sp20 | 35 | 2 | 6 |
| 474 | Neu5Aca2-6Galb1-4GlcNAcb1-2Mana1-6(Neu5Aca2-6Galb1-4GlcNAcb1-2Mana1-3)Manb1-4GlcNAcb1-4(Fuca1-6)GlcNAcb-Sp24 | 35 | 8 | 24 |
| 409 | Fuca1-2Galb1-4(Fuca1-3)GlcNAcb1-3GalNAca-Sp14 | 34 | 3 | 8 |
| 477 | Galb1-4GlcNAcb1-6(Galb1-4GlcNAcb1-2)Mana1-6(Galb1-4GlcNAcb1-2Mana1-3)Manb1-4GlcNAcb1-4(Fuca1-6)GlcNAcb-Sp24 | 34 | 13 | 39 |
| 438 | Fuca1-2Galb1-4 GlcNAcb1-2Mana1-6(Fuca1-2Galb1-4GlcNAcb1-2(Fuca1-2Galb1-4GlcNAcb1-4)Mana1-3)Manb1-4GlcNAcb1-4GlcNAcb-Sp12 | 33 | 3 | 8 |

| | | | | |
|---|---|---|---|---|
| 466 | Fuca1-2Galb1-3(Fuca1-4)GlcNAcb1-2Mana1-6(Fuca1-2Galb1-3(Fuca1-4)GlcNAcb1-2Mana1-3)Manb1-4GlcNAcb1-4(Fuca1-6)GlcNAcb1-4(Fuca1-6)GlcNAcb-Sp19 | 33 | 7 | 22 |
| 535 | GlcNAcb1-3Galb1-4GlcNAcb1-3Galb1-4GlcNAcb1-2Mana1-6(GlcNAcb1-3Galb1-4GlcNAcb1-3Galb1-4GlcNAcb1-2Mana1-3)Manb1-4GlcNAcb1-4GlcNAcb-Sp12 | 32 | 5 | 14 |
| 3 | Mana-Sp8 | 32 | 4 | 14 |
| 70 | Fuca1-2Galb1-4(Fuca1-3)GlcNAcb1-3Galb1-4(Fuca1-3)GlcNAcb1-3Galb1-4(Fuca1-3)GlcNAcb-Sp0 | 32 | 3 | 9 |
| 552 | Galb1-4GlcNAcb1-3Galb1-4GlcNAcb1-3Galb1-4GlcNAcb1-3Galb1-4GlcNAcb1-3Galb1-4GlcNAcb1-2Mana1-6(Galb1-4GlcNAcb1-3Galb1-4GlcNAcb1-3Galb1-4GlcNAcb1-3Galb1-4GlcNAcb1-3Galb1-4GlcNAcb1-2Mana1-3)Manb1-4GlcNAcb1-4GlcNAcb-Sp25 | 32 | 7 | 22 |
| 372 | Neu5Aca2-3Galb1-4(Fuca1-3)GlcNAcb1-3GalNAca-Sp14 | 32 | 4 | 13 |
| 547 | Gala1-3Galb1-4GlcNAcb1-2Mana1-6(Gala1-3Galb1-4GlcNAcb1-2Mana1-3)Manb1-4GlcNAcb1-4GlcNAc-Sp24 | 32 | 7 | 23 |
| 581 | Neu5Aca2-6Galb1-4GlcNAcb1-3Galb1-4GlcNAcb1-3Galb1-4GlcNAcb1-2Mana1-6(Neu5Aca2-6Galb1-4GlcNAcb1-3Galb1-4GlcNAcb1-3Galb1-4GlcNAcb1-2Mana1-3)Manb1-4GlcNAcb1-4GlcNAcb-Sp12 | 32 | 11 | 35 |
| 413 | Fuca1-2Galb1-4GlcNAcb1-2Mana1-6(Fuca1-2Galb1-4GlcNAcb1-2Mana1-3)Manb1-4GlcNAcb1-4(Fuca1-6)GlcNAcb-Sp22 | 31 | 2 | 7 |
| 530 | Galb1-3GalNAcb1-3Gal-Sp21 | 31 | 1 | 3 |
| 23 | 6S(3S)Galb1-4GlcNAcb-Sp0 | 30 | 9 | 30 |
| 359 | Fuca1-4(Galb1-3)GlcNAcb1-2Mana1-6(Fuca1-4(Galb1-3)GlcNAcb1-2Mana1-3)Manb1-4GlcNAcb1-4(Fuca1-6)GlcNAcb-Sp22 | 30 | 7 | 23 |
| 560 | Galb1-4GlcNAcb1-3Galb1-4GlcNAcb1-2Mana1-6(Galb1-4GlcNAcb1-3Galb1-4GlcNAcb1-2Mana1-3)Manb1-4GlcNAcb1-4(Fuca1-6)GlcNAcb-Sp24 | 30 | 5 | 17 |
| 273 | Galb1-3(Fuca1-4)GlcNAcb1-3Galb1-3(Fuca1-4)GlcNAcb-Sp0 | 29 | 5 | 15 |
| 437 | (6S)Galb1-3(6S)GlcNAc-Sp0 | 29 | 4 | 12 |
| 249 | Neu5Aca2-3Galb1-4(Fuca1-3)GlcNAcb1-3Galb1-4(Fuca1-3)GlcNAcb1-3Galb1-4(Fuca1-3)GlcNAcb-Sp0 | 29 | 11 | 40 |
| 537 | Galb1-4GlcNAcb1-3Galb1-4GlcNAcb1-3Galb1-4GlcNAcb1-2Mana1-6(Galb1-4GlcNAcb1-3Galb1-4GlcNAcb1-3Galb1-4GlcNAcb1-2Mana1-3)Manb1-4GlcNAcb1-4GlcNAcb-Sp12 | 28 | 4 | 15 |
| 99 | GalNAcb1-4GlcNAcb-Sp8 | 28 | 13 | 48 |
| 439 | Fuca1-2Galb1-4(Fuca1-3)GlcNAcb1-2Mana1-6(Fuca1-2Galb1-4(Fuca1-3)GlcNAcb1-4(Fuca1-2Galb1-4(Fuca1-3)GlcNAcb1-2)Mana1-3)Manb1-4GlcNAcb1-4GlcNAcb-Sp12 | 28 | 12 | 42 |
| 44 | (6S)Galb1-4GlcNAcb-Sp8 | 27 | 18 | 65 |
| 373 | GalNAcb1-4GlcNAcb1-2Mana1-6(GalNAcb1-4GlcNAcb1-2Mana1-3)Manb1-4GlcNAcb1-4GlcNAc-Sp12 | 27 | 10 | 39 |
| 388 | Gala1-3Galb1-3GlcNAcb1-2Mana1-6(Gala1-3Galb1-3GlcNAcb1-2Mana1-3)Manb1-4GlcNAcb1-4GlcNAc-Sp19 | 27 | 4 | 16 |
| 461 | Gala1-3(Fuca1-2)Galb1-3GalNAcb-Sp8 | 26 | 8 | 30 |
| 108 | Gala1-3(Fuca1-2)Galb-Sp18 | 26 | 8 | 30 |
| 15 | GalNAcb-Sp8 | 25 | 8 | 32 |
| 237 | Neu5Aca2-3Galb1-3(Fuca1-4)GlcNAcb1-3Galb1-4(Fuca1-3)GlcNAcb-Sp0 | 25 | 2 | 7 |
| 583 | Neu5Aca2-6Galb1-4GlcNAcb1-3Galb1-4GlcNAcb1-2Mana1-6(Neu5Aca2-6Galb1-4GlcNAcb1-3Galb1-4GlcNAcb1-2Mana1-3)Manb1-4GlcNAcb1-4GlcNAcb-Sp12 | 25 | 3 | 13 |
| 57 | Neu5Aca2-6Galb1-4GlcNAcb1-2Mana1-6(Neu5Aca2-6Galb1-4GlcNAcb1-2Mana1-3)Manb1-4GlcNAcb1-4GlcNAcb-Sp24 | 25 | 2 | 10 |
| 558 | Galb1-3GlcNAcb1-3Galb1-4GlcNAcb1-6(Galb1-3GlcNAcb1-3Galb1-4GlcNAb1-2)Mana1-6(Galb1-3GlcNAcb1-3Galb1-4GlcNAcb1-2Mana1-3)Manb1-4GlcNAcb1-4(Fuca1-6)GlcNAcb-Sp24 | 25 | 5 | 19 |
| 566 | Galb1-4GlcNAcb1-3Galb1-4GlcNAcb1-6(Galb1-4GlcNAcb1-3Galb1-4GlcNAcb1-3)GalNAca-Sp14 | 25 | 5 | 20 |
| 138 | Neu5Aca2-6(Galb1-3)GlcNAcb1-4Galb1-4Glcb-Sp10 | 25 | 12 | 51 |
| 158 | Galb1-4GalNAcb1-3(Fuca1-2)Galb1-4GlcNAcb-Sp8 | 25 | 5 | 22 |

| | | | | |
|---|---|---|---|---|
| 218 | Fuca1-2(6S)Galb1-4GlcNAcb-Sp0 | 25 | 3 | 10 |
| 286 | Galb1-4(Fuca1-3)(6S)GlcNAcb-Sp0 | 25 | 1 | 5 |
| 379 | Galb1-4(Fuca1-3)GlcNAcb1-6(Fuca1-4(Fuca1-2Galb1-3)GlcNAcb1-3)Galb1-4Glc-Sp21 | 24 | 2 | 9 |
| 555 | (3S)GlcAb1-3Galb1-4GlcNAcb1-3Galb1-4Glc-Sp0 | 24 | 2 | 9 |
| 94 | GalNAcb1-3GalNAca-Sp8 | 24 | 1 | 5 |
| 546 | Galb1-4GlcNAcb1-3Galb1-4GlcNAcb1-6(Galb1-4GlcNAcb1-3Galb1-4GlcNAcb1-2)Mana1-6(Galb1-4GlcNAcb1-3Galb1-4GlcNAcb1-2Mana1-3)Mana1-4GlcNAcb1-4GlcNAc-Sp24 | 24 | 9 | 37 |
| 100 | Gala1-2Galb-Sp8 | 24 | 1 | 5 |
| 109 | Gala1-4(Gala1-3)Galb1-4GlcNAcb-Sp8 | 24 | 10 | 41 |
| 228 | GalNAcb1-4(Neu5Aca2-3)Galb1-4GlcNAcb-Sp0 | 24 | 3 | 11 |
| 430 | Galb1-4GlcNAcb1-6(Galb1-4GlcNAcb1-2)Mana1-6(GlcNAcb1-4)(Galb1-4GlcNAcb1-2Mana1-3)Manb1-4GlcNAcb1-4GlcNAc-Sp21 | 24 | 5 | 20 |
| 412 | Galb1-4(Fuca1-3)GlcNAcb1-2Mana1-6(Galb1-4(Fuca1-3)GlcNAcb1-2Mana1-3)Manb1-4GlcNAcb1-4(Fuca1-6)GlcNAcb-Sp22 | 23 | 7 | 31 |
| 11 | Neu5Acb-Sp8 | 23 | 3 | 13 |
| 37 | (3S)Galb1-4GlcNAcb-Sp8 | 23 | 6 | 26 |
| 429 | Galb1-4GlcNAcb1-2Mana1-6(GlcNAcb1-4)(Galb1-4GlcNAcb1-4(Galb1-4GlcNAcb1-2)Mana1-3)Manb1-4GlcNAcb1-4GlcNAc-Sp21 | 23 | 11 | 49 |
| 325 | Neu5Aca2-3Galb1-3(Fuca1-4)GlcNAcb1-3Galb1-3(Fuca1-4)GlcNAcb-Sp0 | 23 | 2 | 9 |
| 368 | Galα1-3(Fucα1-2)Galβ1-3GlcNAcβ1-2Manα1-6(Galα1-3(Fucα1-2)Galβ1-3GlcNAcβ1-2Manα1-3)Manβ1-4GlcNAcβ1-4GlcNAcβ-Sp20 | 23 | 8 | 35 |
| 67 | Fuca1-2Galb1-3GlcNAcb-Sp0 | 22 | 7 | 30 |
| 397 | Gala1-4Galb1-4GlcNAcb1-2Mana1-6(Gala1-4Galb1-4GlcNAcb1-2Mana1-3)Manb1-4GlcNAcb1-4GlcNAcb-Sp24 | 22 | 3 | 13 |
| 85 | GalNAca1-3(Fuca1-2)Galb1-4GlcNAcb-Sp0 | 22 | 5 | 23 |
| 295 | Neu5Aca2-3Galb1-4(Fuca1-3)GlcNAcb1-6(Galb1-3)GalNAca-Sp14 | 22 | 1 | 6 |
| 311 | Mana1-2Mana1-6(Mana1-3)Mana1-6(Mana1-2Mana1-2Mana1-3)Mana-Sp9 | 22 | 11 | 50 |
| 12 | Galb-Sp8 | 22 | 10 | 47 |
| 123 | Gala1-4GlcNAcb-Sp8 | 22 | 2 | 10 |
| 216 | Neu5Aca2-3Galb1-4GlcNAcb1-3Galb1-4(Fuca1-3)GlcNAcb-Sp0 | 22 | 8 | 39 |
| 253 | Neu5Aca2-3Galb1-4(Fuca1-3)GlcNAcb1-3Galb1-4GlcNAcb-Sp8 | 22 | 2 | 8 |
| 14 | Manb-Sp8 | 22 | 1 | 6 |
| 21 | GlcNAcb1-6(GlcNAcb1-4)(GlcNAcb1-3)GlcNAc-Sp8 | 22 | 10 | 45 |
| 91 | GalNAca1-3GalNAcb-Sp8 | 22 | 6 | 27 |
| 240 | Neu5Aca2-6(Neu5Aca2-3Galb1-3)GalNAca-Sp8 | 22 | 10 | 47 |
| 318 | Neu5Gcb2-6Galb1-4GlcNAc-Sp8 | 22 | 5 | 22 |
| 331 | Neu5Aca2-3Galb1-4(Fuca1-3)GlcNAcb1-6(Neu5Aca2-3Galb1-3)GalNAc-Sp14 | 22 | 3 | 12 |
| 451 | Galb1-4GlcNAcb1-6(Galb1-4GlcNAcb1-2)Mana1-6(Galb1-4GlcNAcb1-2Mana1-3)Manb1-4GlcNAcb1-4GlcNAcb-Sp19 | 22 | 1 | 6 |
| 194 | GlcNAcb1-6Galb1-4GlcNAcb-Sp8 | 21 | 1 | 5 |
| 411 | GalNAca1-3(Fuca1-2)Galb1-4(Fuca1-3)GlcNAcb1-3GalNAc-Sp14 | 21 | 5 | 21 |
| 467 | GlcNAcb1-6(GlcNAcb1-2)Mana1-6(GlcNAcb1-2Mana1-3)Manb1-4GlcNAcb1-4(Fuca1-6)GlcNAcb-Sp24 | 21 | 3 | 14 |
| 578 | Neu5Aca2-3Galb1-4GlcNAcb1-3Galb1-4GlcNAcb1-2Mana1-6(Neu5Aca2-3Galb1-4GlcNAcb1-3Galb1-4GlcNAcb1-2Mana1-3)Manb1-4GlcNAcb1-4GlcNAcb-Sp12 | 21 | 6 | 29 |
| 38 | (3S)Galb-Sp8 | 21 | 1 | 7 |
| 505 | (3S)GalNAcb1-4(3S)GlcNAc-Sp8 | 21 | 5 | 23 |
| 17 | GlcNAcb-Sp8 | 21 | 12 | 56 |
| 162 | Galb1-4GlcNAcb1-3Galb1-4GlcNAcb1-3Galb1-4GlcNAcb-Sp0 | 21 | 5 | 26 |
| 497 | Fuca1-2Galb1-3GlcNAcb1-6(Fuca1-2Galb1-3GlcNAcb1-3)GalNAca-Sp14 | 21 | 6 | 29 |
| 538 | Galb1-3GlcNAcb1-3Galb1-4GlcNAcb1-2Mana1-6(Galb1-3GlcNAcb1-3Galb1-4GlcNAcb1-2Mana1-3)Manb1-4GlcNAcb1-4GlcNAc-Sp25 | 21 | 5 | 25 |
| 153 | Galb1-4(Fuca1-3)GlcNAcb1-3Galb1-4(Fuca1-3)GlcNAcb-Sp0 | 21 | 6 | 30 |

| | | | | |
|---|---|---|---|---|
| 352 | KDNa2-3Galb1-4Glc-Sp0 | 21 | 6 | 28 |
| 450 | GalNAca1-3(Fuca1-2)Galb1-3GlcNAcb1-2Mana1-6(GalNAca1-3(Fuca1-2)Galb1-3GlcNAcb1-2Mana1-3)Manb1-4GlcNAcb1-4(Fuca1-6)GlcNAcb-Sp22 | 21 | 3 | 13 |
| 121 | Gala1-4Galb1-4GlcNAcb-Sp8 | 20 | 7 | 33 |
| 187 | GlcNAcb1-6(GlcNAcb1-4)GalNAca-Sp8 | 20 | 9 | 44 |
| 353 | KDNa2-3Galb1-3GalNAca-Sp14 | 20 | 3 | 14 |
| 484 | (3S)Galb1-3(Fuca1-4)GlcNAcb-Sp0 | 20 | 7 | 32 |
| 1 | Gala-Sp8 | 20 | 3 | 17 |
| 42 | (6S)Galb1-4Glcb-Sp0 | 20 | 6 | 32 |
| 110 | Gala1-3GalNAca-Sp8 | 20 | 5 | 25 |
| 13 | Glcb-Sp8 | 20 | 2 | 12 |
| 34 | (3S)Galb1-4(6S)GlcNAcb-Sp0 | 20 | 4 | 20 |
| 112 | Gala1-3GalNAcb-Sp8 | 20 | 1 | 3 |
| 357 | Gala1-3Galb1-4GlcNAcb1-2Mana1-6(Gala1-3Galb1-4GlcNAcb1-2Mana1-3)Manb1-4GlcNAcb1-4GlcNAcb-Sp20 | 20 | 3 | 13 |
| 427 | GlcNAcb1-6(GlcNAcb1-2)Mana1-6(GlcNAcb1-4)(GlcNAcb1-4(GlcNAcb1-2)Mana1-3)Manb1-4GlcNAcb1-4GlcNAc-Sp21 | 20 | 13 | 66 |
| 448 | Gala1-3(Fuca1-2)Galb1-3GlcNAcb1-2Mana1-6(Gala1-3(Fuca1-2)Galb1-3GlcNAcb1-2Mana1-3)Manb1-4GlcNAcb1-4(Fuca1-6)GlcNAcb-Sp22 | 20 | 3 | 17 |
| 503 | (6S)(4S)GalNAcb1-4GlcNAc-Sp8 | 20 | 8 | 39 |
| 250 | Neu5Aca2-3Galb1-4(Fuca1-3)GlcNAcb-Sp0 | 20 | 1 | 7 |
| 364 | GalNAca1-3(Fuca1-2)Galb1-4GlcNAcb1-2Mana1-6(GalNAca1-3(Fuca1-2)Galb1-4GlcNAcb1-2Mana1-3)Manb1-4GlcNAcb1-4GlcNAcb-Sp20 | 20 | 4 | 21 |
| 367 | GalNAca1-3(Fuca1-2)Galb1-3GlcNAcb1-2Mana1-6(GalNAca1-3(Fuca1-2)Galb1-3GlcNAcb1-2Mana1-3)Manb1-4GlcNAcb1-4GlcNAcb-Sp20 | 20 | 5 | 25 |
| 383 | Fuca1-2Galb1-3GalNAca1-3(Fuca1-2)Galb1-4Glcb-Sp0 | 20 | 7 | 36 |
| 420 | Gala1-3(Fuca1-2)Galb1-4GlcNAcb1-2Mana1-6(Gala1-3(Fuca1-2)Galb1-4GlcNAcb1-2Mana1-3)Manb1-4GlcNAcb1-4(Fuca1-6)GlcNAcb-Sp22 | 20 | 3 | 14 |
| 485 | Galb1-4(Fuca1-3)GlcNAcb1-6(Neu5Aca2-6(Neu5Aca2-3Galb1-3)GlcNAcb1-3)Galb1-4Glc-Sp21 | 20 | 3 | 15 |
| 229 | GalNAcb1-4(Neu5Aca2-3)Galb1-4GlcNAcb-Sp8 | 19 | 12 | 63 |
| 261 | Neu5Aca2-6GalNAca-Sp8 | 19 | 4 | 19 |
| 349 | (6S)GlcNAcb1-3Galb1-4GlcNAcb-Sp0 | 19 | 6 | 32 |
| 403 | Galb1-4GlcNAcb1-6(Neu5Aca2-6Galb1-3GlcNAcb1-3)Galb1-4Glc-Sp21 | 19 | 3 | 17 |
| 440 | Galb1-4(Fuca1-3)GlcNAcb1-6GalNAc-Sp14 | 19 | 2 | 8 |
| 584 | GlcNAcb1-3Fuca-Sp21 | 19 | 3 | 16 |
| 8 | Rhaa-Sp8 | 19 | 2 | 11 |
| 9 | Neu5Aca-Sp8 | 19 | 7 | 36 |
| 20 | Galb1-4GlcNAcb1-6(Galb1-4GlcNAcb1-3)GalNAc-Sp14 | 19 | 14 | 73 |
| 135 | Neu5Aca2-6(Galb1-3)GalNAca-Sp8 | 19 | 3 | 15 |
| 179 | GlcNAcb1-3GalNAca-Sp8 | 19 | 1 | 6 |
| 208 | Mana1-2Mana1-6(Mana1-2Mana1-3)Mana-Sp9 | 19 | 4 | 19 |
| 334 | GlcNAca1-4Galb1-3GlcNAcb-Sp0 | 19 | 3 | 18 |
| 346 | GlcNAcb1-2Mana1-6(GlcNAcb1-2Mana1-3)Manb1-4GlcNAcb1-4(Fuca1-6)GlcNAcb-Sp22 | 19 | 7 | 36 |
| 348 | Galb1-3GlcNAcb1-2Mana1-6(Galb1-3GlcNAcb1-2Mana1-3)Manb1-4GlcNAcb1-4(Fuca1-6)GlcNAcb-Sp22 | 19 | 3 | 15 |
| 454 | Neu5Aca2-3Galb1-4GlcNAcb1-6(Neu5Aca2-3Galb1-4GlcNAcb1-2)Mana1-6(GlcNAcb1-4)(Neu5Aca2-3Galb1-4GlcNAcb1-2Mana1-3)Manb1-4GlcNAcb1-4GlcNAcb-Sp21 | 19 | 6 | 34 |
| 486 | Fuca1-2Galb1-4GlcNAcb1-6GalNAca-Sp14 | 19 | 1 | 7 |
| 35 | (3S)Galb1-4(6S)GlcNAcb-Sp8 | 19 | 4 | 23 |
| 239 | Neu5Aca2-3Galb1-3(6S)GalNAca-Sp8 | 19 | 2 | 12 |
| 447 | GalNAca1-3(Fuca1-2)Galb1-4GlcNAcb1-2Mana1-6(GalNAca1-3(Fuca1-2)Galb1-4GlcNAcb1-2Mana1-3)Manb1-4GlcNAcb1-4(Fuca1-6)GlcNAcb-Sp22 | 19 | 2 | 10 |

| | | | | |
|---|---|---|---|---|
| 449 | Neu5Aca2-6Galb1-4GlcNAcb1-6(Fuca1-2Galb1-3GlcNAcb1-3)Galb1-4Glc-Sp21 | 19 | 12 | 64 |
| 493 | Fuca1-2(6S)Galb1-3(6S)GlcNAcb-Sp0 | 19 | 8 | 43 |
| 525 | Fuca1-4(Galb1-3)GlcNAcb1-2 Mana-Sp0 | 19 | 7 | 37 |
| 27 | (3S)Galb1-4(6S)Glcb-Sp8 | 19 | 1 | 5 |
| 61 | Fuca1-2Galb1-3GalNAca-Sp8 | 19 | 3 | 18 |
| 181 | GlcNAcb1-3Galb-Sp8 | 19 | 2 | 13 |
| 263 | Neu5Aca2-6Galb1-4(6S)GlcNAcb-Sp8 | 19 | 5 | 25 |
| 571 | Neu5Aca2-3Galb1-4GlcNAcb1-3Galb1-4GlcNAcb1-6(Neu5Aca2-3Galb1-4GlcNAcb1-3Galb1-4GlcNAcb1-3)GalNAca-Sp14 | 19 | 2 | 10 |
| 157 | Galb1-4GalNAca1-3(Fuca1-2)Galb1-4GlcNAcb-Sp8 | 18 | 2 | 10 |
| 169 | Galb1-4GlcNAcb-Sp8 | 18 | 1 | 5 |
| 47 | (6S)GlcNAcb-Sp8 | 18 | 2 | 10 |
| 50 | Mana1-6(Mana1-3)Manb1-4GlcNAcb1-4GlcNAcb-Sp12 | 18 | 5 | 30 |
| 104 | Gala1-3(Fuca1-2)Galb1-4(Fuca1-3)GlcNAcb-Sp8 | 18 | 2 | 12 |
| 117 | Gala1-3Galb1-4Glc-Sp10 | 18 | 5 | 26 |
| 380 | Galb1-3GlcNAcb1-3Galb1-4(Fuca1-3)GlcNAcb1-6(Galb1-3GlcNAcb1-3)Galb1-4Glc-Sp21 | 18 | 2 | 10 |
| 507 | (3S)GalNAcb1-4GlcNAc-Sp8 | 18 | 4 | 22 |
| 16 | GlcNAcb-Sp0 | 18 | 5 | 25 |
| 103 | Gala1-3(Fuca1-2)Galb1-4(Fuca1-3)GlcNAcb-Sp0 | 18 | 1 | 3 |
| 192 | GlcNAcb1-6GalNAca-Sp8 | 18 | 2 | 13 |
| 445 | Neu5Aca2-8Neu5Aca2-3Galb1-3GalNAcb1-4(Neu5Aca2-8Neu5Aca2-3)Galb1-4Glcb-Sp0 | 18 | 3 | 18 |
| 105 | Gala1-3(Fuca1-2)Galb1-4GlcNAc-Sp0 | 18 | 1 | 6 |
| 119 | Gala1-4(Fuca1-2)Galb1-4GlcNAcb-Sp8 | 18 | 1 | 7 |
| 161 | Galb1-4GlcNAcb1-3Galb1-4(Fuca1-3)GlcNAcb1-3Galb1-4(Fuca1-3)GlcNAcb-Sp0 | 18 | 2 | 14 |
| 209 | Mana1-2Mana1-3Mana-Sp9 | 18 | 18 | 101 |
| 220 | Fuca1-2(6S)Galb1-4(6S)Glcb-Sp0 | 18 | 3 | 18 |
| 435 | GalNAcb1-6GalNAcb-Sp8 | 18 | 3 | 20 |
| 523 | Galb1-4GlcNAcb1-2 Mana1-6(GlcNAcb1-4)(Galb1-4GlcNAcb1-2Mana1-3)Manb1-4GlcNAcb1-4(Fuca1-6)GlcNAc-Sp21 | 18 | 2 | 10 |
| 6 | Fuca-Sp8 | 17 | 4 | 22 |
| 102 | Gala1-3(Fuca1-2)Galb1-3GlcNAcb-Sp8 | 17 | 1 | 7 |
| 133 | GlcNAcb1-6(Galb1-3)GalNAca-Sp8 | 17 | 1 | 6 |
| 232 | Neu5Aca2-6(Neu5Aca2-3)GalNAca-Sp8 | 17 | 5 | 29 |
| 236 | Neu5Aca2-3Galb1-3(Fuca1-4)GlcNAcb-Sp8 | 17 | 2 | 9 |
| 264 | Neu5Aca2-6Galb1-4GlcNAcb-Sp0 | 17 | 3 | 15 |
| 351 | KDNa2-6Galb1-4GlcNAc-Sp0 | 17 | 3 | 16 |
| 432 | Galb1-4Galb-Sp10 | 17 | 9 | 49 |
| 468 | Galb1-3GlcNAcb1-2Mana1-6(GlcNAcb1-4)(Galb1-3GlcNAcb1-2Mana1-3)Manb1-4GlcNAcb1-4GlcNAcb-Sp21 | 17 | 1 | 6 |
| 516 | Galb1-3GlcNAcb1-2Mana-Sp0 | 17 | 6 | 37 |
| 32 | (3S)Galb1-4(Fuca1-3)GlcNAc-Sp0 | 17 | 3 | 19 |
| 48 | Neu5,9Ac$_2$a-Sp8 | 17 | 3 | 19 |
| 64 | Fuca1-2Galb1-3GalNAcb1-4(Neu5Aca2-3)Galb1-4Glcb-Sp9 | 17 | 3 | 17 |
| 83 | GalNAca1-3(Fuca1-2)Galb1-4(Fuca1-3)GlcNAcb-Sp0 | 17 | 3 | 17 |
| 101 | Gala1-3(Fuca1-2)Galb1-3GlcNAcb-Sp0 | 17 | 6 | 38 |
| 207 | Mana1-2Mana1-2Mana1-3Mana-Sp9 | 17 | 2 | 11 |
| 258 | Fuca1-2Galb1-4(6S)Glcb-Sp0 | 17 | 2 | 14 |
| 329 | GalNAca1-3(Fuca1-2)Galb1-4GlcNAcb1-3Galb1-4GlcNAcb-Sp0 | 17 | 3 | 17 |
| 532 | GlcNAcb1-3Galb1-4GlcNAcb1-2Mana1-6(GlcNAcb1-3Galb1-4GlcNAcb1-2Mana1-3)Manb1-4GlcNAcb1-4GlcNAcb-Sp25 | 17 | 3 | 16 |
| 563 | Galb1-4GlcNAcb1-3Galb1-4GlcNAcb1-3Galb1-4GlcNAcb1-6(Galb1-4GlcNAcb1-3Galb1-4GlcNAcb1-3Galb1-4GlcNAb1-2)Mana1-6(Galb1-4GlcNAcb1-3Galb1- | 17 | 2 | 13 |

| | | | | |
|---|---|---|---|---|
| | 4GlcNAcb1-3Galb1-4GlcNAcb1-2Mana1-3)Manb1-4GlcNAcb1-4(Fuca1-6)GlcNAcb-Sp24 | | | |
| 577 | Neu5Aca2-6Galb1-4GlcNAcb1-6(Galb1-3)GalNAca-Sp14 | 17 | 3 | 16 |
| 30 | (3S)Galb1-3GlcNAcb-Sp0 | 17 | 2 | 13 |
| 60 | Fuca1-2Galb1-3(Fuca1-4)GlcNAcb-Sp8 | 17 | 5 | 28 |
| 89 | GalNAca1-3(Fuca1-2)Galb-Sp8 | 17 | 8 | 49 |
| 111 | Gala1-3GalNAca-Sp16 | 17 | 4 | 21 |
| 196 | Glca1-4Glca-Sp8 | 17 | 1 | 3 |
| 245 | Fuca1-2(6S)Galb1-4Glcb-Sp0 | 17 | 1 | 6 |
| 414 | GlcNAcb1-2(GlcNAcb1-6)Mana1-6(GlcNAcb1-2Mana1-3)Manb1-4GlcNAcb1-4GlcNAcb-Sp19 | 17 | 2 | 10 |
| 456 | Neu5Aca2-6Galb1-4GlcNAcb1-2Mana1-6(GlcNAcb1-4)(Neu5Aca2-6Galb1-4GlcNAcb1-2Mana1-3)Manb1-4GlcNAcb1-4GlcNAcb-Sp21 | 17 | 3 | 18 |
| 476 | Mana1-6(Mana1-3)Manb1-4GlcNAcb1-4(Fuca1-6)GlcNAcb-Sp19 | 17 | 1 | 3 |
| 492 | Fuca1-2Galb1-3(6S)GlcNAcb-Sp0 | 17 | 3 | 20 |
| 580 | Neu5Aca2-6Galb1-4GlcNAcb1-3Galb1-4GlcNAcb1-6(Neu5Aca2-6Galb1-4GlcNAcb1-3Galb1-4GlcNAcb1-3)GalNAca-Sp14 | 17 | 3 | 18 |
| 96 | GalNAcb1-3Gala1-4Galb1-4GlcNAcb-Sp0 | 17 | 3 | 16 |
| 120 | Gala1-4Galb1-4GlcNAcb-Sp0 | 17 | 4 | 23 |
| 141 | Galb1-3GalNAca-Sp16 | 17 | 5 | 30 |
| 350 | KDNa2-3Galb1-4(Fuca1-3)GlcNAc-Sp0 | 17 | 2 | 13 |
| 396 | Gala1-4Galb1-3GlcNAcb1-2Mana1-6(Gala1-4Galb1-3GlcNAcb1-2Mana1-3)Manb1-4GlcNAcb1-4GlcNAcb-Sp19 | 17 | 4 | 26 |
| 399 | Galb1-3GlcNAcb1-6Galb1-4GlcNAcb-Sp0 | 17 | 5 | 27 |
| 543 | Neu5Gca2-8Neu5Gca2-6Galb1-4GlcNAc-Sp0 | 17 | 4 | 22 |
| 585 | Galb1-3GalNAcb1-4(Neu5Aca2-8Neu5Aca2-8Neu5Aca2-3)Galb1-4Glcb-Sp21 | 17 | 2 | 14 |
| 72 | Fuca1-2Galb1-4(Fuca1-3)GlcNAcb-Sp8 | 16 | 1 | 6 |
| 74 | Fuca1-2Galb1-4GlcNAcb1-3Galb1-4GlcNAcb1-3Galb1-4GlcNAcb-Sp0 | 16 | 2 | 13 |
| 95 | GalNAcb1-3(Fuca1-2)Galb-Sp8 | 16 | 2 | 11 |
| 131 | Galb1-4GlcNAcb1-6GalNAca-Sp8 | 16 | 2 | 13 |
| 145 | Galb1-3GalNAcb1-4Galb1-4Glcb-Sp8 | 16 | 1 | 8 |
| 166 | Galb1-4GlcNAcb1-6(Galb1-3)GalNAca-Sp8 | 16 | 2 | 11 |
| 377 | Galb1-4(Fuca1-3)GlcNAcb1-6(Galb1-3GlcNAcb1-3)Galb1-4Glc-Sp21 | 16 | 3 | 21 |
| 479 | Neu5Aca2-6Galb1-4GlcNAcb1-6(Fuca1-2Galb1-4(Fuca1-3)GlcNAcb1-3)Galb1-4Glc-Sp21 | 16 | 12 | 71 |
| 499 | GlcNAcb1-6(GlcNAcb1-2)Mana1-6(GlcNAcb1-4)(GlcNAcb1-4(GlcNAcb1-2)Mana1-3)Manb1-4GlcNAcb1-4(Fuca1-6)GlcNAc-Sp21 | 16 | 2 | 11 |
| 10 | Neu5Aca-Sp11 | 16 | 3 | 17 |
| 248 | Neu5Aca2-3Galb1-4(Fuca1-3)(6S)GlcNAcb-Sp8 | 16 | 3 | 20 |
| 319 | Galb1-3GlcNAcb1-2Mana1-6(Galb1-3GlcNAcb1-2Mana1-3)Manb1-4GlcNAcb1-4GlcNAcb-Sp19 | 16 | 2 | 14 |
| 362 | Galb1-4(Fuca1-3)GlcNAcb1-6(Fuca1-2Galb1-4GlcNAcb1-3)Galb1-4Glc-Sp21 | 16 | 5 | 33 |
| 26 | (3S)Galb1-4(6S)Glcb-Sp0 | 16 | 5 | 34 |
| 49 | Neu5,9Ac2a2-6Galb1-4GlcNAcb-Sp8 | 16 | 4 | 23 |
| 52 | GlcNAcb1-2Mana1-6(GlcNAcb1-2Mana1-3)Manb1-4GlcNAcb1-4GlcNAcb-Sp12 | 16 | 1 | 6 |
| 143 | Galb1-3GalNAcb1-3Gala1-4Galb1-4Glcb-Sp0 | 16 | 2 | 10 |
| 197 | Glca1-6Glca1-6Glcb-Sp8 | 16 | 1 | 6 |
| 287 | Galb1-4(Fuca1-3)(6S)Glcb-Sp0 | 16 | 2 | 11 |
| 347 | Galb1-4GlcNAcb1-2Mana1-6(Galb1-4GlcNAcb1-2Mana1-3)Manb1-4GlcNAcb1-4(Fuca1-6)GlcNAcb-Sp22 | 16 | 3 | 21 |
| 360 | Neu5Aca2-6GlcNAcb1-4GlcNAc-Sp21 | 16 | 4 | 23 |
| 385 | Galb1-3GlcNAcb1-3GalNAca-Sp14 | 16 | 4 | 25 |
| 424 | GlcNAcb1-2Mana1-6(GlcNAcb1-4)(GlcNAcb1-2Mana1-3)Manb1-4GlcNAcb1-4GlcNAc-Sp21 | 16 | 2 | 10 |

| | | | | |
|---|---|---|---|---|
| 425 | GlcNAcb1-2Mana1-6(GlcNAcb1-4)(GlcNAcb1-4(GlcNAcb1-2)Mana1-3)Manb1-4GlcNAcb1-4GlcNAc-Sp21 | 16 | 9 | 57 |
| 509 | Galb1-4(6P)GlcNAcb-Sp0 | 16 | 2 | 10 |
| 511 | GalNAca1-3(Fuca1-2)Galb1-4GlcNAcb1-6GalNAc-Sp14 | 16 | 2 | 14 |
| 541 | Neu5Gca2-8Neu5Aca2-3Galb1-4GlcNAc-Sp0 | 16 | 2 | 12 |
| 4 | GalNAca-Sp8 | 16 | 3 | 19 |
| 195 | Glca1-4Glcb-Sp8 | 16 | 2 | 15 |
| 227 | Neu5Aca2-8Neu5Aca2-8Neu5Aca-Sp8 | 16 | 1 | 4 |
| 288 | Galb1-4(Fuca1-3)GlcNAcb1-3Galb1-3(Fuca1-4)GlcNAcb-Sp0 | 16 | 2 | 15 |
| 462 | Glca1-6Glca1-6Glca1-6Glcb-Sp10 | 16 | 2 | 11 |
| 482 | Galb1-3(Fuca1-4)GlcNAcb1-6GalNAca-Sp14 | 16 | 6 | 40 |
| 500 | Galb1-4GlcNAcb1-6(Galb1-4GlcNAcb1-2)Mana1-6(GlcNAcb1-4)Galb1-4GlcNAcb1-4(Gal b1-4GlcNAcb1-2)Mana1-3)Manb1-4GlcNAcb1-4(Fuca1-6)GlcNAc-Sp21 | 16 | 3 | 19 |
| 19 | Galb1-4GlcNAcb1-6(Galb1-4GlcNAcb1-3)GalNAca-Sp8 | 15 | 7 | 44 |
| 46 | Neu5Aca2-3(6S)Galb1-4GlcNAcb-Sp8 | 15 | 8 | 55 |
| 185 | GlcNAcb1-3Galb1-4Glcb-Sp0 | 15 | 3 | 22 |
| 312 | Mana1-2Mana1-6(Mana1-2Mana1-3)Mana1-6(Mana1-2Mana1-2Mana1-3)Mana-Sp9 | 15 | 6 | 38 |
| 376 | Galb1-3GlcNAcb1-3Galb1-4GlcNAcb1-6(Galb1-3GlcNAcb1-3)Galb1-4Glcb-Sp0 | 15 | 2 | 11 |
| 401 | GalNAcb1-3Gala1-6Galb1-4Glcb-Sp8 | 15 | 2 | 15 |
| 436 | (6S)Galb1-3GlcNAcb-Sp0 | 15 | 6 | 36 |
| 483 | Neu5Aca2-3Galb1-3GlcNAcb1-6GalNAca-Sp14 | 15 | 3 | 17 |
| 504 | (6S)GalNAcb1-4GlcNAc-Sp8 | 15 | 4 | 26 |
| 510 | (6P)Galb1-4GlcNAcb-SP0 | 15 | 3 | 20 |
| 139 | Galb1-3GalNAca-Sp8 | 15 | 4 | 24 |
| 154 | Galb1-4(Fuca1-3)GlcNAcb1-3Galb1-4(Fuca1-3)GlcNAcb1-3Galb1-4(Fuca1-3)GlcNAcb-Sp0 | 15 | 2 | 15 |
| 198 | Glcb1-4Glcb-Sp8 | 15 | 1 | 9 |
| 213 | Mana1-6(Mana1-3)Mana1-6(Mana1-2Mana1-3)Manb1-4GlcNAcb1-4GlcNAcb-Sp12 | 15 | 2 | 11 |
| 246 | Neu5Aca2-3Galb1-3GlcNAcb-Sp0 | 15 | 2 | 11 |
| 495 | GalNAcb1-4(Fuca1-3)(6S)GlcNAcb-Sp8 | 15 | 2 | 14 |
| 526 | Neu5Aca2-3Galb1-4(Fuca1-3)GlcNAcb1-2Mana-Sp0 | 15 | 2 | 16 |
| 206 | KDNa2-3Galb1-4GlcNAcb-Sp0 | 15 | 4 | 26 |
| 267 | Neu5Aca2-6Galb1-4GlcNAcb1-3Galb1-4GlcNAcb-Sp0 | 15 | 2 | 12 |
| 330 | GalNAca1-3(Fuca1-2)Galb1-4GlcNAcb1-3Galb1-4GlcNAcb1-3Galb1-4GlcNAcb-Sp0 | 15 | 4 | 27 |
| 363 | Galb1-4GlcNAcb1-2Mana1-6(Galb1-4GlcNAcb1-4(Galb1-4GlcNAcb1-2)Mana1-3)Manb1-4GlcNAcb1-4GlcNAc-Sp21 | 15 | 3 | 20 |
| 404 | Galb1-3GalNAcb1-4(Neu5Aca2-8Neu5Aca2-3)Galb1-4Glcb-Sp0 | 15 | 4 | 24 |
| 498 | GalNAca1-3(Fuca1-2)Galb1-3GlcNAcb1-6GalNAca-Sp14 | 15 | 29 | 198 |
| 514 | Gala1-3(Fuca1-2)Galb1-4GlcNAcb1-2Mana-Sp0 | 15 | 4 | 28 |
| 126 | Galb1-3(Fuca1-4)GlcNAcb1-3Galb1-4(Fuca1-3)GlcNAcb-Sp0 | 15 | 4 | 27 |
| 148 | Galb1-3GlcNAcb1-3Galb1-4Glcb-Sp10 | 15 | 1 | 4 |
| 149 | Galb1-3GlcNAcb-Sp0 | 15 | 2 | 13 |
| 212 | Mana1-2Mana1-2Mana1-6(Mana1-3)Mana-Sp9 | 15 | 1 | 9 |
| 221 | Neu5Aca2-3Galb1-3GalNAca-Sp8 | 15 | 5 | 35 |
| 225 | Neu5Aca2-8Neu5Aca2-8Neu5Aca2-3Galb1-4Glcb-Sp0 | 15 | 3 | 18 |
| 265 | Neu5Aca2-6Galb1-4GlcNAcb-Sp8 | 15 | 2 | 12 |
| 281 | Neu5Gca2-6GalNAca-Sp0 | 15 | 4 | 30 |
| 328 | GalNAcb1-3Gala1-4Galb1-4GlcNAcb1-3Galb1-4Glcb-Sp0 | 15 | 3 | 21 |
| 338 | GlcNAca1-4Galb1-3GalNAc-Sp14 | 15 | 7 | 46 |
| 463 | Glca1-4Glca1-4Glca1-4Glcb-Sp10 | 15 | 2 | 12 |
| 506 | GalNAcb1-4(6S)GlcNAc-Sp8 | 15 | 1 | 4 |
| 529 | Gala1-3(Fuca1-2)Galb1-3GalNAcb1-3Gala1-4Galb1-4Glc-Sp21 | 15 | 3 | 21 |
| 2 | Glca-Sp8 | 14 | 7 | 49 |
| 7 | Fuca-Sp9 | 14 | 9 | 61 |

| | | | | |
|---|---|---|---|---|
| 18 | GlcN(Gc)b-Sp8 | 14 | 10 | 73 |
| 43 | (6S)Galb1-4Glcb-Sp8 | 14 | 2 | 13 |
| 90 | GalNAca1-3(Fuca1-2)Galb-Sp18 | 14 | 2 | 14 |
| 113 | Gala1-3Galb1-4(Fuca1-3)GlcNAcb-Sp8 | 14 | 3 | 19 |
| 118 | Gala1-3Galb-Sp8 | 14 | 3 | 18 |
| 130 | Fuca1-4(Galb1-3)GlcNAcb-Sp8 | 14 | 2 | 12 |
| 142 | Galb1-3GalNAcb-Sp8 | 14 | 2 | 14 |
| 167 | Galb1-4GlcNAcb1-6(Galb1-3)GalNAc-Sp14 | 14 | 2 | 16 |
| 199 | Glcb1-6Glcb-Sp8 | 14 | 3 | 24 |
| 219 | Fuca1-2Galb1-4(6S)GlcNAcb-Sp8 | 14 | 4 | 31 |
| 300 | Galb1-4GlcNAca1-6Galb1-4GlcNAcb-Sp0 | 14 | 2 | 13 |
| 378 | Galb1-4GlcNAcb1-6(Fuca1-4(Fuca1-2Galb1-3)GlcNAcb1-3)Galb1-4Glc-Sp21 | 14 | 2 | 14 |
| 381 | Galb1-4GlcNAcb1-6(Galb1-4GlcNAcb1-2)Mana1-6(Galb1-4GlcNAcb1-4(Galb1-4GlcNAcb1-2)Mana1-3)Manb1-4GlcNAcb1-4GlcNAcb-Sp21 | 14 | 2 | 11 |
| 390 | GlcNAcb1-2Mana1-6(Galb1-4GlcNAcb1-2Mana1-3)Manb1-4GlcNAcb1-4GlcNAc-Sp12 | 14 | 2 | 13 |
| 394 | Galb1-4(Fuca1-3)GlcNAcb1-3GalNAca-Sp14 | 14 | 3 | 20 |
| 442 | Fuca1-2Galb1-4GlcNAcb1-6(Fuca1-2Galb1-4GlcNAcb1-3)GalNAc-Sp14 | 14 | 3 | 23 |
| 508 | (4S)GalNAcb-Sp10 | 14 | 3 | 18 |
| 539 | Neu5Gca2-8Neu5Gca2-3Galb1-4GlcNAc-Sp0 | 14 | 3 | 18 |
| 5 | GalNAca-Sp15 | 14 | 3 | 19 |
| 124 | Gala1-6Glcb-Sp8 | 14 | 7 | 52 |
| 134 | GlcNAcb1-6(Galb1-3)GalNAca-Sp14 | 14 | 2 | 12 |
| 159 | Galb1-4GlcNAcb1-3GalNAca-Sp8 | 14 | 2 | 13 |
| 393 | Fuca1-2Galb1-4GlcNAcb1-3GalNAca-Sp14 | 14 | 5 | 32 |
| 487 | Gala1-3Galb1-4GlcNAcb1-6GalNAca-Sp14 | 14 | 3 | 19 |
| 527 | GlcNAcb1-3Galb1-4GlcNAcb1-6(GlcNAcb1-3)Galb1-4GlcNAc-Sp0 | 14 | 5 | 35 |
| 151 | Galb1-4(Fuca1-3)GlcNAcb-Sp0 | 14 | 3 | 18 |
| 156 | Galb1-4(6S)Glcb-Sp8 | 14 | 1 | 9 |
| 188 | GlcNAcb1-4Galb1-4GlcNAcb-Sp8 | 14 | 3 | 21 |
| 211 | Mana1-6(Mana1-3)Mana-Sp9 | 14 | 4 | 26 |
| 361 | Neu5Aca2-6GlcNAcb1-4GlcNAcb1-4GlcNAc-Sp21 | 14 | 2 | 12 |
| 441 | Galb1-4GlcNAcb1-2Mana-Sp0 | 14 | 5 | 33 |
| 65 | Fuca1-2Galb1-3GlcNAcb1-3Galb1-4Glcb-Sp8 | 14 | 16 | 117 |
| 93 | GalNAca1-4(Fuca1-2)Galb1-4GlcNAcb-Sp8 | 14 | 4 | 27 |
| 186 | GlcNAcb1-4-MDPLys | 14 | 1 | 10 |
| 231 | Neu5Aca2-3Galb1-3GalNAcb1-4(Neu5Aca2-3)Galb1-4Glcb-Sp0 | 14 | 1 | 4 |
| 241 | Neu5Aca2-6(Neu5Aca2-3Galb1-3)GalNAca-Sp14 | 14 | 2 | 18 |
| 277 | Neu5Gca2-3Galb1-3GlcNAcb-Sp0 | 14 | 1 | 4 |
| 289 | Galb1-4GlcNAcb1-3Galb1-3GlcNAcb-Sp0 | 14 | 7 | 51 |
| 314 | Neu5Aca2-6Galb1-4GlcNAcb1-2Mana1-6(Neu5Aca2-3Galb1-4GlcNAcb1-2Mana1-3)Manb1-4GlcNAcb1-4GlcNAcb-Sp12 | 14 | 6 | 44 |
| 400 | Galb1-3GlcNAca1-6Galb1-4GlcNAcb-Sp0 | 14 | 3 | 20 |
| 402 | Gala1-3(Fuca1-2)Galb1-4(Fuca1-3)Glcb-Sp21 | 14 | 6 | 41 |
| 561 | GlcNAcb1-3Galb1-4GlcNAcb1-3Galb1-4GlcNAcb1-2Mana1-6(GlcNAcb1-3Galb1-4GlcNAcb1-3Galb1-4GlcNAcb1-2Mana1-3)Manb1-4GlcNAcb1-4(Fuca1-6)GlcNAcb-Sp24 | 14 | 2 | 18 |
| 565 | Galb1-4GlcNAcb1-3Galb1-4GlcNAcb1-6(Galb1-3)GalNAca-Sp14 | 14 | 5 | 34 |
| 574 | Galb1-4GlcNAcb1-3Galb1-3GalNAca-Sp14 | 14 | 1 | 7 |
| 25 | (3S)Galb1-4Glcb-Sp8 | 13 | 2 | 13 |
| 36 | (3S)Galb1-4GlcNAcb-Sp0 | 13 | 3 | 23 |
| 73 | Fuca1-2Galb1-4GlcNAcb1-3Galb1-4GlcNAcb-Sp0 | 13 | 6 | 47 |
| 168 | Galb1-4GlcNAcb-Sp0 | 13 | 2 | 14 |
| 238 | Neu5Aca2-3Galb1-4(Neu5Aca2-3Galb1-3)GlcNAcb-Sp8 | 13 | 1 | 9 |
| 274 | Neu5Acb2-6GalNAca-Sp8 | 13 | 4 | 29 |

| | | | | |
|---|---|---|---|---|
| 307 | GlcNAcb1-4GlcNAcb-Sp12 | 13 | 1 | 7 |
| 308 | MurNAcb1-4GlcNAcb-Sp10 | 13 | 1 | 7 |
| 405 | Neu5Aca2-3Galb1-3GalNAcb1-4(Neu5Aca2-8Neu5Aca2-3)Galb1-4Glcb-Sp0 | 13 | 3 | 23 |
| 421 | Galb1-3GlcNAcb1-6(Galb1-3GlcNAcb1-2)Mana1-6(Galb1-3GlcNAcb1-2Mana1-3)Manb1-4GlcNAcb1-4GlcNAcb-Sp19 | 13 | 1 | 7 |
| 481 | Gala1-3Galb1-3GlcNAcb1-6GalNAca-Sp14 | 13 | 2 | 11 |
| 522 | GlcNAcb1-2 Mana1-6(GlcNAcb1-4)(GlcNAcb1-2Mana1-3)Manb1-4GlcNAcb1-4(Fuca1-6)GlcNAc-Sp21 | 13 | 3 | 25 |
| 524 | Galb1-4GlcNAcb1-2 Mana1-6(Galb1-4GlcNAcb1-4)(Galb1-4GlcNAcb1-2Mana1-3)Manb1-4GlcNAcb1-4(Fuca1-6)GlcNAc-Sp21 | 13 | 2 | 17 |
| 569 | GlcNAcb1-3Galb1-4GlcNAcb1-6(Galb1-3)GalNAca-Sp14 | 13 | 2 | 14 |
| 28 | (3S)Galb1-3(Fuca1-4)GlcNAcb-Sp8 | 13 | 5 | 38 |
| 33 | (3S)Galb1-4(Fuca1-3)GlcNAc-Sp8 | 13 | 2 | 13 |
| 54 | Galb1-4GlcNAcb1-2Mana1-6(Galb1-4GlcNAcb1-2Mana1-3)Manb1-4GlcNAcb1-4GlcNAcb-Sp12 | 13 | 1 | 11 |
| 122 | Gala1-4Galb1-4Glcb-Sp0 | 13 | 1 | 11 |
| 146 | Galb1-3Galb-Sp8 | 13 | 2 | 14 |
| 170 | Galb1-4GlcNAcb-Sp23 | 13 | 2 | 15 |
| 205 | KDNa2-3Galb1-3GlcNAcb-Sp0 | 13 | 2 | 17 |
| 234 | Neu5Aca2-3GalNAcb1-4GlcNAcb-Sp0 | 13 | 1 | 11 |
| 242 | Neu5Aca2-3Galb-Sp8 | 13 | 3 | 21 |
| 320 | Neu5Aca2-3Galb1-4GlcNAcb1-2Mana1-6(Neu5Aca2-3Galb1-4GlcNAcb1-2Mana1-3)Manb1-4GlcNAcb1-4GlcNAcb-Sp12 | 13 | 1 | 11 |
| 423 | Fuca1-3GlcNAcb1-6(Galb1-4GlcNAcb1-3)Galb1-4Glc-Sp21 | 13 | 1 | 11 |
| 444 | GalNAca1-3(Fuca1-2)Galb1-4GlcNAcb1-6(GalNAca1-3(Fuca1-2)Galb1-4GlcNAcb1-3)GalNAc-Sp14 | 13 | 1 | 6 |
| 460 | Gala1-3(Fuca1-2)Galb1-3GalNAca-Sp8 | 13 | 1 | 6 |
| 488 | Galb1-4(Fuca1-3)GlcNAcb1-2Mana-Sp0 | 13 | 12 | 89 |
| 496 | (3S)GalNAcb1-4(Fuca1-3)GlcNAcb-Sp8 | 13 | 5 | 41 |
| 501 | Galb1-3GlcNAca1-3Galb1-4GlcNAcb-Sp8 | 13 | 1 | 6 |
| 517 | Gala1-3(Fuca1-2)Galb1-3GlcNAcb1-6GalNAc-Sp14 | 13 | 2 | 17 |
| 553 | GlcNAb1-3Galb1-3GalNAc-Sp14 | 13 | 1 | 6 |
| 125 | Galb1-2Galb-Sp8 | 13 | 2 | 16 |
| 247 | Neu5Aca2-3Galb1-4(6S)GlcNAcb-Sp8 | 13 | 1 | 8 |
| 297 | Neu5Aca2-6Galb1-4GlcNAcb1-2Mana1-6(Galb1-4GlcNAcb1-2Mana1-3)Manb1-4GlcNAcb1-4GlcNAcb-Sp12 | 13 | 1 | 8 |
| 315 | Galb1-4GlcNAcb1-2Mana1-6(Neu5Aca2-6Galb1-4GlcNAcb1-2Mana1-3)Manb1-4GlcNAcb1-4GlcNAcb-Sp12 | 13 | 2 | 12 |
| 327 | Gala1-4Galb1-4GlcNAcb1-3Galb1-4Glcb-Sp0 | 13 | 1 | 4 |
| 358 | Galb1-4GlcNAcb1-2Mana1-6(Mana1-3)Manb1-4GlcNAcb1-4GlcNAcb-Sp12 | 13 | 3 | 21 |
| 369 | Fuca1-4(Fuca1-2Galb1-3)GlcNAcb1-2Mana1-3(Fuca1-4(Fuca1-2Galb1-3)GlcNAcb1-2Mana1-3)Manb1-4GlcNAcb1-4GlcNAcb-Sp19 | 13 | 3 | 23 |
| 371 | Neu5Aca2-6Galb1-4GlcNAcb1-3GalNAc-Sp14 | 13 | 2 | 17 |
| 406 | Gala1-3(Fuca1-2)Galb1-4GlcNAcb1-3GalNAca-Sp14 | 13 | 8 | 65 |
| 452 | Neu5Aca2-3Galb1-4GlcNAcb1-2Mana1-6(GlcNAcb1-4)(Neu5Aca2-3Galb1-4GlcNAcb1-2Mana1-3)Manb1-4GlcNAcb1-4GlcNAcb-Sp21 | 13 | 2 | 13 |
| 55 | Neu5Aca2-6Galb1-4GlcNAcb1-2Mana1-6(Neu5Aca2-6Galb1-4GlcNAcb1-2Mana1-3)Manb1-4GlcNAcb1-4GlcNAcb-Sp12 | 13 | 2 | 14 |
| 75 | Fuca1-2Galb1-4GlcNAcb-Sp0 | 13 | 3 | 21 |
| 116 | Gala1-3Galb1-4Glcb-Sp0 | 13 | 4 | 32 |
| 136 | Neu5Aca2-6(Galb1-3)GalNAca-Sp14 | 13 | 1 | 8 |
| 180 | GlcNAcb1-3GalNAca-Sp14 | 13 | 1 | 10 |
| 193 | GlcNAcb1-6GalNAca-Sp14 | 13 | 3 | 25 |
| 255 | Neu5Aca2-3Galb1-4GlcNAcb-Sp0 | 13 | 3 | 20 |
| 280 | Neu5Gca2-3Galb1-4Glcb-Sp0 | 13 | 2 | 19 |

| | | | | |
|---|---|---|---|---|
| 333 | GlcNAca1-4Galb1-4GlcNAcb-Sp0 | 13 | 5 | 44 |
| 386 | GalNAcb1-4(Neu5Aca2-3)Galb1-4GlcNAcb1-3GalNAca-Sp14 | 13 | 3 | 27 |
| 387 | GalNAca1-3(Fuca1-2)Galb1-3GalNAca1-3(Fuca1-2)Galb1-4GlcNAcb-Sp0 | 13 | 3 | 27 |
| 453 | Neu5Aca2-3Galb1-4GlcNAcb1-4Mana1-6(GlcNAcb1-4)(Neu5Aca2-3Galb1-4GlcNAcb1-4(Neu5Aca2-3Galb1-4GlcNAcb1-2)Mana1-3)Manb1-4GlcNAcb1-4GlcNAcb-Sp21 | 13 | 5 | 41 |
| 489 | Fuca1-2(6S)Galb1-3GlcNAcb-Sp0 | 13 | 8 | 63 |
| 491 | Fuca1-2Galb1-4GlcNAcb1-2Mana-Sp0 | 13 | 2 | 15 |
| 494 | Neu5Aca2-6GalNAcb1-4(6S)GlcNAcb-Sp8 | 13 | 2 | 14 |
| 549 | GalNAcb1-3GlcNAcb-Sp0 | 13 | 1 | 10 |
| 66 | Fuca1-2Galb1-3GlcNAcb1-3Galb1-4Glcb-Sp10 | 12 | 2 | 12 |
| 152 | Galb1-4(Fuca1-3)GlcNAcb-Sp8 | 12 | 3 | 22 |
| 215 | Manb1-4GlcNAcb-Sp0 | 12 | 2 | 18 |
| 259 | Neu5Aca2-3Galb1-4Glcb-Sp0 | 12 | 4 | 36 |
| 260 | Neu5Aca2-3Galb1-4Glcb-Sp8 | 12 | 3 | 25 |
| 321 | Neu5Aca2-3Galb1-4GlcNAcb1-2Mana1-6(Neu5Aca2-6Galb1-4GlcNAcb1-2Mana1-3)Manb1-4GlcNAcb1-4GlcNAcb-Sp12 | 12 | 2 | 15 |
| 343 | Galb1-4GlcNAcb1-2Mana1-3Manb1-4GlcNAcb1-4GlcNAc-Sp12 | 12 | 1 | 10 |
| 365 | Gala1-3(Fuca1-2)Galb1-4GlcNAcb1-2Mana1-6(Gala1-3(Fuca1-2)Galb1-4GlcNAcb1-2Mana1-3)Manb1-4GlcNAcb1-4GlcNAcb-Sp20 | 12 | 2 | 17 |
| 471 | Neu5Aca2-3Galb1-4GlcNAcb1-6GalNAca-Sp14 | 12 | 4 | 32 |
| 475 | Neu5Aca2-3Galb1-4GlcNAcb1-2Mana1-6(Neu5Aca2-3Galb1-4GlcNAcb1-2Mana1-3)Manb1-4GlcNAcb1-4(Fuca1-6)GlcNAcb-Sp24 | 12 | 11 | 92 |
| 521 | Neu5Aca2-3Galb1-3GalNAcb1-4Galb1-4Glcb-Sp0 | 12 | 6 | 52 |
| 533 | Galβ1-4GlcNAcβ1-3Galβ1-4GlcNAcβ1-2Manα1-6(Galβ1-4GlcNAcβ1-3Galβ1-4GlcNAcβ1-2Manα1-3)Manβ1-4GlcNAcβ1-4GlcNAcβ-Sp12 | 12 | 2 | 12 |
| 564 | Galb1-4GlcNAcb1-3Galb1-4GlcNAcb1-3GalNAca-Sp14 | 12 | 4 | 29 |
| 568 | GlcNAcb1-3Galb1-4GlcNAcb1-3GalNAca-Sp14 | 12 | 1 | 8 |
| 31 | (3S)Galb1-3GlcNAcb-Sp8 | 12 | 1 | 10 |
| 51 | Mana1-6(Mana1-3)Manb1-4GlcNAcb1-4GlcNAcb-Sp13 | 12 | 2 | 17 |
| 62 | Fuca1-2Galb1-3GalNAca-Sp14 | 12 | 27 | 223 |
| 254 | Neu5Aca2-3Galb1-4GlcNAcb1-3Galb1-4GlcNAcb1-3Galb1-4GlcNAcb-Sp0 | 12 | 5 | 40 |
| 270 | Neu5Aca2-6Galb-Sp8 | 12 | 1 | 10 |
| 272 | Neu5Aca2-8Neu5Aca2-3Galb1-4Glcb-Sp0 | 12 | 5 | 41 |
| 324 | Neu5Aca2-6Galb1-4GlcNAcb1-3Galb1-3GlcNAcb-Sp0 | 12 | 1 | 12 |
| 339 | Neu5Aca2-6Galb1-4GlcNAcb1-2Mana1-6(Mana1-3)Manb1-4GlcNAcb1-4GlcNAc-Sp12 | 12 | 1 | 10 |
| 422 | Galb1-4GlcNAcb1-6(Fuca1-2Galb1-3GlcNAcb1-3)Galb1-4Glc-Sp21 | 12 | 1 | 7 |
| 431 | Galb1-4GlcNAcb1-6(Galb1-4GlcNAcb1-2)Mana1-6(GlcNAcb1-4)(Galb1-4GlcNAcb1-4(Galb1-4GlcNAcb1-2)Mana1-3)Manb1-4GlcNAcb1-4GlcNAc-Sp21 | 12 | 1 | 12 |
| 446 | GalNAcb1-4Galb1-4Glcb-Sp0 | 12 | 6 | 50 |
| 519 | Gala1-3Galb1-3GlcNAcb1-2Mana-Sp0 | 12 | 2 | 15 |
| 520 | GalNAcb1-4GlcNAcb1-2Mana-Sp0 | 12 | 1 | 12 |
| 29 | (3S)Galb1-3GalNAca-Sp8 | 12 | 2 | 13 |
| 298 | Galb1-4GlcNAcb1-6(Galb1-4GlcNAcb1-3)Galb1-4GlcNAc-Sp0 | 12 | 2 | 19 |
| 342 | Neu5Aca2-6Galb1-4GlcNAcb1-2Mana1-3Manb1-4GlcNAcb1-4GlcNAc-Sp12 | 12 | 1 | 4 |
| 419 | Fuca1-2Galb1-3GlcNAcb1-2Mana1-6(Fuca1-2Galb1-3GlcNAcb1-2Mana1-3)Manb1-4GlcNAcb1-4(Fuca1-6)GlcNAcb-Sp22 | 12 | 3 | 28 |
| 53 | GlcNAcb1-2Mana1-6(GlcNAcb1-2Mana1-3)Manb1-4GlcNAcb1-4GlcNAcb-Sp13 | 12 | 2 | 18 |
| 81 | Fucb1-3GlcNAcb-Sp8 | 12 | 1 | 9 |
| 132 | Galb1-4GlcNAcb1-6GalNAc-Sp14 | 12 | 3 | 25 |
| 164 | Galb1-4GlcNAcb1-3Galb1-4Glcb-Sp0 | 12 | 7 | 59 |
| 256 | Neu5Aca2-3Galb1-4GlcNAcb-Sp8 | 12 | 2 | 18 |
| 276 | Neu5Gca2-3Galb1-3(Fuca1-4)GlcNAcb-Sp0 | 12 | 3 | 27 |
| 304 | Neu5Aca2-6Galb1-4GlcNAcb1-2Mana1-6(GlcNAcb1-2Mana1-3)Manb1-4GlcNAcb1-4GlcNAcb-Sp12 | 12 | 2 | 21 |

| | | | | |
|---|---|---|---|---|
| 428 | Galb1-4GlcNAcb1-2Mana1-6(GlcNAcb1-4)(Galb1-4GlcNAcb1-2Mana1-3)Manb1-4GlcNAcb1-4GlcNAc-Sp21 | 12 | 3 | 27 |
| 550 | GalNAcb1-4GlcNAcb1-3GalNAcb1-4GlcNAcb-Sp0 | 12 | 3 | 23 |
| 45 | (6S)Galb1-4(6S)Glcb-Sp8 | 11 | 3 | 22 |
| 56 | Neu5Aca2-6Galb1-4GlcNAcb1-2Mana1-6(Neu5Aca2-6Galb1-4GlcNAcb1-2Man-a1-3)Manb1-4GlcNAcb1-4GlcNAcb-Sp21 | 11 | 3 | 23 |
| 76 | Fuca1-2Galb1-4GlcNAcb-Sp8 | 11 | 5 | 41 |
| 107 | Gala1-3(Fuca1-2)Galb-Sp8 | 11 | 2 | 18 |
| 155 | Galb1-4(6S)Glcb-Sp0 | 11 | 2 | 17 |
| 214 | Mana1-6(Mana1-3)Mana1-6(Mana1-3)Manb1-4GlcNAcb1-4GlcNAcb-Sp12 | 11 | 8 | 74 |
| 257 | Neu5Aca2-3Galb1-4GlcNAcb1-3Galb1-4GlcNAcb-Sp0 | 11 | 3 | 26 |
| 283 | Neu5Gca-Sp8 | 11 | 2 | 20 |
| 392 | Neu5Aca2-3Galb1-3GlcNAcb1-3GalNAca-Sp14 | 11 | 1 | 4 |
| 472 | Neu5Aca2-6Galb1-4GlcNAcb1-6GalNAca-Sp14 | 11 | 1 | 9 |
| 518 | Neu5Aca2-3Galb1-3GlcNAcb1-2Mana-Sp0 | 11 | 1 | 9 |
| 542 | Neu5Gca2-8Neu5Gca2-3Galb1-4GlcNAcb1-3Galb1-4GlcNAc-Sp0 | 11 | 4 | 34 |
| 559 | Neu5Aca2-8Neu5Aca2-3Galb1-3GalNAcb1-4(Neu5Aca2-3)Galb1-4Glc-Sp21 | 11 | 3 | 22 |
| 576 | Neu5Aca2-6Galb1-4GlcNAcb1-3Galb1-4GlcNAcb1-6(Galb1-3)GalNAca-Sp14 | 11 | 11 | 102 |
| 92 | GalNAca1-3Galb-Sp8 | 11 | 3 | 26 |
| 115 | Gala1-3Galb1-4GlcNAcb-Sp8 | 11 | 9 | 80 |
| 160 | Galb1-4GlcNAcb1-3GalNAc-Sp14 | 11 | 1 | 7 |
| 269 | Neu5Aca2-6Galb1-4Glcb-Sp8 | 11 | 1 | 13 |
| 284 | Neu5Aca2-3Galb1-4GlcNAcb1-6(Galb1-3)GalNAca-Sp14 | 11 | 3 | 29 |
| 294 | (6P)Glcb-Sp10 | 11 | 3 | 27 |
| 326 | Neu5Aca2-6Galb1-4GlcNAcb1-3Galb1-4GlcNAcb1-3Galb1-4GlcNAcb-Sp0 | 11 | 1 | 7 |
| 469 | Neu5Aca2-6Galb1-4GlcNAcb1-6(Galb1-3GlcNAcb1-3)Galb1-4Glcb-Sp21 | 11 | 2 | 15 |
| 528 | GalNAca1-3(Fuca1-2)Galb1-3GalNAcb1-3Gala1-4Galb1-4Glc-Sp21 | 11 | 2 | 22 |
| 579 | GlcNAcb1-6(Neu5Aca2-3Galb1-3)GalNAca-Sp14 | 11 | 1 | 7 |
| 128 | Galb1-3(Fuca1-4)GlcNAc-Sp0 | 11 | 7 | 67 |
| 323 | Neu5,9Ac2a2-3Galb1-3GlcNAcb-Sp0 | 11 | 3 | 23 |
| 341 | Neu5Aca2-6Galb1-4GlcNAcb1-2Mana1-6Manb1-4GlcNAcb1-4GlcNAc-Sp12 | 11 | 1 | 9 |
| 398 | Gala1-3Galb1-4GlcNAcb1-3GalNAca-Sp14 | 11 | 2 | 16 |
| 417 | GalNAca1-3(Fuca1-2)Galb1-3GlcNAcb1-3GalNAc-Sp14 | 11 | 2 | 19 |
| 434 | Neu5Aca2-3Galb1-4GlcNAcb1-3Galb-Sp8 | 11 | 2 | 14 |
| 464 | Neu5Aca2-3Galb1-4GlcNAcb1-6(Neu5Aca2-3Galb1-4GlcNAcb1-3)GalNAca-Sp14 | 11 | 2 | 19 |
| 513 | Gala1-3Galb1-4GlcNAcb1-2Mana-Sp0 | 11 | 3 | 23 |
| 554 | Galb1-3GlcNAcb1-6(Galb1-3)GalNAc-Sp14 | 11 | 2 | 14 |
| 575 | Neu5Aca2-3Galb1-4GlcNAcb1-3Galb1-4GlcNAcb1-6(Galb1-3)GalNAca-Sp14 | 11 | 3 | 23 |
| 59 | Fuca1-2Galb1-3GalNAcb1-3Gala1-4Galb1-4Glcb-Sp9 | 11 | 1 | 12 |
| 71 | Fuca1-2Galb1-4(Fuca1-3)GlcNAcb-Sp0 | 11 | 1 | 5 |
| 163 | Galb1-4GlcNAcb1-3Galb1-4GlcNAcb-Sp0 | 11 | 1 | 12 |
| 171 | Galb1-4Glcb-Sp0 | 11 | 3 | 30 |
| 252 | Neu5Aca2-3Galb1-4(Fuca1-3)GlcNAcb1-3Galb-Sp8 | 11 | 3 | 25 |
| 271 | Neu5Aca2-8Neu5Aca-Sp8 | 11 | 2 | 18 |
| 340 | Mana1-6(Neu5Aca2-6Galb1-4GlcNAcb1-2Mana1-3)Manb1-4GlcNAcb1-4GlcNAc-Sp12 | 11 | 4 | 33 |
| 470 | Neu5Aca2-3Galb1-4GlcNAcb1-2Mana-Sp0 | 11 | 6 | 55 |
| 77 | Fuca1-2Galb1-4Glcb-Sp0 | 10 | 2 | 17 |
| 127 | Galb1-3GlcNAcb1-3Galb1-4(Fuca1-3)GlcNAcb-Sp0 | 10 | 1 | 9 |
| 140 | Galb1-3GalNAca-Sp14 | 10 | 3 | 30 |
| 175 | GlcNAcb1-2Galb1-3GalNAca-Sp8 | 10 | 3 | 26 |
| 230 | GalNAcb1-4(Neu5Aca2-3)Galb1-4Glcb-Sp0 | 10 | 2 | 23 |
| 233 | Neu5Aca2-3GalNAca-Sp8 | 10 | 1 | 9 |
| 382 | GlcNAcb1-2Mana1-6(GlcNAcb1-4(GlcNAcb1-2)Mana1-3)Manb1-4GlcNAcb1-4GlcNAc-Sp21 | 10 | 1 | 9 |

| | | | | |
|---|---|---|---|---|
| 408 | GalNAca1-3GalNAcb1-3Gala1-4Galb1-4Glcb-Sp0 | 10 | 3 | 32 |
| 433 | Galb1-6Galb-Sp10 | 10 | 2 | 20 |
| 458 | Neu5Aca2-6Galb1-4GlcNAcb1-6(Neu5Aca2-6Galb1-4GlcNAcb1-2)Mana1-6(GlcNAcb1-4)(Neu5Aca2-6Galb1-4GlcNAcb1-2Mana1-3)Manb1-4GlcNAcb1-4GlcNAcb-Sp21 | 10 | 3 | 28 |
| 544 | Neu5Aca2-8Neu5Aca2-3Galb1-4GlcNAc-Sp0 | 10 | 1 | 5 |
| 68 | Fuca1-2Galb1-3GlcNAcb-Sp8 | 10 | 2 | 16 |
| 176 | GlcNAcb1-6(GlcNAcb1-3)GalNAca-Sp8 | 10 | 2 | 18 |
| 203 | GlcAb1-3Galb-Sp8 | 10 | 2 | 16 |
| 210 | Mana1-2Mana1-6(Mana1-2Mana1-3)Mana1-6(Mana1-2Mana1-2Mana1-3)Manb1-4GlcNAcb1-4GlcNAcb-Sp12 | 10 | 1 | 14 |
| 540 | Neu5Aca2-8Neu5Gca2-3Galb1-4GlcNAc-Sp0 | 10 | 5 | 47 |
| 178 | GlcNAcb1-6(GlcNAcb1-3)Galb1-4GlcNAcb-Sp8 | 10 | 1 | 10 |
| 204 | GlcAb1-6Galb-Sp8 | 10 | 4 | 36 |
| 268 | Neu5Aca2-6Galb1-4Glcb-Sp0 | 10 | 2 | 15 |
| 375 | Galb1-3GalNAca1-3(Fuca1-2)Galb1-4GlcNAc-Sp0 | 10 | 3 | 29 |
| 455 | Neu5Aca2-3Galb1-4GlcNAcb1-6(Neu5Aca2-3Galb1-4GlcNAcb1-2)Mana1-6(GlcNAcb1-4)(Neu5Aca2-3Galb1-4GlcNAcb1-4(Neu5Aca2-3Galb1-4GlcNAcb1-2)Mana1-3)Manb1-4GlcNAcb1-4GlcNAcb-Sp21 | 10 | 2 | 19 |
| 79 | Fuca1-3GlcNAcb-Sp8 | 10 | 2 | 18 |
| 165 | Galb1-4GlcNAcb1-3Galb1-4Glcb-Sp8 | 10 | 3 | 30 |
| 201 | GlcAa-Sp8 | 10 | 2 | 20 |
| 224 | GalNAcb1-4(Neu5Aca2-8Neu5Aca2-8Neu5Aca2-3)Galb1-4Glcb-Sp0 | 10 | 3 | 28 |
| 226 | GalNAcb1-4(Neu5Aca2-8Neu5Aca2-3)Galb1-4Glcb-Sp0 | 10 | 2 | 18 |
| 235 | Neu5Aca2-3Galb1-3(6S)GlcNAc-Sp8 | 10 | 3 | 33 |
| 306 | GlcNAcb1-4GlcNAcb-Sp10 | 10 | 2 | 25 |
| 322 | Galb1-4(Fuca1-3)GlcNAcb1-2Mana1-6(Galb1-4(Fuca1-3)GlcNAcb1-2Mana1-3)Manb1-4GlcNAcb1-4GlcNAcb-Sp20 | 10 | 1 | 6 |
| 551 | GlcNAcb1-3Galb1-4GlcNAcb1-3Galb1-4GlcNAcb1-3Galb1-4GlcNAcb1-3Galb1-4GlcNAcb1-2Mana1-6(GlcNAcb1-3Galb1-4GlcNAcb1-3Galb1-4GlcNAcb1-3Galb1-4GlcNAcb1-3Galb1-4GlcNAcb1-2Mana1-3)Manb1-4GlcNAcb1-4GlcNAcb-Sp25 | 10 | 3 | 28 |
| 572 | Neu5Aca2-6Galb1-4GlcNAcb1-3Galb1-4GlcNAcb1-3GalNAca-Sp14 | 10 | 4 | 40 |
| 41 | (6P)Mana-Sp8 | 9 | 3 | 30 |
| 58 | Fuca1-2Galb1-3GalNAcb1-3Gala-Sp9 | 9 | 2 | 26 |
| 63 | Fuca1-2Galb1-3GalNAcb1-4(Neu5Aca2-3)Galb1-4Glcb-Sp0 | 9 | 1 | 14 |
| 147 | Galb1-3GlcNAcb1-3Galb1-4GlcNAcb-Sp0 | 9 | 3 | 28 |
| 189 | GlcNAcb1-4GlcNAcb1-4GlcNAcb1-4GlcNAcb1-4GlcNAcb1-4GlcNAcb1-Sp8 | 9 | 2 | 22 |
| 191 | GlcNAcb1-4GlcNAcb1-4GlcNAcb-Sp8 | 9 | 1 | 10 |
| 200 | G-ol-Sp8 | 9 | 4 | 44 |
| 223 | GalNAcb1-4(Neu5Aca2-8Neu5Aca2-8Neu5Aca2-8Neu5Aca2-3)Galb1-4Glcb-Sp0 | 9 | 4 | 44 |
| 278 | Neu5Gca2-3Galb1-4(Fuca1-3)GlcNAcb-Sp0 | 9 | 2 | 18 |
| 302 | GalNAcb1-3Galb-Sp8 | 9 | 2 | 16 |
| 415 | Fuca1-2Galb1-3GlcNAcb1-3GalNAc-Sp14 | 9 | 2 | 24 |
| 515 | GalNAca1-3(Fuca1-2)Galb1-4 GlcNAcb1-2Mana-Sp0 | 9 | 1 | 5 |
| 78 | Fuca1-2Galb-Sp8 | 9 | 1 | 9 |
| 84 | (3S)Galb1-4(Fuca1-3)Glcb-Sp0 | 9 | 15 | 168 |
| 144 | Galb1-3GalNAcb1-4(Neu5Aca2-3)Galb1-4Glcb-Sp0 | 9 | 3 | 33 |
| 173 | GlcNAca1-3Galb1-4GlcNAcb-Sp8 | 9 | 5 | 51 |
| 243 | Neu5Aca2-3Galb1-3GalNAcb1-3Gala1-4Galb1-4Glcb-Sp0 | 9 | 3 | 33 |
| 251 | Neu5Aca2-3Galb1-4(Fuca1-3)GlcNAcb-Sp8 | 9 | 1 | 13 |
| 282 | Neu5Gca2-6Galb1-4GlcNAcb-Sp0 | 9 | 3 | 37 |
| 292 | 4S(3S)Galb1-4GlcNAcb-Sp0 | 9 | 1 | 16 |
| 416 | Gala1-3(Fuca1-2)Galb1-3GlcNAcb1-3GalNAc-Sp14 | 9 | 2 | 26 |
| 457 | Neu5Aca2-6Galb1-4GlcNAcb1-4Mana1-6(GlcNAcb1-4)(Neu5Aca2-6Galb1-4GlcNAcb1-4(Neu5Aca2-6Galb1-4GlcNAcb1-2)Mana1-3)Manb1-4GlcNAcb1-4GlcNAcb-Sp21 | 9 | 4 | 40 |

| | | | | |
|---|---|---|---|---|
| 490 | Gala1-3(Fuca1-2)Galb1-4GlcNAcb1-6GalNAca-Sp14 | 9 | 2 | 24 |
| 172 | Galb1-4Glcb-Sp8 | 9 | 2 | 17 |
| 182 | GlcNAcb1-3Galb1-4GlcNAcb-Sp0 | 9 | 1 | 11 |
| 291 | Neu5Aca2-3Galb1-4GlcNAcb1-3Galb1-3GlcNAcb-Sp0 | 9 | 3 | 29 |
| 303 | GlcAb1-3GlcNAcb-Sp8 | 9 | 2 | 25 |
| 310 | Mana1-6(Mana1-3)Mana1-6(Mana1-3)Manb-Sp10 | 9 | 1 | 6 |
| 345 | Mana1-6(Galb1-4GlcNAcb1-2Mana1-3)Manb1-4GlcNAcb1-4GlcNAcb-Sp12 | 9 | 1 | 6 |
| 556 | (3S)GlcAb1-3Galb1-4GlcNAcb1-2Mana-Sp0 | 9 | 2 | 22 |
| 80 | Fuca1-4GlcNAcb-Sp8 | 9 | 1 | 15 |
| 106 | Gala1-3(Fuca1-2)Galb1-4Glcb-Sp0 | 9 | 2 | 24 |
| 384 | Fuca1-2Galb1-3GalNAca1-3(Fuca1-2)Galb1-4GlcNAcb-Sp0 | 9 | 4 | 51 |
| 129 | Galb1-3(Fuca1-4)GlcNAc-Sp8 | 8 | 2 | 25 |
| 480 | Galb1-3GlcNAcb1-6GalNAca-Sp14 | 8 | 7 | 86 |
| 202 | GlcAb-Sp8 | 8 | 6 | 81 |
| 222 | Neu5Aca2-3Galb1-3GalNAca-Sp14 | 8 | 1 | 10 |
| 344 | Galb1-4GlcNAcb1-2Mana1-6Manb1-4GlcNAcb1-4GlcNAc-Sp12 | 8 | 7 | 87 |
| 374 | Galb1-3GalNAca1-3(Fuca1-2)Galb1-4Glc-Sp0 | 8 | 3 | 37 |
| 391 | Galb1-4GlcNAcb1-2Mana1-6(GlcNAcb1-2Mana1-3)Manb1-4GlcNAcb1-4GlcNAc-Sp12 | 8 | 0 | 0 |
| 570 | GlcNAcb1-3Galb1-4GlcNAcb1-6(GlcNAcb1-3Galb1-4GlcNAcb1-3)GalNAca-Sp14 | 8 | 2 | 20 |
| 87 | GalNAca1-3(Fuca1-2)Galb1-4Glcb-Sp0 | 8 | 3 | 32 |
| 290 | Neu5Aca2-3Galb1-3GlcNAcb1-3Galb1-3GlcNAcb-Sp0 | 8 | 2 | 26 |
| 296 | Galb1-3Galb1-4GlcNAcb-Sp8 | 8 | 1 | 17 |
| 370 | Neu5Aca2-3Galb1-4GlcNAcb1-3GalNAc-Sp14 | 8 | 2 | 28 |
| 279 | Neu5Gca2-3Galb1-4GlcNAcb-Sp0 | 7 | 1 | 13 |
| 309 | Mana1-6Manb-Sp10 | 7 | 1 | 17 |
| 337 | GlcNAca1-4Galb1-4GlcNAcb1-3Galb1-4GlcNAcb-Sp0 | 7 | 7 | 97 |
| 395 | GalNAca1-3GalNAcb1-3Gala1-4Galb1-4GlcNAcb-Sp0 | 7 | 3 | 41 |
| 177 | GlcNAcb1-6(GlcNAcb1-3)GalNAca-Sp14 | 7 | 2 | 31 |
| 305 | GlcNAcb1-3Man-Sp10 | 7 | 3 | 39 |
| 418 | Gala1-3Galb1-3GlcNAcb1-3GalNAc-Sp14 | 7 | 2 | 31 |
| 548 | GlcNAcb1-3Galb1-4GlcNAcb1-6(GlcNAcb1-3Galb1-3)GalNAca-Sp14 | 7 | 2 | 23 |
| 82 | GalNAca1-3(Fuca1-2)Galb1-3GlcNAcb-Sp0 | 7 | 1 | 7 |
| 150 | Galb1-3GlcNAcb-Sp8 | 7 | 1 | 14 |
| 183 | GlcNAcb1-3Galb1-4GlcNAcb-Sp8 | 7 | 1 | 9 |
| 410 | Gala1-3(Fuca1-2)Galb1-4(Fuca1-3)GlcNAcb1-3GalNAc-Sp14 | 7 | 1 | 15 |
| 174 | GlcNAca1-6Galb1-4GlcNAcb-Sp8 | 6 | 1 | 20 |
| 190 | GlcNAcb1-4GlcNAcb1-4GlcNAcb1-4GlcNAcb1-4GlcNAcb1-Sp8 | 6 | 2 | 24 |
| 299 | GlcNAcb1-6(Galb1-4GlcNAcb1-3)Galb1-4GlcNAc-Sp0 | 6 | 5 | 82 |
| 317 | Neu5Aca2-8Neu5Aca2-8Neu5Acb-Sp8 | 6 | 2 | 33 |
| 512 | Neu5Aca2-6Galb1-4GlcNAcb1-2Man-Sp0 | 6 | 3 | 42 |
| 69 | Fuca1-2Galb1-4(Fuca1-3)GlcNAcb1-3Galb1-4(Fuca1-3)GlcNAcb-Sp0 | 6 | 3 | 58 |
| 293 | (6S)Galb1-4(6S)GlcNAcb-Sp0 | 6 | 4 | 71 |
| 443 | Gala1-3(Fuca1-2)Galb1-4GlcNAcb1-6(Gala1-3(Fuca1-2)Galb1-4GlcNAcb1-3)GalNAc-Sp14 | 6 | 2 | 33 |
| 573 | GlcNAcb1-3Galb1-4GlcNAcb1-3Galb1-4GlcNAcb1-3GalNAca-Sp14 | 6 | 5 | 83 |
| 184 | GlcNAcb1-3Galb1-4GlcNAcb1-3Galb1-4GlcNAcb-Sp0 | 6 | 1 | 23 |
| 301 | Galb1-4GlcNAcb1-6Galb1-4GlcNAcb-Sp0 | 6 | 1 | 10 |
| 313 | Neu5Aca2-3Galb1-4GlcNAcb1-6(Neu5Aca2-3Galb1-3)GalNAca-Sp14 | 6 | 1 | 23 |
| 316 | Neu5Aca2-8Neu5Acb-Sp17 | 6 | 2 | 31 |
| 407 | GalNAca1-3(Fuca1-2)Galb1-4GlcNAcb1-3GalNAca-Sp14 | 6 | 2 | 43 |
| 426 | GlcNAcb1-6(GlcNAcb1-2)Mana1-6(GlcNAcb1-4)(GlcNAcb1-2Mana1-3)Manb1-4GlcNAcb1-4GlcNAc-Sp21 | 6 | 8 | 144 |
| 478 | Neu5Aca2-3Galb1-3GlcNAcb1-2Mana1-6(GlcNAcb1-4)(Neu5Aca2-3Galb1-3GlcNAcb1-2Mana1-3)Manb1-4GlcNAcb1-4GlcNAc-Sp21 | 6 | 3 | 62 |

| | | | | |
|---|---|---|---|---|
| 567 | Neu5Aca2-3Galb1-4GlcNAcb1-3Galb1-4GlcNAcb1-3GalNAca-Sp14 | 6 | 4 | 67 |
| 335 | GlcNAca1-4Galb1-4GlcNAcb1-3Galb1-4Glcb-Sp0 | 5 | 3 | 59 |
| 244 | Neu5Aca2-3Galb1-3GlcNAcb1-3Galb1-4GlcNAcb-Sp0 | 5 | 5 | 112 |
| 473 | Neu5Aca2-6Galb1-4 GlcNAcb1-6(Neu5Aca2-6Galb1-4GlcNAcb1-3)GalNAca-Sp14 | 5 | 3 | 65 |
| 114 | Gala1-3Galb1-3GlcNAcb-Sp0 | 4 | 2 | 50 |
| 266 | Neu5Aca2-6Galb1-4GlcNAcb1-3Galb1-4(Fuca1-3)GlcNAcb1-3Galb1-4(Fuca1-3)GlcNAcb-Sp0 | 3 | 9 | 315 |
| 275 | Neu5Acb2-6Galb1-4GlcNAcb-Sp8 | 2 | 10 | 424 |
| 137 | Neu5Acb2-6(Galb1-3)GalNAca-Sp8 | 2 | 5 | 304 |
| 332 | GlcNAca1-4Galb1-4GlcNAcb1-3Galb1-4GlcNAcb1-3Galb1-4GlcNAcb-Sp0 | -6 | 17 | -280 |

**Figure S3: Glycans in microarray stratified by RFU for 034-10 4G02**

| Chart ID | Glycan | Average RFU | StDev | %CV |
|---|---|---|---|---|
| **39** | (6S)(4S)Galb1-4GlcNAcb-Sp0 | 29900 | 2320 | 8 |
| **40** | (4S)Galb1-4GlcNAcb-Sp8 | 26144 | 1822 | 7 |
| 459 | Neu5Aca2-6Galb1-4GlcNAcb1-6(Neu5Aca2-6Galb1-4GlcNAcb1-2)Mana1-6(GlcNAcb1-4)(Neu5Aca2-6Galb1-4GlcNAcb1-4(Neu5Aca2-6Galb1-4GlcNAcb1-2)Mana1-3)Manb1-4GlcNAcb1-4GlcNAcb-Sp21 | 1106 | 163 | 15 |
| 285 | Galb1-3GlcNAcb1-3Galb1-3GlcNAcb-Sp0 | 316 | 25 | 8 |
| 562 | Galb1-4GlcNAcb1-3Galb1-4GlcNAcb1-6(Galb1-4GlcNAcb1-3Galb1-4GlcNAb1-2)Mana1-6(Galb1-4GlcNAcb1-3Galb1-4GlcNAcb1-2Mana1-3)Manb1-4GlcNAcb1-4(Fuca1-6)GlcNAcb-Sp24 | 103 | 4 | 4 |
| 23 | 6S(3S)Galb1-4GlcNAcb-Sp0 | 82 | 19 | 23 |
| 545 | GlcNAcb1-3Galb1-4GlcNAcb1-6(GlcNAcb1-3Galb1-4GlcNAcb1-2)Mana1-6(GlcNAcb1-3Galb1-4GlcNAcb1-2Man a1-3)Manb1-4GlcNAcb1-4GlcNAc-Sp24 | 74 | 6 | 8 |
| 37 | (3S)Galb1-4GlcNAcb-Sp8 | 73 | 18 | 25 |
| 534 | Fuca1-2Galb1-4GlcNAcb1-3Galb1-4GlcNAcb1-2Mana1-6(Fuca1-2Galb1-4GlcNAcb1-3Galb1-4GlcNAcb1-2Mana1-3)Manb1-4GlcNAcb1-4GlcNAcb-Sp24 | 69 | 11 | 15 |
| 531 | GlcNAcb1-3Galb1-4GlcNAcb1-2Mana1-6(GlcNAcb1-3Galb1-4GlcNAcb1-2Mana1-3)Manb1-4GlcNAcb1-4GlcNAcb-Sp12 | 68 | 11 | 17 |
| 356 | Fuca1-2Galb1-4(Fuca1-3)GlcNAcb1-2Mana1-6(Fuca1-2Galb1-4(Fuca1-3)GlcNAcb1-2Mana1-3)Manb1-4GlcNAcb1-4GlcNAb-Sp20 | 67 | 4 | 6 |
| 88 | GlcNAcb1-3Galb1-3GalNAca-Sp8 | 63 | 3 | 5 |
| 336 | GlcNAca1-4Galb1-4GlcNAcb1-3Galb1-4(Fuca1-3)GlcNAcb1-3Galb1-4(Fuca1-3)GlcNAcb-Sp0 | 60 | 4 | 6 |
| 217 | (3S)Galb1-4(Fuca1-3)(6S)GlcNAcb-Sp8 | 59 | 5 | 9 |
| 465 | Fuca1-2Galb1-4(Fuca1-3)GlcNAcb1-2Mana1-6(Fuca1-2Galb1-4(Fuca1-3)GlcNAcb1-2Mana1-3)Manb1-4GlcNAcb1-4(Fuca1-6)GlcNAcb-Sp24 | 58 | 5 | 9 |
| 474 | Neu5Aca2-6Galb1-4GlcNAcb1-2Mana1-6(Neu5Aca2-6Galb1-4GlcNAcb1-2Mana1-3)Manb1-4GlcNAcb1-4(Fuca1-6)GlcNAcb-Sp24 | 57 | 11 | 19 |
| 44 | (6S)Galb1-4GlcNAcb-Sp8 | 56 | 8 | 14 |
| 24 | (3S)Galb1-4(Fuca1-3)(6S)Glc-Sp0 | 55 | 20 | 37 |
| 552 | Galb1-4GlcNAcb1-3Galb1-4GlcNAcb1-3Galb1-4GlcNAcb1-3Galb1-4GlcNAcb1-3Galb1-4GlcNAcb1-2Mana1-6(Galb1-4GlcNAcb1-3Galb1-4GlcNAcb1-3Galb1-4GlcNAcb1-3Galb1-4GlcNAcb1-3Galb1-4GlcNAcb1-2Mana1-3)Manb1-4GlcNAcb1-4GlcNAcb-Sp25 | 51 | 4 | 8 |
| 32 | (3S)Galb1-4(Fuca1-3)GlcNAc-Sp0 | 50 | 4 | 7 |
| 98 | GalNAcb1-4GlcNAcb-Sp0 | 50 | 51 | 102 |
| 557 | Galb1-3GlcNAcb1-3Galb1-4GlcNAcb1-3Galb1-4GlcNAcb1-6(Galb1-3GlcNAcb1-3Galb1-4GlcNAcb1-3Galb1-4GlcNAb1-2)Mana1-6(Galb1-3GlcNAcb1-3Galb1-4GlcNAcb1-3Galb1-4GlcNAcb1-2Mana1-3)Manb1-4GlcNAcb1-4(Fuca1-6)GlcNAcb-Sp24 | 50 | 11 | 21 |
| 372 | Neu5Aca2-3Galb1-4(Fuca1-3)GlcNAcb1-3GalNAca-Sp14 | 48 | 3 | 6 |
| 409 | Fuca1-2Galb1-4(Fuca1-3)GlcNAcb1-3GalNAca-Sp14 | 48 | 7 | 15 |
| 536 | GlcNAcb1-3Galb1-4GlcNAcb1-3Galb1-4GlcNAcb1-2Mana1-6(GlcNAcb1-3Galb1-4GlcNAcb1-3Galb1-4GlcNAcb1-2Mana1-3)Manb1-4GlcNAcb1-4GlcNAcb-Sp25 | 47 | 8 | 17 |
| 466 | Fuca1-2Galb1-3(Fuca1-4)GlcNAcb1-2Mana1-6(Fuca1-2Galb1-3(Fuca1-4)GlcNAcb1-2Mana1-3)Manb1-4GlcNAcb1-4(Fuca1-6)GlcNAcb1-4(Fuca1-6)GlcNAcb-Sp19 | 46 | 10 | 22 |
| 331 | Neu5Aca2-3Galb1-4(Fuca1-3)GlcNAcb1-6(Neu5Aca2-3Galb1-3)GalNAc-Sp14 | 46 | 5 | 11 |
| 437 | (6S)Galb1-3(6S)GlcNAc-Sp0 | 45 | 6 | 13 |
| 535 | GlcNAcb1-3Galb1-4GlcNAcb1-3Galb1-4GlcNAcb1-2Mana1-6(GlcNAcb1-3Galb1-4GlcNAcb1-3Galb1-4GlcNAcb1-2Mana1-3)Manb1-4GlcNAcb1-4GlcNAcb-Sp12 | 44 | 3 | 7 |
| 36 | (3S)Galb1-4GlcNAcb-Sp0 | 43 | 6 | 15 |
| 477 | Galb1-4GlcNAcb1-6(Galb1-4GlcNAcb1-2)Mana1-6(Galb1-4GlcNAcb1-2Mana1-3)Manb1-4GlcNAcb1-4(Fuca1-6)GlcNAcb-Sp24 | 43 | 15 | 34 |
| 582 | Neu5Aca2-3Galb1-4GlcNAcb1-3Galb1-4GlcNAcb1-3Galb1-4GlcNAcb1-2Mana1-6(Neu5Aca2-3Galb1-4GlcNAcb1-3Galb1-4GlcNAcb1-3Galb1-4GlcNAcb1-2Mana1-3)Manb1-4GlcNAcb1-4GlcNAcb-Sp12 | 43 | 1 | 2 |
| 354 | Fuca1-2Galb1-3GlcNAcb1-2Mana1-6(Fuca1-2Galb1-3GlcNAcb1-2Mana1-3)Manb1-4GlcNAcb1-4GlcNAcb-Sp20 | 43 | 9 | 20 |
| 547 | Gala1-3Galb1-4GlcNAcb1-2Mana1-6(Gala1-3Galb1-4GlcNAcb1-2Mana1-3)Manb1-4GlcNAcb1-4GlcNAc-Sp24 | 43 | 10 | 24 |

| | | | | |
|---|---|---|---|---|
| 389 | Gala1-3Galb1-3(Fuca1-4)GlcNAcb1-2Mana1-6(Gala1-3Galb1-3(Fuca1-4)GlcNAcb1-2Mana1-3)Manb1-4GlcNAcb1-4GlcNAc-Sp19 | 43 | 8 | 18 |
| 558 | Galb1-3GlcNAcb1-3Galb1-4GlcNAcb1-6(Galb1-3GlcNAcb1-3Galb1-4GlcNAb1-2)Mana1-6(Galb1-3GlcNAcb1-3Galb1-4GlcNAcb1-2Mana1-3)Manb1-4GlcNAcb1-4(Fuca1-6)GlcNAcb-Sp24 | 43 | 6 | 15 |
| 84 | (3S)Galb1-4(Fuca1-3)Glcb-Sp0 | 42 | 8 | 18 |
| 22 | 6S(3S)Galb1-4(6S)GlcNAcb-Sp0 | 41 | 10 | 23 |
| 216 | Neu5Aca2-3Galb1-4GlcNAcb1-3Galb1-4(Fuca1-3)GlcNAcb-Sp0 | 40 | 9 | 22 |
| 413 | Fuca1-2Galb1-4GlcNAcb1-2Mana1-6(Fuca1-2Galb1-4GlcNAcb1-2Mana1-3)Manb1-4GlcNAcb1-4(Fuca1-6)GlcNAcb-Sp22 | 40 | 5 | 14 |
| 438 | Fuca1-2Galb1-4 GlcNAcb1-2Mana1-6(Fuca1-2Galb1-4GlcNAcb1-2(Fuca1-2Galb1-4GlcNAcb1-4)Mana1-3)Manb1-4GlcNAcb1-4GlcNAcb-Sp12 | 40 | 6 | 16 |
| 26 | (3S)Galb1-4(6S)Glcb-Sp0 | 40 | 5 | 12 |
| 249 | Neu5Aca2-3Galb1-4(Fuca1-3)GlcNAcb1-3Galb1-4(Fuca1-3)GlcNAcb1-3Galb1-4(Fuca1-3)GlcNAcb-Sp0 | 40 | 4 | 9 |
| 220 | Fuca1-2(6S)Galb1-4(6S)Glcb-Sp0 | 39 | 8 | 20 |
| 359 | Fuca1-4(Galb1-3)GlcNAcb1-2Mana1-6(Fuca1-4(Galb1-3)GlcNAcb1-2Mana1-3)Manb1-4GlcNAcb1-4(Fuca1-6)GlcNAcb-Sp22 | 39 | 8 | 21 |
| 492 | Fuca1-2Galb1-3(6S)GlcNAcb-Sp0 | 38 | 24 | 63 |
| 537 | Galb1-4GlcNAcb1-3Galb1-4GlcNAcb1-3Galb1-4GlcNAcb1-2Mana1-6(Galb1-4GlcNAcb1-3Galb1-4GlcNAcb1-3Galb1-4GlcNAcb1-2Mana1-3)Manb1-4GlcNAcb1-4GlcNAcb-Sp12 | 38 | 6 | 16 |
| 97 | GalNAcb1-4(Fuca1-3)GlcNAcb-Sp0 | 38 | 10 | 28 |
| 546 | Galb1-4GlcNAcb1-3Galb1-4GlcNAcb1-6(Galb1-4GlcNAcb1-3Galb1-4GlcNAcb1-2)Mana1-6(Galb1-4GlcNAcb1-3Galb1-4GlcNAcb1-2Mana1-3)Mana1-4GlcNAcb1-4GlcNAc-Sp24 | 38 | 7 | 17 |
| 57 | Neu5Aca2-6Galb1-4GlcNAcb1-2Mana1-6(Neu5Aca2-6Galb1-4GlcNAcb1-2Mana1-3)Manb1-4GlcNAcb1-4GlcNAcb-Sp24 | 37 | 1 | 3 |
| 325 | Neu5Aca2-3Galb1-3(Fuca1-4)GlcNAcb1-3Galb1-3(Fuca1-4)GlcNAcb-Sp0 | 37 | 10 | 28 |
| 366 | Gala1-3Galb1-4(Fuca1-3)GlcNAcb1-2Mana1-6(Gala1-3Galb1-4(Fuca1-3)GlcNAcb1-2Mana1-3)Manb1-4GlcNAcb1-4GlcNAcb-Sp20 | 36 | 3 | 8 |
| 3 | Mana-Sp8 | 36 | 9 | 25 |
| 439 | Fuca1-2Galb1-4(Fuca1-3)GlcNAcb1-2Mana1-6(Fuca1-2Galb1-4(Fuca1-3)GlcNAcb1-4(Fuca1-2Galb1-4(Fuca1-3)GlcNAcb1-2)Mana1-3)Manb1-4GlcNAcb1-4GlcNAcb-Sp12 | 36 | 15 | 43 |
| 538 | Galb1-3GlcNAcb1-3Galb1-4GlcNAcb1-2Mana1-6(Galb1-3GlcNAcb1-3Galb1-4GlcNAcb1-2Mana1-3)Manb1-4GlcNAcb1-4GlcNAc-Sp25 | 36 | 8 | 24 |
| 9 | Neu5Aca-Sp8 | 35 | 7 | 19 |
| 218 | Fuca1-2(6S)Galb1-4GlcNAcb-Sp0 | 35 | 4 | 10 |
| 467 | GlcNAcb1-6(GlcNAcb1-2)Mana1-6(GlcNAcb1-2Mana1-3)Manb1-4GlcNAcb1-4(Fuca1-6)GlcNAcb-Sp24 | 35 | 6 | 18 |
| 475 | Neu5Aca2-3Galb1-4GlcNAcb1-2Mana1-6(Neu5Aca2-3Galb1-4GlcNAcb1-2Mana1-3)Manb1-4GlcNAcb1-4(Fuca1-6)GlcNAcb-Sp24 | 35 | 9 | 27 |
| 273 | Galb1-3(Fuca1-4)GlcNAcb1-3Galb1-3(Fuca1-4)GlcNAcb-Sp0 | 35 | 2 | 6 |
| 412 | Galb1-4(Fuca1-3)GlcNAcb1-2Mana1-6(Galb1-4(Fuca1-3)GlcNAcb1-2Mana1-3)Manb1-4GlcNAcb1-4(Fuca1-6)GlcNAcb-Sp22 | 34 | 5 | 14 |
| 99 | GalNAcb1-4GlcNAcb-Sp8 | 34 | 13 | 40 |
| 397 | Gala1-4Galb1-4GlcNAcb1-2Mana1-6(Gala1-4Galb1-4GlcNAcb1-2Mana1-3)Manb1-4GlcNAcb1-4GlcNAcb-Sp24 | 33 | 5 | 14 |
| 103 | Gala1-3(Fuca1-2)Galb1-4(Fuca1-3)GlcNAcb-Sp0 | 33 | 2 | 6 |
| 388 | Gala1-3Galb1-3GlcNAcb1-2Mana1-6(Gala1-3Galb1-3GlcNAcb1-2Mana1-3)Manb1-4GlcNAcb1-4GlcNAc-Sp19 | 33 | 8 | 25 |
| 355 | Fuca1-2Galb1-4GlcNAcb1-2Mana1-6(Fuca1-2Galb1-4GlcNAcb1-2Mana1-3)Manb1-4GlcNAcb1-4GlcNAcb-Sp20 | 32 | 4 | 12 |
| 560 | Galb1-4GlcNAcb1-3Galb1-4GlcNAcb1-2Mana1-6(Galb1-4GlcNAcb1-3Galb1-4GlcNAcb1-2Mana1-3)Manb1-4GlcNAcb1-4(Fuca1-6)GlcNAcb-Sp24 | 32 | 2 | 5 |
| 108 | Gala1-3(Fuca1-2)Galb-Sp18 | 32 | 7 | 21 |
| 253 | Neu5Aca2-3Galb1-4(Fuca1-3)GlcNAcb1-3Galb1-4GlcNAcb-Sp8 | 32 | 2 | 5 |
| 530 | Galb1-3GalNAcb1-3Gal-Sp21 | 32 | 3 | 8 |
| 581 | Neu5Aca2-6Galb1-4GlcNAcb1-3Galb1-4GlcNAcb1-3Galb1-4GlcNAcb1-2Mana1-6(Neu5Aca2-6Galb1-4GlcNAcb1-3Galb1-4GlcNAcb1-3Galb1-4GlcNAcb1-2Mana1-3)Manb1-4GlcNAcb1-4GlcNAcb-Sp12 | 31 | 5 | 15 |

| | | | | |
|---|---|---|---|---|
| 295 | Neu5Aca2-3Galb1-4(Fuca1-3)GlcNAcb1-6(Galb1-3)GalNAca-Sp14 | 31 | 2 | 6 |
| 449 | Neu5Aca2-6Galb1-4GlcNAcb1-6(Fuca1-2Galb1-3GlcNAcb1-3)Galb1-4Glc-Sp21 | 30 | 7 | 22 |
| 577 | Neu5Aca2-6Galb1-4GlcNAcb1-6(Galb1-3)GalNAca-Sp14 | 30 | 10 | 34 |
| 497 | Fuca1-2Galb1-3GlcNAcb1-6(Fuca1-2Galb1-3GlcNAcb1-3)GalNAca-Sp14 | 30 | 6 | 19 |
| 100 | Gala1-2Galb-Sp8 | 29 | 2 | 8 |
| 493 | Fuca1-2(6S)Galb1-3(6S)GlcNAcb-Sp0 | 29 | 5 | 16 |
| 525 | Fuca1-4(Galb1-3)GlcNAcb1-2 Mana-Sp0 | 29 | 12 | 41 |
| 237 | Neu5Aca2-3Galb1-3(Fuca1-4)GlcNAcb1-3Galb1-4(Fuca1-3)GlcNAcb-Sp0 | 29 | 2 | 5 |
| 239 | Neu5Aca2-3Galb1-3(6S)GalNAca-Sp8 | 29 | 4 | 14 |
| 450 | GalNAca1-3(Fuca1-2)Galb1-3GlcNAcb1-2Mana1-6(GalNAca1-3(Fuca1-2)Galb1-3GlcNAcb1-2Mana1-3)Manb1-4GlcNAcb1-4(Fuca1-6)GlcNAcb-Sp22 | 29 | 4 | 12 |
| 248 | Neu5Aca2-3Galb1-4(Fuca1-3)(6S)GlcNAcb-Sp8 | 29 | 10 | 36 |
| 348 | Galb1-3GlcNAcb1-2Mana1-6(Galb1-3GlcNAcb1-2Mana1-3)Manb1-4GlcNAcb1-4(Fuca1-6)GlcNAcb-Sp22 | 29 | 6 | 21 |
| 350 | KDNa2-3Galb1-4(Fuca1-3)GlcNAc-Sp0 | 29 | 7 | 23 |
| 378 | Galb1-4GlcNAcb1-6(Fuca1-4(Fuca1-2Galb1-3)GlcNAcb1-3)Galb1-4Glc-Sp21 | 29 | 3 | 9 |
| 445 | Neu5Aca2-8Neu5Aca2-3Galb1-3GalNAcb1-4(Neu5Aca2-8Neu5Aca2-3)Galb1-4Glcb-Sp0 | 29 | 8 | 29 |
| 50 | Mana1-6(Mana1-3)Manb1-4GlcNAcb1-4GlcNAcb-Sp12 | 28 | 1 | 3 |
| 121 | Gala1-4Galb1-4GlcNAcb-Sp8 | 28 | 9 | 34 |
| 60 | Fuca1-2Galb1-3(Fuca1-4)GlcNAcb-Sp8 | 28 | 1 | 5 |
| 318 | Neu5Gcb2-6Galb1-4GlcNAc-Sp8 | 28 | 3 | 12 |
| 70 | Fuca1-2Galb1-4(Fuca1-3)GlcNAcb1-3Galb1-4(Fuca1-3)GlcNAcb1-3Galb1-4(Fuca1-3)GlcNAcb-Sp0 | 28 | 2 | 8 |
| 286 | Galb1-4(Fuca1-3)(6S)GlcNAcb-Sp0 | 28 | 1 | 5 |
| 440 | Galb1-4(Fuca1-3)GlcNAcb1-6GalNAc-Sp14 | 28 | 8 | 28 |
| 109 | Gala1-4(Gala1-3)Galb1-4GlcNAcb-Sp8 | 27 | 11 | 42 |
| 367 | GalNAca1-3(Fuca1-2)Galb1-3GlcNAcb1-2Mana1-6(GalNAca1-3(Fuca1-2)Galb1-3GlcNAcb1-2Mana1-3)Manb1-4GlcNAcb1-4GlcNAcb-Sp20 | 27 | 5 | 18 |
| 110 | Gala1-3GalNAca-Sp8 | 27 | 6 | 22 |
| 245 | Fuca1-2(6S)Galb1-4Glcb-Sp0 | 27 | 3 | 9 |
| 228 | GalNAcb1-4(Neu5Aca2-3)Galb1-4GlcNAcb-Sp0 | 26 | 2 | 7 |
| 373 | GalNAcb1-4GlcNAcb1-2Mana1-6(GalNAcb1-4GlcNAcb1-2Mana1-3)Manb1-4GlcNAcb1-4GlcNAc-Sp12 | 26 | 8 | 30 |
| 11 | Neu5Acb-Sp8 | 26 | 4 | 16 |
| 208 | Mana1-2Mana1-6(Mana1-2Mana1-3)Mana-Sp9 | 26 | 3 | 11 |
| 209 | Mana1-2Mana1-3Mana-Sp9 | 26 | 3 | 12 |
| 346 | GlcNAcb1-2Mana1-6(GlcNAcb1-2Mana1-3)Manb1-4GlcNAcb1-4(Fuca1-6)GlcNAcb-Sp22 | 26 | 9 | 33 |
| 379 | Galb1-4(Fuca1-3)GlcNAcb1-6(Fuca1-4(Fuca1-2Galb1-3)GlcNAcb1-3)Galb1-4Glc-Sp21 | 26 | 3 | 10 |
| 102 | Gala1-3(Fuca1-2)Galb1-3GlcNAcb-Sp8 | 26 | 5 | 18 |
| 158 | Galb1-4GalNAcb1-3(Fuca1-2)Galb1-4GlcNAcb-Sp8 | 26 | 2 | 9 |
| 468 | Galb1-3GlcNAcb1-2Mana1-6(GlcNAcb1-4)(Galb1-3GlcNAcb1-2Mana1-3)Manb1-4GlcNAcb1-4GlcNAcb-Sp21 | 26 | 7 | 28 |
| 170 | Galb1-4GlcNAcb-Sp23 | 25 | 2 | 6 |
| 13 | Glcb-Sp8 | 25 | 5 | 22 |
| 85 | GalNAca1-3(Fuca1-2)Galb1-4GlcNAcb-Sp0 | 25 | 1 | 6 |
| 124 | Gala1-6Glcb-Sp8 | 25 | 4 | 15 |
| 157 | Galb1-4GalNAca1-3(Fuca1-2)Galb1-4GlcNAcb-Sp8 | 25 | 2 | 10 |
| 263 | Neu5Aca2-6Galb1-4(6S)GlcNAcb-Sp8 | 25 | 6 | 26 |
| 401 | GalNAcb1-3Gala1-6Galb1-4Glcb-Sp8 | 25 | 2 | 9 |
| 578 | Neu5Aca2-3Galb1-4GlcNAcb1-3Galb1-4GlcNAcb1-2Mana1-6(Neu5Aca2-3Galb1-4GlcNAcb1-3Galb1-4GlcNAcb1-2Mana1-3)Manb1-4GlcNAcb1-4GlcNAcb-Sp12 | 25 | 7 | 27 |
| 95 | GalNAcb1-3(Fuca1-2)Galb-Sp8 | 25 | 1 | 2 |

| | | | | |
|---|---|---|---|---|
| 194 | GlcNAcb1-6Galb1-4GlcNAcb-Sp8 | 25 | 2 | 9 |
| 153 | Galb1-4(Fuca1-3)GlcNAcb1-3Galb1-4(Fuca1-3)GlcNAcb-Sp0 | 25 | 1 | 5 |
| 207 | Mana1-2Mana1-2Mana1-3Mana-Sp9 | 25 | 4 | 16 |
| 275 | Neu5Acb2-6Galb1-4GlcNAcb-Sp8 | 25 | 3 | 14 |
| 364 | GalNAca1-3(Fuca1-2)Galb1-4GlcNAcb1-2Mana1-6(GalNAca1-3(Fuca1-2)Galb1-4GlcNAcb1-2Mana1-3)Manb1-4GlcNAcb1-4GlcNAcb-Sp20 | 25 | 1 | 4 |
| 583 | Neu5Aca2-6Galb1-4GlcNAcb1-3Galb1-4GlcNAcb1-2Mana1-6(Neu5Aca2-6Galb1-4GlcNAcb1-3Galb1-4GlcNAcb1-2Mana1-3)Manb1-4GlcNAcb1-4GlcNAcb-Sp12 | 24 | 5 | 21 |
| 14 | Manb-Sp8 | 24 | 6 | 27 |
| 15 | GalNAcb-Sp8 | 24 | 7 | 29 |
| 53 | GlcNAcb1-2Mana1-6(GlcNAcb1-2Mana1-3)Manb1-4GlcNAcb1-4GlcNAcb-Sp13 | 24 | 4 | 17 |
| 21 | GlcNAcb1-6(GlcNAcb1-4)(GlcNAcb1-3)GlcNAc-Sp8 | 24 | 10 | 40 |
| 101 | Gala1-3(Fuca1-2)Galb1-3GlcNAcb-Sp0 | 24 | 2 | 7 |
| 123 | Gala1-4GlcNAcb-Sp8 | 24 | 3 | 13 |
| 161 | Galb1-4GlcNAcb1-3Galb1-4(Fuca1-3)GlcNAcb1-3Galb1-4(Fuca1-3)GlcNAcb-Sp0 | 24 | 1 | 5 |
| 240 | Neu5Aca2-6(Neu5Aca2-3Galb1-3)GalNAca-Sp8 | 24 | 1 | 4 |
| 16 | GlcNAcb-Sp0 | 24 | 10 | 44 |
| 135 | Neu5Aca2-6(Galb1-3)GalNAca-Sp8 | 24 | 1 | 4 |
| 261 | Neu5Aca2-6GalNAca-Sp8 | 23 | 2 | 8 |
| 264 | Neu5Aca2-6Galb1-4GlcNAcb-Sp0 | 23 | 7 | 28 |
| 319 | Galb1-3GlcNAcb1-2Mana1-6(Galb1-3GlcNAcb1-2Mana1-3)Manb1-4GlcNAcb1-4GlcNAcb-Sp19 | 23 | 3 | 11 |
| 411 | GalNAca1-3(Fuca1-2)Galb1-4(Fuca1-3)GlcNAcb1-3GalNAc-Sp14 | 23 | 8 | 34 |
| 27 | (3S)Galb1-4(6S)Glcb-Sp8 | 23 | 2 | 8 |
| 188 | GlcNAcb1-4Galb1-4GlcNAcb-Sp8 | 23 | 4 | 18 |
| 293 | (6S)Galb1-4(6S)GlcNAcb-Sp0 | 23 | 3 | 13 |
| 476 | Mana1-6(Mana1-3)Manb1-4GlcNAcb1-4(Fuca1-6)GlcNAcb-Sp19 | 23 | 1 | 6 |
| 488 | Galb1-4(Fuca1-3)GlcNAcb1-2Mana-Sp0 | 23 | 12 | 51 |
| 96 | GalNAcb1-3Gala1-4Galb1-4GlcNAcb-Sp0 | 23 | 5 | 20 |
| 369 | Fuca1-4(Fuca1-2Galb1-3)GlcNAcb1-2Mana1-3(Fuca1-4(Fuca1-2Galb1-3)GlcNAcb1-2Mana1-3)Manb1-4GlcNAcb1-4GlcNAcb-Sp19 | 23 | 4 | 18 |
| 38 | (3S)Galb-Sp8 | 23 | 2 | 9 |
| 94 | GalNAcb1-3GalNAca-Sp8 | 23 | 2 | 8 |
| 141 | Galb1-3GalNAca-Sp16 | 23 | 2 | 9 |
| 180 | GlcNAcb1-3GalNAca-Sp14 | 23 | 1 | 4 |
| 368 | Galα1-3(Fucα1-2)Galβ1-3GlcNAcβ1-2Manα1-6(Galα1-3(Fucα1-2)Galβ1-3GlcNAcβ1-2Manα1-3)Manβ1-4GlcNAcβ1-4GlcNAcβ-Sp20 | 23 | 1 | 3 |
| 414 | GlcNAcb1-2(GlcNAcb1-6)Mana1-6(GlcNAcb1-2Mana1-3)Manb1-4GlcNAcb1-4GlcNAcb-Sp19 | 23 | 2 | 8 |
| 104 | Gala1-3(Fuca1-2)Galb1-4(Fuca1-3)GlcNAcb-Sp8 | 22 | 3 | 12 |
| 112 | Gala1-3GalNAcb-Sp8 | 22 | 2 | 8 |
| 352 | KDNa2-3Galb1-4Glc-Sp0 | 22 | 9 | 41 |
| 386 | GalNAcb1-4(Neu5Aca2-3)Galb1-4GlcNAcb1-3GalNAca-Sp14 | 22 | 1 | 4 |
| 451 | Galb1-4GlcNAcb1-6(Galb1-4GlcNAcb1-2)Mana1-6(Galb1-4GlcNAcb1-2Mana1-3)Manb1-4GlcNAcb1-4GlcNAcb-Sp19 | 22 | 5 | 23 |
| 2 | Glca-Sp8 | 22 | 9 | 40 |
| 18 | GlcN(Gc)b-Sp8 | 22 | 8 | 37 |
| 19 | Galb1-4GlcNAcb1-6(Galb1-4GlcNAcb1-3)GalNAca-Sp8 | 22 | 9 | 39 |
| 250 | Neu5Aca2-3Galb1-4(Fuca1-3)GlcNAcb-Sp0 | 22 | 1 | 4 |
| 486 | Fuca1-2Galb1-4GlcNAcb1-6GalNAca-Sp14 | 22 | 7 | 34 |
| 61 | Fuca1-2Galb1-3GalNAca-Sp8 | 22 | 2 | 7 |
| 72 | Fuca1-2Galb1-4(Fuca1-3)GlcNAcb-Sp8 | 22 | 2 | 7 |
| 516 | Galb1-3GlcNAcb1-2Mana-Sp0 | 22 | 7 | 32 |
| 566 | Galb1-4GlcNAcb1-3Galb1-4GlcNAcb1-6(Galb1-4GlcNAcb1-3Galb1-4GlcNAcb1-3)GalNAca-Sp14 | 22 | 3 | 11 |

| | | | | |
|---|---|---|---|---|
| 214 | Mana1-6(Mana1-3)Mana1-6(Mana1-3)Manb1-4GlcNAcb1-4GlcNAcb-Sp12 | 22 | 3 | 14 |
| 47 | (6S)GlcNAcb-Sp8 | 21 | 6 | 26 |
| 193 | GlcNAcb1-6GalNAca-Sp14 | 21 | 6 | 28 |
| 381 | Galb1-4GlcNAcb1-6(Galb1-4GlcNAcb1-2)Mana1-6(Galb1-4GlcNAcb1-4(Galb1-4GlcNAcb1-2)Mana1-3)Manb1-4GlcNAcb1-4GlcNAcb-Sp21 | 21 | 1 | 2 |
| 390 | GlcNAcb1-2Mana1-6(Galb1-4GlcNAcb1-2Mana1-3)Manb1-4GlcNAcb1-4GlcNAc-Sp12 | 21 | 5 | 22 |
| 501 | Galb1-3GlcNAca1-3Galb1-4GlcNAcb-Sp8 | 21 | 2 | 7 |
| 8 | Rhaa-Sp8 | 21 | 3 | 12 |
| 73 | Fuca1-2Galb1-4GlcNAcb1-3Galb1-4GlcNAcb-Sp0 | 21 | 1 | 7 |
| 181 | GlcNAcb1-3Galb-Sp8 | 21 | 2 | 11 |
| 195 | Glca1-4Glcb-Sp8 | 21 | 3 | 13 |
| 338 | GlcNAca1-4Galb1-3GalNAc-Sp14 | 21 | 2 | 10 |
| 357 | Gala1-3Galb1-4GlcNAcb1-2Mana1-6(Gala1-3Galb1-4GlcNAcb1-2Mana1-3)Manb1-4GlcNAcb1-4GlcNAcb-Sp20 | 21 | 2 | 12 |
| 83 | GalNAca1-3(Fuca1-2)Galb1-4(Fuca1-3)GlcNAcb-Sp0 | 21 | 2 | 10 |
| 196 | Glca1-4Glca-Sp8 | 21 | 2 | 11 |
| 363 | Galb1-4GlcNAcb1-2Mana1-6(Galb1-4GlcNAcb1-4(Galb1-4GlcNAcb1-2)Mana1-3)Manb1-4GlcNAcb1-4GlcNAc-Sp21 | 21 | 2 | 11 |
| 380 | Galb1-3GlcNAcb1-3Galb1-4(Fuca1-3)GlcNAcb1-6(Galb1-3GlcNAcb1-3)Galb1-4Glc-Sp21 | 21 | 4 | 18 |
| 137 | Neu5Acb2-6(Galb1-3)GalNAca-Sp8 | 21 | 2 | 12 |
| 206 | KDNa2-3Galb1-4GlcNAcb-Sp0 | 21 | 5 | 27 |
| 280 | Neu5Gca2-3Galb1-4Glcb-Sp0 | 21 | 3 | 12 |
| 526 | Neu5Aca2-3Galb1-4(Fuca1-3)GlcNAcb1-2Mana-Sp0 | 21 | 8 | 38 |
| 539 | Neu5Gca2-8Neu5Gca2-3Galb1-4GlcNAc-Sp0 | 21 | 7 | 34 |
| 553 | GlcNAb1-3Galb1-3GalNAc-Sp14 | 21 | 2 | 12 |
| 17 | GlcNAcb-Sp8 | 20 | 5 | 26 |
| 105 | Gala1-3(Fuca1-2)Galb1-4GlcNAc-Sp0 | 20 | 3 | 13 |
| 119 | Gala1-4(Fuca1-2)Galb1-4GlcNAcb-Sp8 | 20 | 1 | 2 |
| 334 | GlcNAca1-4Galb1-3GlcNAcb-Sp0 | 20 | 3 | 13 |
| 347 | Galb1-4GlcNAcb1-2Mana1-6(Galb1-4GlcNAcb1-2Mana1-3)Manb1-4GlcNAcb1-4(Fuca1-6)GlcNAcb-Sp22 | 20 | 1 | 6 |
| 420 | Gala1-3(Fuca1-2)Galb1-4GlcNAcb1-2Mana1-6(Gala1-3(Fuca1-2)Galb1-4GlcNAcb1-2Mana1-3)Manb1-4GlcNAcb1-4(Fuca1-6)GlcNAcb-Sp22 | 20 | 5 | 26 |
| 461 | Gala1-3(Fuca1-2)Galb1-3GalNAcb-Sp8 | 20 | 2 | 9 |
| 30 | (3S)Galb1-3GlcNAcb-Sp0 | 20 | 3 | 17 |
| 169 | Galb1-4GlcNAcb-Sp8 | 20 | 1 | 7 |
| 266 | Neu5Aca2-6Galb1-4GlcNAcb1-3Galb1-4(Fuca1-3)GlcNAcb1-3Galb1-4(Fuca1-3)GlcNAcb-Sp0 | 20 | 1 | 7 |
| 452 | Neu5Aca2-3Galb1-4GlcNAcb1-2Mana1-6(GlcNAcb1-4)(Neu5Aca2-3Galb1-4GlcNAcb1-2Mana1-3)Manb1-4GlcNAcb1-4GlcNAcb-Sp21 | 20 | 3 | 15 |
| 35 | (3S)Galb1-4(6S)GlcNAcb-Sp8 | 20 | 2 | 8 |
| 45 | (6S)Galb1-4(6S)Glcb-Sp8 | 20 | 3 | 15 |
| 69 | Fuca1-2Galb1-4(Fuca1-3)GlcNAcb1-3Galb1-4(Fuca1-3)GlcNAcb-Sp0 | 20 | 3 | 13 |
| 74 | Fuca1-2Galb1-4GlcNAcb1-3Galb1-4GlcNAcb1-3Galb1-4GlcNAcb-Sp0 | 20 | 1 | 6 |
| 179 | GlcNAcb1-3GalNAca-Sp8 | 20 | 2 | 11 |
| 187 | GlcNAcb1-6(GlcNAcb1-4)GalNAca-Sp8 | 20 | 10 | 49 |
| 131 | Galb1-4GlcNAcb1-6GalNAca-Sp8 | 20 | 2 | 9 |
| 133 | GlcNAcb1-6(Galb1-3)GalNAca-Sp8 | 20 | 4 | 19 |
| 134 | GlcNAcb1-6(Galb1-3)GalNAca-Sp14 | 20 | 2 | 9 |
| 272 | Neu5Aca2-8Neu5Aca2-3Galb1-4Glcb-Sp0 | 20 | 5 | 24 |
| 303 | GlcAb1-3GlcNAcb-Sp8 | 20 | 1 | 5 |
| 333 | GlcNAca1-4Galb1-4GlcNAcb-Sp0 | 20 | 3 | 16 |
| 435 | GalNAcb1-6GalNAcb-Sp8 | 20 | 12 | 60 |
| 576 | Neu5Aca2-6Galb1-4GlcNAcb1-3Galb1-4GlcNAcb1-6(Galb1-3)GalNAca-Sp14 | 20 | 1 | 5 |

| | | | | |
|---|---|---|---|---|
| 584 | GlcNAcb1-3Fuca-Sp21 | 20 | 2 | 9 |
| 48 | Neu5,9Ac2a-Sp8 | 19 | 3 | 16 |
| 113 | Gala1-3Galb1-4(Fuca1-3)GlcNAcb-Sp8 | 19 | 4 | 20 |
| 126 | Galb1-3(Fuca1-4)GlcNAcb1-3Galb1-4(Fuca1-3)GlcNAcb-Sp0 | 19 | 3 | 13 |
| 159 | Galb1-4GlcNAcb1-3GalNAca-Sp8 | 19 | 2 | 12 |
| 167 | Galb1-4GlcNAcb1-6(Galb1-3)GalNAc-Sp14 | 19 | 2 | 9 |
| 223 | GalNAcb1-4(Neu5Aca2-8Neu5Aca2-8Neu5Aca2-8Neu5Aca2-3)Galb1-4Glcb-Sp0 | 19 | 2 | 9 |
| 241 | Neu5Aca2-6(Neu5Aca2-3Galb1-3)GalNAca-Sp14 | 19 | 2 | 12 |
| 274 | Neu5Acb2-6GalNAca-Sp8 | 19 | 6 | 32 |
| 300 | Galb1-4GlcNAca1-6Galb1-4GlcNAcb-Sp0 | 19 | 4 | 19 |
| 212 | Mana1-2Mana1-2Mana1-6(Mana1-3)Mana-Sp9 | 19 | 2 | 11 |
| 242 | Neu5Aca2-3Galb-Sp8 | 19 | 3 | 15 |
| 246 | Neu5Aca2-3Galb1-3GlcNAcb-Sp0 | 19 | 0 | 0 |
| 255 | Neu5Aca2-3Galb1-4GlcNAcb-Sp0 | 19 | 1 | 6 |
| 288 | Galb1-4(Fuca1-3)GlcNAcb1-3Galb1-3(Fuca1-4)GlcNAcb-Sp0 | 19 | 3 | 18 |
| 349 | (6S)GlcNAcb1-3Galb1-4GlcNAcb-Sp0 | 19 | 2 | 13 |
| 404 | Galb1-3GalNAcb1-4(Neu5Aca2-8Neu5Aca2-3)Galb1-4Glcb-Sp0 | 19 | 8 | 42 |
| 472 | Neu5Aca2-6Galb1-4GlcNAcb1-6GalNAca-Sp14 | 19 | 6 | 32 |
| 574 | Galb1-4GlcNAcb1-3Galb1-3GalNAca-Sp14 | 19 | 5 | 29 |
| 1 | Gala-Sp8 | 19 | 1 | 7 |
| 6 | Fuca-Sp8 | 19 | 6 | 29 |
| 52 | GlcNAcb1-2Mana1-6(GlcNAcb1-2Mana1-3)Manb1-4GlcNAcb1-4GlcNAcb-Sp12 | 19 | 2 | 9 |
| 65 | Fuca1-2Galb1-3GlcNAcb1-3Galb1-4Glcb-Sp8 | 19 | 1 | 3 |
| 111 | Gala1-3GalNAca-Sp16 | 19 | 3 | 16 |
| 145 | Galb1-3GalNAcb1-4Galb1-4Glcb-Sp8 | 19 | 3 | 13 |
| 287 | Galb1-4(Fuca1-3)(6S)Glcb-Sp0 | 19 | 1 | 3 |
| 320 | Neu5Aca2-3Galb1-4GlcNAcb1-2Mana1-6(Neu5Aca2-3Galb1-4GlcNAcb1-2Mana1-3)Manb1-4GlcNAcb1-4GlcNAcb-Sp12 | 19 | 2 | 9 |
| 344 | Galb1-4GlcNAcb1-2Mana1-6Manb1-4GlcNAcb1-4GlcNAc-Sp12 | 19 | 3 | 13 |
| 361 | Neu5Aca2-6GlcNAcb1-4GlcNAcb1-4GlcNAc-Sp21 | 19 | 2 | 12 |
| 362 | Galb1-4(Fuca1-3)GlcNAcb1-6(Fuca1-2Galb1-4GlcNAcb1-3)Galb1-4Glc-Sp21 | 19 | 2 | 8 |
| 436 | (6S)Galb1-3GlcNAcb-Sp0 | 19 | 6 | 34 |
| 564 | Galb1-4GlcNAcb1-3Galb1-4GlcNAcb1-3GalNAca-Sp14 | 19 | 5 | 24 |
| 29 | (3S)Galb1-3GalNAca-Sp8 | 19 | 1 | 7 |
| 186 | GlcNAcb1-4-MDPLys | 19 | 2 | 9 |
| 262 | Neu5Aca2-6GalNAcb1-4GlcNAcb-Sp0 | 19 | 3 | 14 |
| 265 | Neu5Aca2-6Galb1-4GlcNAcb-Sp8 | 19 | 3 | 17 |
| 271 | Neu5Aca2-8Neu5Aca-Sp8 | 19 | 1 | 7 |
| 342 | Neu5Aca2-6Galb1-4GlcNAcb1-2Mana1-3Manb1-4GlcNAcb1-4GlcNAc-Sp12 | 19 | 2 | 9 |
| 555 | (3S)GlcAb1-3Galb1-4GlcNAcb1-3Galb1-4Glc-Sp0 | 19 | 6 | 31 |
| 219 | Fuca1-2Galb1-4(6S)GlcNAcb-Sp8 | 18 | 1 | 5 |
| 419 | Fuca1-2Galb1-3GlcNAcb1-2Mana1-6(Fuca1-2Galb1-3GlcNAcb1-2Mana1-3)Manb1-4GlcNAcb1-4(Fuca1-6)GlcNAcb-Sp22 | 18 | 5 | 25 |
| 426 | GlcNAcb1-6(GlcNAcb1-2)Mana1-6(GlcNAcb1-4)(GlcNAcb1-2Mana1-3)Manb1-4GlcNAcb1-4GlcNAc-Sp21 | 18 | 4 | 22 |
| 470 | Neu5Aca2-3Galb1-4GlcNAcb1-2Mana-Sp0 | 18 | 7 | 39 |
| 521 | Neu5Aca2-3Galb1-3GalNAcb1-4Galb1-4Glcb-Sp0 | 18 | 5 | 27 |
| 25 | (3S)Galb1-4Glcb-Sp8 | 18 | 6 | 31 |
| 33 | (3S)Galb1-4(Fuca1-3)GlcNAc-Sp8 | 18 | 1 | 5 |
| 115 | Gala1-3Galb1-4GlcNAcb-Sp8 | 18 | 4 | 23 |
| 142 | Galb1-3GalNAcb-Sp8 | 18 | 3 | 18 |
| 199 | Glcb1-6Glcb-Sp8 | 18 | 2 | 10 |
| 227 | Neu5Aca2-8Neu5Aca2-8Neu5Aca-Sp8 | 18 | 2 | 10 |
| 353 | KDNa2-3Galb1-3GalNAca-Sp14 | 18 | 4 | 20 |

| | | | | |
|---|---|---|---|---|
| 402 | Gala1-3(Fuca1-2)Galb1-4(Fuca1-3)Glcb-Sp21 | 18 | 3 | 19 |
| 482 | Galb1-3(Fuca1-4)GlcNAcb1-6GalNAca-Sp14 | 18 | 7 | 41 |
| 10 | Neu5Aca-Sp11 | 18 | 6 | 35 |
| 89 | GalNAca1-3(Fuca1-2)Galb-Sp8 | 18 | 1 | 7 |
| 192 | GlcNAcb1-6GalNAca-Sp8 | 18 | 2 | 13 |
| 329 | GalNAca1-3(Fuca1-2)Galb1-4GlcNAcb1-3Galb1-4GlcNAcb-Sp0 | 18 | 1 | 7 |
| 360 | Neu5Aca2-6GlcNAcb1-4GlcNAc-Sp21 | 18 | 1 | 5 |
| 484 | (3S)Galb1-3(Fuca1-4)GlcNAcb-Sp0 | 18 | 3 | 17 |
| 487 | Gala1-3Galb1-4GlcNAcb1-6GalNAca-Sp14 | 18 | 5 | 27 |
| 499 | GlcNAcb1-6(GlcNAcb1-2)Mana1-6(GlcNAcb1-4)(GlcNAcb1-4(GlcNAcb1-2)Mana1-3)Manb1-4GlcNAcb1-4(Fuca1-6)GlcNAc-Sp21 | 18 | 1 | 7 |
| 505 | (3S)GalNAcb1-4(3S)GlcNAc-Sp8 | 18 | 5 | 26 |
| 49 | Neu5,9Ac2a2-6Galb1-4GlcNAcb-Sp8 | 18 | 1 | 7 |
| 150 | Galb1-3GlcNAcb-Sp8 | 18 | 5 | 27 |
| 151 | Galb1-4(Fuca1-3)GlcNAcb-Sp0 | 18 | 1 | 7 |
| 205 | KDNa2-3Galb1-3GlcNAcb-Sp0 | 18 | 1 | 3 |
| 225 | Neu5Aca2-8Neu5Aca2-8Neu5Aca2-3Galb1-4Glcb-Sp0 | 18 | 2 | 14 |
| 298 | Galb1-4GlcNAcb1-6(Galb1-4GlcNAcb1-3)Galb1-4GlcNAc-Sp0 | 18 | 1 | 3 |
| 365 | Gala1-3(Fuca1-2)Galb1-4GlcNAcb1-2Mana1-6(Gala1-3(Fuca1-2)Galb1-4GlcNAcb1-2Mana1-3)Manb1-4GlcNAcb1-4GlcNAcb-Sp20 | 18 | 5 | 26 |
| 396 | Gala1-4Galb1-3GlcNAcb1-2Mana1-6(Gala1-4Galb1-3GlcNAcb1-2Mana1-3)Manb1-4GlcNAcb1-4GlcNAcb-Sp19 | 18 | 6 | 37 |
| 441 | Galb1-4GlcNAcb1-2Mana-Sp0 | 18 | 6 | 34 |
| 495 | GalNAcb1-4(Fuca1-3)(6S)GlcNAcb-Sp8 | 18 | 3 | 16 |
| 532 | GlcNAcb1-3Galb1-4GlcNAcb1-2Mana1-6(GlcNAcb1-3Galb1-4GlcNAcb1-2Mana1-3)Manb1-4GlcNAcb1-4GlcNAcb-Sp25 | 18 | 3 | 19 |
| 541 | Neu5Gca2-8Neu5Aca2-3Galb1-4GlcNAc-Sp0 | 18 | 1 | 7 |
| 580 | Neu5Aca2-6Galb1-4GlcNAcb1-3Galb1-4GlcNAcb1-6(Neu5Aca2-6Galb1-4GlcNAcb1-3Galb1-4GlcNAcb1-3)GalNAca-Sp14 | 18 | 7 | 41 |
| 585 | Galb1-3GalNAcb1-4(Neu5Aca2-8Neu5Aca2-8Neu5Aca2-3)Galb1-4Glcb-Sp21 | 18 | 5 | 27 |
| 4 | GalNAca-Sp8 | 17 | 6 | 36 |
| 5 | GalNAca-Sp15 | 17 | 2 | 13 |
| 28 | (3S)Galb1-3(Fuca1-4)GlcNAcb-Sp8 | 17 | 5 | 29 |
| 122 | Gala1-4Galb1-4Glcb-Sp0 | 17 | 2 | 9 |
| 128 | Galb1-3(Fuca1-4)GlcNAc-Sp0 | 17 | 3 | 14 |
| 144 | Galb1-3GalNAcb1-4(Neu5Aca2-3)Galb1-4Glcb-Sp0 | 17 | 4 | 23 |
| 462 | Glca1-6Glca1-6Glca1-6Glcb-Sp10 | 17 | 3 | 15 |
| 485 | Galb1-4(Fuca1-3)GlcNAcb1-6(Neu5Aca2-6(Neu5Aca2-3Galb1-3)GlcNAcb1-3)Galb1-4Glc-Sp21 | 17 | 5 | 27 |
| 540 | Neu5Aca2-8Neu5Gca2-3Galb1-4GlcNAc-Sp0 | 17 | 2 | 10 |
| 20 | Galb1-4GlcNAcb1-6(Galb1-4GlcNAcb1-3)GalNAc-Sp14 | 17 | 7 | 39 |
| 66 | Fuca1-2Galb1-3GlcNAcb1-3Galb1-4Glcb-Sp10 | 17 | 1 | 8 |
| 91 | GalNAca1-3GalNAcb-Sp8 | 17 | 3 | 19 |
| 118 | Gala1-3Galb-Sp8 | 17 | 2 | 10 |
| 149 | Galb1-3GlcNAcb-Sp0 | 17 | 1 | 7 |
| 166 | Galb1-4GlcNAcb1-6(Galb1-3)GalNAca-Sp8 | 17 | 1 | 5 |
| 197 | Glca1-6Glca1-6Glcb-Sp8 | 17 | 2 | 13 |
| 226 | GalNAcb1-4(Neu5Aca2-8Neu5Aca2-3)Galb1-4Glcb-Sp0 | 17 | 2 | 11 |
| 231 | Neu5Aca2-3Galb1-3GalNAcb1-4(Neu5Aca2-3)Galb1-4Glcb-Sp0 | 17 | 1 | 5 |
| 247 | Neu5Aca2-3Galb1-4(6S)GlcNAcb-Sp8 | 17 | 1 | 7 |
| 321 | Neu5Aca2-3Galb1-4GlcNAcb1-2Mana1-6(Neu5Aca2-6Galb1-4GlcNAcb1-2Mana1-3)Manb1-4GlcNAcb1-4GlcNAcb-Sp12 | 17 | 1 | 8 |
| 510 | (6P)Galb1-4GlcNAcb-SP0 | 17 | 4 | 24 |
| 12 | Galb-Sp8 | 17 | 6 | 38 |
| 129 | Galb1-3(Fuca1-4)GlcNAc-Sp8 | 17 | 2 | 9 |
| 132 | Galb1-4GlcNAcb1-6GalNAc-Sp14 | 17 | 2 | 12 |
| 136 | Neu5Aca2-6(Galb1-3)GalNAca-Sp14 | 17 | 1 | 6 |

| | | | | |
|---|---|---|---|---|
| 252 | Neu5Aca2-3Galb1-4(Fuca1-3)GlcNAcb1-3Galb-Sp8 | 17 | 1 | 6 |
| 277 | Neu5Gca2-3Galb1-3GlcNAcb-Sp0 | 17 | 1 | 3 |
| 281 | Neu5Gca2-6GalNAca-Sp0 | 17 | 2 | 13 |
| 282 | Neu5Gca2-6Galb1-4GlcNAcb-Sp0 | 17 | 1 | 6 |
| 339 | Neu5Aca2-6Galb1-4GlcNAcb1-2Mana1-6(Mana1-3)Manb1-4GlcNAcb1-4GlcNAc-Sp12 | 17 | 2 | 12 |
| 376 | Galb1-3GlcNAcb1-3Galb1-4GlcNAcb1-6(Galb1-3GlcNAcb1-3)Galb1-4Glcb-Sp0 | 17 | 1 | 8 |
| 382 | GlcNAcb1-2Mana1-6(GlcNAcb1-4(GlcNAcb1-2)Mana1-3)Manb1-4GlcNAcb1-4GlcNAc-Sp21 | 17 | 4 | 26 |
| 500 | Galb1-4GlcNAcb1-6(Galb1-4GlcNAcb1-2)Mana1-6(GlcNAcb1-4)Galb1-4GlcNAcb1-4(Gal b1-4GlcNAcb1-2)Mana1-3)Manb1-4GlcNAcb1-4(Fuca1-6)GlcNAc-Sp21 | 17 | 4 | 23 |
| 571 | Neu5Aca2-3Galb1-4GlcNAcb1-3Galb1-4GlcNAcb1-6(Neu5Aca2-3Galb1-4GlcNAcb1-3Galb1-4GlcNAcb1-3)GalNAca-Sp14 | 17 | 3 | 17 |
| 7 | Fuca-Sp9 | 17 | 1 | 6 |
| 41 | (6P)Mana-Sp8 | 17 | 2 | 14 |
| 75 | Fuca1-2Galb1-4GlcNAcb-Sp0 | 17 | 1 | 3 |
| 130 | Fuca1-4(Galb1-3)GlcNAcb-Sp8 | 17 | 1 | 8 |
| 178 | GlcNAcb1-6(GlcNAcb1-3)Galb1-4GlcNAcb-Sp8 | 17 | 1 | 6 |
| 201 | GlcAa-Sp8 | 17 | 3 | 16 |
| 221 | Neu5Aca2-3Galb1-3GalNAca-Sp8 | 17 | 3 | 15 |
| 267 | Neu5Aca2-6Galb1-4GlcNAcb1-3Galb1-4GlcNAcb-Sp0 | 17 | 2 | 10 |
| 270 | Neu5Aca2-6Galb-Sp8 | 17 | 1 | 8 |
| 421 | Galb1-3GlcNAcb1-6(Galb1-3GlcNAcb1-2)Mana1-6(Galb1-3GlcNAcb1-2Mana1-3)Manb1-4GlcNAcb1-4GlcNAcb-Sp19 | 17 | 3 | 19 |
| 428 | Galb1-4GlcNAcb1-2Mana1-6(GlcNAcb1-4)(Galb1-4GlcNAcb1-2Mana1-3)Manb1-4GlcNAcb1-4GlcNAc-Sp21 | 17 | 2 | 12 |
| 446 | GalNAcb1-4Galb1-4Glcb-Sp0 | 17 | 5 | 29 |
| 448 | Gala1-3(Fuca1-2)Galb1-3GlcNAcb1-2Mana1-6(Gala1-3(Fuca1-2)Galb1-3GlcNAcb1-2Mana1-3)Manb1-4GlcNAcb1-4(Fuca1-6)GlcNAcb-Sp22 | 17 | 5 | 29 |
| 469 | Neu5Aca2-6Galb1-4GlcNAcb1-6(Galb1-3GlcNAcb1-3)Galb1-4Glcb-Sp21 | 17 | 3 | 21 |
| 514 | Gala1-3(Fuca1-2)Galb1-4GlcNAcb1-2Mana-Sp0 | 17 | 4 | 24 |
| 46 | Neu5Aca2-3(6S)Galb1-4GlcNAcb-Sp8 | 16 | 4 | 24 |
| 54 | Galb1-4GlcNAcb1-2Mana1-6(Galb1-4GlcNAcb1-2Mana1-3)Manb1-4GlcNAcb1-4GlcNAcb-Sp12 | 16 | 1 | 6 |
| 71 | Fuca1-2Galb1-4(Fuca1-3)GlcNAcb-Sp0 | 16 | 1 | 6 |
| 87 | GalNAca1-3(Fuca1-2)Galb1-4Glcb-Sp0 | 16 | 1 | 8 |
| 114 | Gala1-3Galb1-3GlcNAcb-Sp0 | 16 | 3 | 18 |
| 202 | GlcAb-Sp8 | 16 | 3 | 21 |
| 233 | Neu5Aca2-3GalNAca-Sp8 | 16 | 3 | 16 |
| 251 | Neu5Aca2-3Galb1-4(Fuca1-3)GlcNAcb-Sp8 | 16 | 2 | 9 |
| 260 | Neu5Aca2-3Galb1-4Glcb-Sp8 | 16 | 2 | 14 |
| 328 | GalNAcb1-3Gala1-4Galb1-4GlcNAcb1-3Galb1-4Glcb-Sp0 | 16 | 3 | 16 |
| 403 | Galb1-4GlcNAcb1-6(Neu5Aca2-6Galb1-3GlcNAcb1-3)Galb1-4Glc-Sp21 | 16 | 3 | 20 |
| 478 | Neu5Aca2-3Galb1-3GlcNAcb1-2Mana1-6(GlcNAcb1-4)(Neu5Aca2-3Galb1-3GlcNAcb1-2Mana1-3)Manb1-4GlcNAcb1-4GlcNAc-Sp21 | 16 | 3 | 20 |
| 31 | (3S)Galb1-3GlcNAcb-Sp8 | 16 | 2 | 11 |
| 51 | Mana1-6(Mana1-3)Manb1-4GlcNAcb1-4GlcNAcb-Sp13 | 16 | 3 | 16 |
| 62 | Fuca1-2Galb1-3GalNAca-Sp14 | 16 | 0 | 0 |
| 93 | GalNAca1-4(Fuca1-2)Galb1-4GlcNAcb-Sp8 | 16 | 1 | 9 |
| 107 | Gala1-3(Fuca1-2)Galb-Sp8 | 16 | 2 | 10 |
| 146 | Galb1-3Galb-Sp8 | 16 | 2 | 15 |
| 168 | Galb1-4GlcNAcb-Sp0 | 16 | 1 | 7 |
| 185 | GlcNAcb1-3Galb1-4Glcb-Sp0 | 16 | 0 | 0 |
| 269 | Neu5Aca2-6Galb1-4Glcb-Sp8 | 16 | 3 | 21 |
| 315 | Galb1-4GlcNAcb1-2Mana1-6(Neu5Aca2-6Galb1-4GlcNAcb1-2Mana1-3)Manb1-4GlcNAcb1-4GlcNAcb-Sp12 | 16 | 1 | 7 |

| | | | | |
|---|---|---|---|---|
| 340 | Mana1-6(Neu5Aca2-6Galb1-4GlcNAcb1-2Mana1-3)Manb1-4GlcNAcb1-4GlcNAc-Sp12 | 16 | 1 | 9 |
| 391 | Galb1-4GlcNAcb1-2Mana1-6(GlcNAcb1-2Mana1-3)Manb1-4GlcNAcb1-4GlcNAc-Sp12 | 16 | 7 | 44 |
| 442 | Fuca1-2Galb1-4GlcNAcb1-6(Fuca1-2Galb1-4GlcNAcb1-3)GalNAc-Sp14 | 16 | 2 | 14 |
| 454 | Neu5Aca2-3Galb1-4GlcNAcb1-6(Neu5Aca2-3Galb1-4GlcNAcb1-2)Mana1-6(GlcNAcb1-4)(Neu5Aca2-3Galb1-4GlcNAcb1-2Mana1-3)Manb1-4GlcNAcb1-4GlcNAcb-Sp21 | 16 | 2 | 15 |
| 523 | Galb1-4GlcNAcb1-2 Mana1-6(GlcNAcb1-4)(Galb1-4GlcNAcb1-2Mana1-3)Manb1-4GlcNAcb1-4(Fuca1-6)GlcNAc-Sp21 | 16 | 1 | 7 |
| 43 | (6S)Galb1-4Glcb-Sp8 | 16 | 2 | 13 |
| 154 | Galb1-4(Fuca1-3)GlcNAcb1-3Galb1-4(Fuca1-3)GlcNAcb1-3Galb1-4(Fuca1-3)GlcNAcb-Sp0 | 16 | 2 | 11 |
| 156 | Galb1-4(6S)Glcb-Sp8 | 16 | 2 | 10 |
| 238 | Neu5Aca2-3Galb1-4(Neu5Aca2-3Galb1-3)GlcNAcb-Sp8 | 16 | 1 | 3 |
| 308 | MurNAcb1-4GlcNAcb-Sp10 | 16 | 5 | 30 |
| 337 | GlcNAca1-4Galb1-4GlcNAcb1-3Galb1-4GlcNAcb-Sp0 | 16 | 1 | 3 |
| 405 | Neu5Aca2-3Galb1-3GalNAcb1-4(Neu5Aca2-8Neu5Aca2-3)Galb1-4Glcb-Sp0 | 16 | 4 | 28 |
| 430 | Galb1-4GlcNAcb1-6(Galb1-4GlcNAcb1-2)Mana1-6(GlcNAcb1-4)(Galb1-4GlcNAcb1-2Mana1-3)Manb1-4GlcNAcb1-4GlcNAc-Sp21 | 16 | 3 | 16 |
| 504 | (6S)GalNAcb1-4GlcNAc-Sp8 | 16 | 3 | 22 |
| 511 | GalNAca1-3(Fuca1-2)Galb1-4GlcNAcb1-6GalNAc-Sp14 | 16 | 2 | 14 |
| 575 | Neu5Aca2-3Galb1-4GlcNAcb1-3Galb1-4GlcNAcb1-6(Galb1-3)GalNAca-Sp14 | 16 | 5 | 32 |
| 56 | Neu5Aca2-6Galb1-4GlcNAcb1-2Mana1-6(Neu5Aca2-6Galb1-4GlcNAcb1-2Man-a1-3)Manb1-4GlcNAcb1-4GlcNAcb-Sp21 | 16 | 1 | 4 |
| 86 | GalNAca1-3(Fuca1-2)Galb1-4GlcNAcb-Sp8 | 16 | 1 | 4 |
| 307 | GlcNAcb1-4GlcNAcb-Sp12 | 16 | 1 | 8 |
| 332 | GlcNAca1-4Galb1-4GlcNAcb1-3Galb1-4GlcNAcb1-3Galb1-4GlcNAcb-Sp0 | 16 | 1 | 6 |
| 345 | Mana1-6(Galb1-4GlcNAcb1-2Mana1-3)Manb1-4GlcNAcb1-4GlcNAcb-Sp12 | 16 | 2 | 11 |
| 383 | Fuca1-2Galb1-3GalNAca1-3(Fuca1-2)Galb1-4Glcb-Sp0 | 16 | 6 | 39 |
| 447 | GalNAca1-3(Fuca1-2)Galb1-4GlcNAcb1-2Mana1-6(GalNAca1-3(Fuca1-2)Galb1-4GlcNAcb1-2Mana1-3)Manb1-4GlcNAcb1-4(Fuca1-6)GlcNAcb-Sp22 | 16 | 1 | 8 |
| 460 | Gala1-3(Fuca1-2)Galb1-3GalNAca-Sp8 | 16 | 3 | 20 |
| 479 | Neu5Aca2-6Galb1-4GlcNAcb1-6(Fuca1-2Galb1-4(Fuca1-3)GlcNAcb1-3)Galb1-4Glc-Sp21 | 16 | 7 | 43 |
| 498 | GalNAca1-3(Fuca1-2)Galb1-3GlcNAcb1-6GalNAca-Sp14 | 16 | 2 | 13 |
| 542 | Neu5Gca2-8Neu5Gca2-3Galb1-4GlcNAcb1-3Galb1-4GlcNAc-Sp0 | 16 | 4 | 25 |
| 80 | Fuca1-4GlcNAcb-Sp8 | 15 | 3 | 18 |
| 127 | Galb1-3GlcNAcb1-3Galb1-4(Fuca1-3)GlcNAcb-Sp0 | 15 | 4 | 26 |
| 143 | Galb1-3GalNAcb1-3Gala1-4Galb1-4Glcb-Sp0 | 15 | 2 | 10 |
| 236 | Neu5Aca2-3Galb1-3(Fuca1-4)GlcNAcb-Sp8 | 15 | 1 | 6 |
| 243 | Neu5Aca2-3Galb1-3GalNAcb1-3Gala1-4Galb1-4Glcb-Sp0 | 15 | 5 | 30 |
| 425 | GlcNAcb1-2Mana1-6(GlcNAcb1-4)(GlcNAcb1-4(GlcNAcb1-2)Mana1-3)Manb1-4GlcNAcb1-4GlcNAc-Sp21 | 15 | 2 | 12 |
| 490 | Gala1-3(Fuca1-2)Galb1-4GlcNAcb1-6GalNAca-Sp14 | 15 | 5 | 35 |
| 491 | Fuca1-2Galb1-4GlcNAcb1-2Mana-Sp0 | 15 | 3 | 17 |
| 496 | (3S)GalNAcb1-4(Fuca1-3)GlcNAcb-Sp8 | 15 | 2 | 10 |
| 507 | (3S)GalNAcb1-4GlcNAc-Sp8 | 15 | 3 | 19 |
| 155 | Galb1-4(6S)Glcb-Sp0 | 15 | 2 | 12 |
| 494 | Neu5Aca2-6GalNAcb1-4(6S)GlcNAcb-Sp8 | 15 | 6 | 37 |
| 55 | Neu5Aca2-6Galb1-4GlcNAcb1-2Mana1-6(Neu5Aca2-6Galb1-4GlcNAcb1-2Mana1-3)Manb1-4GlcNAcb1-4GlcNAcb-Sp12 | 15 | 1 | 6 |
| 198 | Glcb1-4Glcb-Sp8 | 15 | 1 | 6 |
| 203 | GlcAb1-3Galb-Sp8 | 15 | 2 | 14 |
| 224 | GalNAcb1-4(Neu5Aca2-8Neu5Aca2-8Neu5Aca2-3)Galb1-4Glcb-Sp0 | 15 | 2 | 10 |
| 292 | 4S(3S)Galb1-4GlcNAcb-Sp0 | 15 | 6 | 37 |
| 327 | Gala1-4Galb1-4GlcNAcb1-3Galb1-4Glcb-Sp0 | 15 | 1 | 6 |

| | | | | |
|---|---|---|---|---|
| 330 | GalNAca1-3(Fuca1-2)Galb1-4GlcNAcb1-3Galb1-4GlcNAcb1-3Galb1-4GlcNAcb-Sp0 | 15 | 4 | 27 |
| 358 | Galb1-4GlcNAcb1-2Mana1-6(Mana1-3)Manb1-4GlcNAcb1-4GlcNAcb-Sp12 | 15 | 2 | 10 |
| 394 | Galb1-4(Fuca1-3)GlcNAcb1-3GalNAca-Sp14 | 15 | 2 | 12 |
| 410 | Gala1-3(Fuca1-2)Galb1-4(Fuca1-3)GlcNAcb1-3GalNAc-Sp14 | 15 | 6 | 38 |
| 506 | GalNAcb1-4(6S)GlcNAc-Sp8 | 15 | 3 | 23 |
| 554 | Galb1-3GlcNAcb1-6(Galb1-3)GalNAc-Sp14 | 15 | 4 | 24 |
| 76 | Fuca1-2Galb1-4GlcNAcb-Sp8 | 15 | 1 | 7 |
| 175 | GlcNAcb1-2Galb1-3GalNAca-Sp8 | 15 | 1 | 4 |
| 211 | Mana1-6(Mana1-3)Mana-Sp9 | 15 | 2 | 16 |
| 215 | Manb1-4GlcNAcb-Sp0 | 15 | 2 | 12 |
| 309 | Mana1-6Manb-Sp10 | 15 | 1 | 9 |
| 322 | Galb1-4(Fuca1-3)GlcNAcb1-2Mana1-6(Galb1-4(Fuca1-3)GlcNAcb1-2Mana1-3)Manb1-4GlcNAcb1-4GlcNAcb-Sp20 | 15 | 3 | 23 |
| 371 | Neu5Aca2-6Galb1-4GlcNAcb1-3GalNAc-Sp14 | 15 | 1 | 4 |
| 377 | Galb1-4(Fuca1-3)GlcNAcb1-6(Galb1-3GlcNAcb1-3)Galb1-4Glc-Sp21 | 15 | 2 | 12 |
| 387 | GalNAca1-3(Fuca1-2)Galb1-3GalNAca1-3(Fuca1-2)Galb1-4GlcNAcb-Sp0 | 15 | 3 | 17 |
| 513 | Gala1-3Galb1-4GlcNAcb1-2Mana-Sp0 | 15 | 1 | 9 |
| 90 | GalNAca1-3(Fuca1-2)Galb-Sp18 | 14 | 2 | 11 |
| 258 | Fuca1-2Galb1-4(6S)Glcb-Sp0 | 14 | 2 | 11 |
| 489 | Fuca1-2(6S)Galb1-3GlcNAcb-Sp0 | 14 | 2 | 13 |
| 522 | GlcNAcb1-2 Mana1-6(GlcNAcb1-4)(GlcNAcb1-2Mana1-3)Manb1-4GlcNAcb1-4(Fuca1-6)GlcNAc-Sp21 | 14 | 1 | 7 |
| 528 | GalNAca1-3(Fuca1-2)Galb1-3GalNAcb1-3Gala1-4Galb1-4Glc-Sp21 | 14 | 2 | 12 |
| 533 | Galβ1-4GlcNAcβ1-3Galβ1-4GlcNAcβ1-2Manα1-6(Galβ1-4GlcNAcβ1-3Galβ1-4GlcNAcβ1-2Manα1-3)Manβ1-4GlcNAcβ1-4GlcNAcβ-Sp12 | 14 | 4 | 26 |
| 559 | Neu5Aca2-8Neu5Aca2-3Galb1-3GalNAcb1-4(Neu5Aca2-3)Galb1-4Glc-Sp21 | 14 | 5 | 32 |
| 563 | Galb1-4GlcNAcb1-3Galb1-4GlcNAcb1-3Galb1-4GlcNAcb1-6(Galb1-4GlcNAcb1-3Galb1-4GlcNAcb1-3Galb1-4GlcNAb1-2)Mana1-6(Galb1-4GlcNAcb1-3Galb1-4GlcNAcb1-3Galb1-4GlcNAcb1-2Mana1-3)Manb1-4GlcNAcb1-4(Fuca1-6)GlcNAcb-Sp24 | 14 | 4 | 25 |
| 568 | GlcNAcb1-3Galb1-4GlcNAcb1-3GalNAca-Sp14 | 14 | 1 | 9 |
| 569 | GlcNAcb1-3Galb1-4GlcNAcb1-6(Galb1-3)GalNAca-Sp14 | 14 | 1 | 7 |
| 139 | Galb1-3GalNAca-Sp8 | 14 | 1 | 10 |
| 140 | Galb1-3GalNAca-Sp14 | 14 | 3 | 20 |
| 148 | Galb1-3GlcNAcb1-3Galb1-4Glcb-Sp10 | 14 | 1 | 6 |
| 204 | GlcAb1-6Galb-Sp8 | 14 | 1 | 8 |
| 232 | Neu5Aca2-6(Neu5Aca2-3)GalNAca-Sp8 | 14 | 1 | 6 |
| 234 | Neu5Aca2-3GalNAcb1-4GlcNAcb-Sp0 | 14 | 2 | 17 |
| 343 | Galb1-4GlcNAcb1-2Mana1-3Manb1-4GlcNAcb1-4GlcNAc-Sp12 | 14 | 1 | 6 |
| 392 | Neu5Aca2-3Galb1-3GlcNAcb1-3GalNAca-Sp14 | 14 | 2 | 17 |
| 433 | Galb1-6Galb-Sp10 | 14 | 3 | 21 |
| 473 | Neu5Aca2-6Galb1-4 GlcNAcb1-6(Neu5Aca2-6Galb1-4GlcNAcb1-3)GalNAca-Sp14 | 14 | 2 | 13 |
| 503 | (6S)(4S)GalNAcb1-4GlcNAc-Sp8 | 14 | 7 | 48 |
| 509 | Galb1-4(6P)GlcNAcb-Sp0 | 14 | 3 | 19 |
| 524 | Galb1-4GlcNAcb1-2 Mana1-6(Galb1-4GlcNAcb1-4)(Galb1-4GlcNAcb1-2Mana1-3)Manb1-4GlcNAcb1-4(Fuca1-6)GlcNAc-Sp21 | 14 | 2 | 13 |
| 543 | Neu5Gca2-8Neu5Gca2-6Galb1-4GlcNAc-Sp0 | 14 | 3 | 25 |
| 551 | GlcNAcb1-3Galb1-4GlcNAcb1-3Galb1-4GlcNAcb1-3Galb1-4GlcNAcb1-3Galb1-4GlcNAcb1-2Mana1-6(GlcNAcb1-3Galb1-4GlcNAcb1-3Galb1-4GlcNAcb1-3Galb1-4GlcNAcb1-3Galb1-4GlcNAcb1-2Mana1-3)Manb1-4GlcNAcb1-4GlcNAcb-Sp25 | 14 | 5 | 34 |
| 42 | (6S)Galb1-4Glcb-Sp0 | 14 | 1 | 7 |
| 116 | Gala1-3Galb1-4Glcb-Sp0 | 14 | 2 | 11 |
| 117 | Gala1-3Galb1-4Glc-Sp10 | 14 | 5 | 36 |
| 173 | GlcNAca1-3Galb1-4GlcNAcb-Sp8 | 14 | 2 | 12 |

| | | | | |
|---|---|---|---|---|
| 191 | GlcNAcb1-4GlcNAcb1-4GlcNAcb-Sp8 | 14 | 3 | 21 |
| 213 | Mana1-6(Mana1-3)Mana1-6(Mana1-2Mana1-3)Manb1-4GlcNAcb1-4GlcNAcb-Sp12 | 14 | 2 | 12 |
| 268 | Neu5Aca2-6Galb1-4Glcb-Sp0 | 14 | 2 | 12 |
| 289 | Galb1-4GlcNAcb1-3Galb1-3GlcNAcb-Sp0 | 14 | 3 | 25 |
| 351 | KDNa2-6Galb1-4GlcNAc-Sp0 | 14 | 6 | 43 |
| 508 | (4S)GalNAcb-Sp10 | 14 | 1 | 4 |
| 529 | Gala1-3(Fuca1-2)Galb1-3GalNAcb1-3Gala1-4Galb1-4Glc-Sp21 | 14 | 3 | 23 |
| 120 | Gala1-4Galb1-4GlcNAcb-Sp0 | 14 | 2 | 13 |
| 182 | GlcNAcb1-3Galb1-4GlcNAcb-Sp0 | 14 | 3 | 23 |
| 210 | Mana1-2Mana1-6(Mana1-2Mana1-3)Mana1-6(Mana1-2Mana1-2Mana1-3)Manb1-4GlcNAcb1-4GlcNAcb-Sp12 | 14 | 3 | 25 |
| 256 | Neu5Aca2-3Galb1-4GlcNAcb-Sp8 | 14 | 2 | 13 |
| 259 | Neu5Aca2-3Galb1-4Glcb-Sp0 | 14 | 2 | 14 |
| 294 | (6P)Glcb-Sp10 | 14 | 1 | 4 |
| 326 | Neu5Aca2-6Galb1-4GlcNAcb1-3Galb1-4GlcNAcb1-3Galb1-4GlcNAcb-Sp0 | 14 | 1 | 4 |
| 393 | Fuca1-2Galb1-4GlcNAcb1-3GalNAca-Sp14 | 14 | 3 | 20 |
| 147 | Galb1-3GlcNAcb1-3Galb1-4GlcNAcb-Sp0 | 13 | 3 | 25 |
| 164 | Galb1-4GlcNAcb1-3Galb1-4Glcb-Sp0 | 13 | 1 | 7 |
| 176 | GlcNAcb1-6(GlcNAcb1-3)GalNAca-Sp8 | 13 | 1 | 9 |
| 297 | Neu5Aca2-6Galb1-4GlcNAcb1-2Mana1-6(Galb1-4GlcNAcb1-2Mana1-3)Manb1-4GlcNAcb1-4GlcNAcb-Sp12 | 13 | 1 | 7 |
| 335 | GlcNAca1-4Galb1-4GlcNAcb1-3Galb1-4Glcb-Sp0 | 13 | 2 | 17 |
| 341 | Neu5Aca2-6Galb1-4GlcNAcb1-2Mana1-6Manb1-4GlcNAcb1-4GlcNAc-Sp12 | 13 | 1 | 7 |
| 434 | Neu5Aca2-3Galb1-4GlcNAcb1-3Galb-Sp8 | 13 | 5 | 36 |
| 456 | Neu5Aca2-6Galb1-4GlcNAcb1-2Mana1-6(GlcNAcb1-4)(Neu5Aca2-6Galb1-4GlcNAcb1-2Mana1-3)Manb1-4GlcNAcb1-4GlcNAcb-Sp21 | 13 | 4 | 32 |
| 463 | Glca1-4Glca1-4Glca1-4Glcb-Sp10 | 13 | 4 | 26 |
| 502 | Galb1-3(6S)GlcNAcb-Sp8 | 13 | 10 | 76 |
| 565 | Galb1-4GlcNAcb1-3Galb1-4GlcNAcb1-6(Galb1-3)GalNAca-Sp14 | 13 | 4 | 32 |
| 59 | Fuca1-2Galb1-3GalNAcb1-3Gala1-4Galb1-4Glcb-Sp9 | 13 | 1 | 9 |
| 67 | Fuca1-2Galb1-3GlcNAcb-Sp0 | 13 | 3 | 20 |
| 244 | Neu5Aca2-3Galb1-3GlcNAcb1-3Galb1-4GlcNAcb-Sp0 | 13 | 2 | 17 |
| 278 | Neu5Gca2-3Galb1-4(Fuca1-3)GlcNAcb-Sp0 | 13 | 2 | 14 |
| 284 | Neu5Aca2-3Galb1-4GlcNAcb1-6(Galb1-3)GalNAca-Sp14 | 13 | 1 | 6 |
| 302 | GalNAcb1-3Galb-Sp8 | 13 | 2 | 17 |
| 311 | Mana1-2Mana1-6(Mana1-3)Mana1-6(Mana1-2Mana1-2Mana1-3)Mana-Sp9 | 13 | 1 | 11 |
| 314 | Neu5Aca2-6Galb1-4GlcNAcb1-2Mana1-6(Neu5Aca2-3Galb1-4GlcNAcb1-2Mana1-3)Manb1-4GlcNAcb1-4GlcNAcb-Sp12 | 13 | 2 | 14 |
| 317 | Neu5Aca2-8Neu5Aca2-8Neu5Acb-Sp8 | 13 | 2 | 17 |
| 415 | Fuca1-2Galb1-3GlcNAcb1-3GalNAc-Sp14 | 13 | 6 | 44 |
| 422 | Galb1-4GlcNAcb1-6(Fuca1-2Galb1-3GlcNAcb1-3)Galb1-4Glc-Sp21 | 13 | 2 | 13 |
| 431 | Galb1-4GlcNAcb1-6(Galb1-4GlcNAcb1-2)Mana1-6(GlcNAcb1-4)(Galb1-4GlcNAcb1-4(Galb1-4GlcNAcb1-2)Mana1-3)Manb1-4GlcNAcb1-4GlcNAc-Sp21 | 13 | 1 | 11 |
| 458 | Neu5Aca2-6Galb1-4GlcNAcb1-6(Neu5Aca2-6Galb1-4GlcNAcb1-2)Mana1-6(GlcNAcb1-4)(Neu5Aca2-6Galb1-4GlcNAcb1-2Mana1-3)Manb1-4GlcNAcb1-4GlcNAcb-Sp21 | 13 | 0 | 0 |
| 481 | Gala1-3Galb1-3GlcNAcb1-6GalNAca-Sp14 | 13 | 2 | 17 |
| 517 | Gala1-3(Fuca1-2)Galb1-3GlcNAcb1-6GalNAc-Sp14 | 13 | 5 | 37 |
| 519 | Gala1-3Galb1-3GlcNAcb1-2Mana-Sp0 | 13 | 2 | 15 |
| 162 | Galb1-4GlcNAcb1-3Galb1-4GlcNAcb1-3Galb1-4GlcNAcb-Sp0 | 13 | 3 | 23 |
| 400 | Galb1-3GlcNAca1-6Galb1-4GlcNAcb-Sp0 | 13 | 2 | 13 |
| 423 | Fuca1-3GlcNAcb1-6(Galb1-4GlcNAcb1-3)Galb1-4Glc-Sp21 | 13 | 2 | 16 |
| 432 | Galb1-4Galb-Sp10 | 13 | 3 | 24 |
| 483 | Neu5Aca2-3Galb1-3GlcNAcb1-6GalNAca-Sp14 | 13 | 1 | 4 |

| | | | | |
|---|---|---|---|---|
| 544 | Neu5Aca2-8Neu5Aca2-3Galb1-4GlcNAc-Sp0 | 13 | 3 | 24 |
| 92 | GalNAca1-3Galb-Sp8 | 13 | 3 | 23 |
| 163 | Galb1-4GlcNAcb1-3Galb1-4GlcNAcb-Sp0 | 13 | 2 | 19 |
| 172 | Galb1-4Glcb-Sp8 | 13 | 2 | 15 |
| 222 | Neu5Aca2-3Galb1-3GalNAca-Sp14 | 13 | 2 | 14 |
| 229 | GalNAcb1-4(Neu5Aca2-3)Galb1-4GlcNAcb-Sp8 | 13 | 3 | 24 |
| 230 | GalNAcb1-4(Neu5Aca2-3)Galb1-4Glcb-Sp0 | 13 | 3 | 27 |
| 306 | GlcNAcb1-4GlcNAcb-Sp10 | 13 | 1 | 10 |
| 417 | GalNAca1-3(Fuca1-2)Galb1-3GlcNAcb1-3GalNAc-Sp14 | 13 | 2 | 14 |
| 520 | GalNAcb1-4GlcNAcb1-2Mana-Sp0 | 13 | 1 | 8 |
| 527 | GlcNAcb1-3Galb1-4GlcNAcb1-6(GlcNAcb1-3)Galb1-4GlcNAc-Sp0 | 13 | 1 | 10 |
| 125 | Galb1-2Galb-Sp8 | 12 | 1 | 4 |
| 304 | Neu5Aca2-6Galb1-4GlcNAcb1-2Mana1-6(GlcNAcb1-2Mana1-3)Manb1-4GlcNAcb1-4GlcNAcb-Sp12 | 12 | 1 | 8 |
| 512 | Neu5Aca2-6Galb1-4GlcNAcb1-2Man-Sp0 | 12 | 3 | 20 |
| 550 | GalNAcb1-4GlcNAcb1-3GalNAcb1-4GlcNAcb-Sp0 | 12 | 1 | 4 |
| 63 | Fuca1-2Galb1-3GalNAcb1-4(Neu5Aca2-3)Galb1-4Glcb-Sp0 | 12 | 4 | 33 |
| 77 | Fuca1-2Galb1-4Glcb-Sp0 | 12 | 1 | 7 |
| 152 | Galb1-4(Fuca1-3)GlcNAcb-Sp8 | 12 | 4 | 32 |
| 183 | GlcNAcb1-3Galb1-4GlcNAcb-Sp8 | 12 | 3 | 23 |
| 283 | Neu5Gca-Sp8 | 12 | 1 | 12 |
| 453 | Neu5Aca2-3Galb1-4GlcNAcb1-4Mana1-6(GlcNAcb1-4)(Neu5Aca2-3Galb1-4GlcNAcb1-4(Neu5Aca2-3Galb1-4GlcNAcb1-2)Mana1-3)Manb1-4GlcNAcb1-4GlcNAcb-Sp21 | 12 | 3 | 25 |
| 464 | Neu5Aca2-3Galb1-4GlcNAcb1-6(Neu5Aca2-3Galb1-4GlcNAcb1-3)GalNAca-Sp14 | 12 | 4 | 30 |
| 518 | Neu5Aca2-3Galb1-3GlcNAcb1-2Mana-Sp0 | 12 | 2 | 15 |
| 549 | GalNAcb1-3GlcNAcb-Sp0 | 12 | 1 | 12 |
| 235 | Neu5Aca2-3Galb1-3(6S)GlcNAc-Sp8 | 12 | 4 | 37 |
| 257 | Neu5Aca2-3Galb1-4GlcNAcb1-3Galb1-4GlcNAcb-Sp0 | 12 | 2 | 15 |
| 279 | Neu5Gca2-3Galb1-4GlcNAcb-Sp0 | 12 | 4 | 31 |
| 305 | GlcNAcb1-3Man-Sp10 | 12 | 2 | 16 |
| 310 | Mana1-6(Mana1-3)Mana1-6(Mana1-3)Manb-Sp10 | 12 | 2 | 18 |
| 312 | Mana1-2Mana1-6(Mana1-2Mana1-3)Mana1-6(Mana1-2Mana1-2Mana1-3)Mana-Sp9 | 12 | 2 | 15 |
| 324 | Neu5Aca2-6Galb1-4GlcNAcb1-3Galb1-3GlcNAcb-Sp0 | 12 | 1 | 8 |
| 416 | Gala1-3(Fuca1-2)Galb1-3GlcNAcb1-3GalNAc-Sp14 | 12 | 2 | 13 |
| 480 | Galb1-3GlcNAcb1-6GalNAca-Sp14 | 12 | 3 | 21 |
| 79 | Fuca1-3GlcNAcb-Sp8 | 12 | 4 | 35 |
| 165 | Galb1-4GlcNAcb1-3Galb1-4Glcb-Sp8 | 12 | 3 | 22 |
| 290 | Neu5Aca2-3Galb1-3GlcNAcb1-3Galb1-3GlcNAcb-Sp0 | 12 | 4 | 31 |
| 375 | Galb1-3GalNAca1-3(Fuca1-2)Galb1-4GlcNAc-Sp0 | 12 | 4 | 32 |
| 399 | Galb1-3GlcNAcb1-6Galb1-4GlcNAcb-Sp0 | 12 | 2 | 15 |
| 427 | GlcNAcb1-6(GlcNAcb1-2)Mana1-6(GlcNAcb1-4)(GlcNAcb1-4(GlcNAcb1-2)Mana1-3)Manb1-4GlcNAcb1-4GlcNAc-Sp21 | 12 | 5 | 44 |
| 429 | Galb1-4GlcNAcb1-2Mana1-6(GlcNAcb1-4)(Galb1-4GlcNAcb1-4(Galb1-4GlcNAcb1-2)Mana1-3)Manb1-4GlcNAcb1-4GlcNAc-Sp21 | 12 | 3 | 23 |
| 457 | Neu5Aca2-6Galb1-4GlcNAcb1-4Mana1-6(GlcNAcb1-4)(Neu5Aca2-6Galb1-4GlcNAcb1-4(Neu5Aca2-6Galb1-4GlcNAcb1-2)Mana1-3)Manb1-4GlcNAcb1-4GlcNAcb-Sp21 | 12 | 3 | 22 |
| 515 | GalNAca1-3(Fuca1-2)Galb1-4 GlcNAcb1-2Mana-Sp0 | 12 | 1 | 11 |
| 573 | GlcNAcb1-3Galb1-4GlcNAcb1-3Galb1-4GlcNAcb1-3GalNAca-Sp14 | 12 | 2 | 21 |
| 160 | Galb1-4GlcNAcb1-3GalNAc-Sp14 | 11 | 1 | 4 |
| 254 | Neu5Aca2-3Galb1-4GlcNAcb1-3Galb1-4GlcNAcb1-3Galb1-4GlcNAcb-Sp0 | 11 | 2 | 20 |
| 276 | Neu5Gca2-3Galb1-3(Fuca1-4)GlcNAcb-Sp0 | 11 | 4 | 39 |
| 296 | Galb1-3Galb1-4GlcNAcb-Sp8 | 11 | 5 | 41 |
| 323 | Neu5,9Ac2a2-3Galb1-3GlcNAcb-Sp0 | 11 | 1 | 9 |

| | | | | |
|---|---|---|---|---|
| 443 | Gala1-3(Fuca1-2)Galb1-4GlcNAcb1-6(Gala1-3(Fuca1-2)Galb1-4GlcNAcb1-3)GalNAc-Sp14 | 11 | 5 | 40 |
| 572 | Neu5Aca2-6Galb1-4GlcNAcb1-3Galb1-4GlcNAcb1-3GalNAca-Sp14 | 11 | 2 | 15 |
| 444 | GalNAca1-3(Fuca1-2)Galb1-4GlcNAcb1-6(GalNAca1-3(Fuca1-2)Galb1-4GlcNAcb1-3)GalNAc-Sp14 | 11 | 2 | 21 |
| 78 | Fuca1-2Galb-Sp8 | 11 | 3 | 32 |
| 301 | Galb1-4GlcNAcb1-6Galb1-4GlcNAcb-Sp0 | 11 | 5 | 46 |
| 424 | GlcNAcb1-2Mana1-6(GlcNAcb1-4)(GlcNAcb1-2Mana1-3)Manb1-4GlcNAcb1-4GlcNAc-Sp21 | 11 | 1 | 5 |
| 579 | GlcNAcb1-6(Neu5Aca2-3Galb1-3)GalNAca-Sp14 | 11 | 2 | 16 |
| 398 | Gala1-3Galb1-4GlcNAcb1-3GalNAca-Sp14 | 10 | 2 | 17 |
| 567 | Neu5Aca2-3Galb1-4GlcNAcb1-3Galb1-4GlcNAcb1-3GalNAca-Sp14 | 10 | 1 | 9 |
| 34 | (3S)Galb1-4(6S)GlcNAcb-Sp0 | 10 | 2 | 18 |
| 64 | Fuca1-2Galb1-3GalNAcb1-4(Neu5Aca2-3)Galb1-4Glcb-Sp9 | 10 | 3 | 28 |
| 106 | Gala1-3(Fuca1-2)Galb1-4Glcb-Sp0 | 10 | 1 | 14 |
| 189 | GlcNAcb1-4GlcNAcb1-4GlcNAcb1-4GlcNAcb1-4GlcNAcb1-4GlcNAcb1-Sp8 | 10 | 1 | 8 |
| 408 | GalNAca1-3GalNAcb1-3Gala1-4Galb1-4Glcb-Sp0 | 10 | 2 | 22 |
| 455 | Neu5Aca2-3Galb1-4GlcNAcb1-6(Neu5Aca2-3Galb1-4GlcNAcb1-2)Mana1-6(GlcNAcb1-4)(Neu5Aca2-3Galb1-4GlcNAcb1-4(Neu5Aca2-3Galb1-4GlcNAcb1-2)Mana1-3)Manb1-4GlcNAcb1-4GlcNAcb-Sp21 | 10 | 0 | 0 |
| 177 | GlcNAcb1-6(GlcNAcb1-3)GalNAca-Sp14 | 10 | 2 | 24 |
| 291 | Neu5Aca2-3Galb1-4GlcNAcb1-3Galb1-3GlcNAcb-Sp0 | 10 | 3 | 26 |
| 370 | Neu5Aca2-3Galb1-4GlcNAcb1-3GalNAc-Sp14 | 10 | 2 | 23 |
| 561 | GlcNAcb1-3Galb1-4GlcNAcb1-3Galb1-4GlcNAcb1-2Mana1-6(GlcNAcb1-3Galb1-4GlcNAcb1-3Galb1-4GlcNAcb1-2Mana1-3)Manb1-4GlcNAcb1-4(Fuca1-6)GlcNAcb-Sp24 | 10 | 1 | 13 |
| 138 | Neu5Aca2-6(Galb1-3)GlcNAcb1-4Galb1-4Glcb-Sp10 | 10 | 4 | 40 |
| 374 | Galb1-3GalNAca1-3(Fuca1-2)Galb1-4Glc-Sp0 | 10 | 5 | 47 |
| 58 | Fuca1-2Galb1-3GalNAcb1-3Gala-Sp9 | 9 | 3 | 36 |
| 395 | GalNAca1-3GalNAcb1-3Gala1-4Galb1-4GlcNAcb-Sp0 | 9 | 3 | 31 |
| 471 | Neu5Aca2-3Galb1-4GlcNAcb1-6GalNAca-Sp14 | 9 | 2 | 18 |
| 556 | (3S)GlcAb1-3Galb1-4GlcNAcb1-2Mana-Sp0 | 9 | 2 | 20 |
| 82 | GalNAca1-3(Fuca1-2)Galb1-3GlcNAcb-Sp0 | 9 | 1 | 14 |
| 406 | Gala1-3(Fuca1-2)Galb1-4GlcNAcb1-3GalNAca-Sp14 | 9 | 4 | 43 |
| 171 | Galb1-4Glcb-Sp0 | 9 | 1 | 12 |
| 313 | Neu5Aca2-3Galb1-4GlcNAcb1-6(Neu5Aca2-3Galb1-3)GalNAca-Sp14 | 9 | 1 | 12 |
| 548 | GlcNAcb1-3Galb1-4GlcNAcb1-6(GlcNAcb1-3Galb1-3)GalNAca-Sp14 | 9 | 2 | 23 |
| 299 | GlcNAcb1-6(Galb1-4GlcNAcb1-3)Galb1-4GlcNAc-Sp0 | 8 | 1 | 15 |
| 68 | Fuca1-2Galb1-3GlcNAcb-Sp8 | 8 | 2 | 27 |
| 570 | GlcNAcb1-3Galb1-4GlcNAcb1-6(GlcNAcb1-3Galb1-4GlcNAcb1-3)GalNAca-Sp14 | 8 | 3 | 40 |
| 174 | GlcNAca1-6Galb1-4GlcNAcb-Sp8 | 7 | 3 | 36 |
| 184 | GlcNAcb1-3Galb1-4GlcNAcb1-3Galb1-4GlcNAcb-Sp0 | 7 | 2 | 21 |
| 384 | Fuca1-2Galb1-3GalNAca1-3(Fuca1-2)Galb1-4GlcNAcb-Sp0 | 7 | 3 | 40 |
| 81 | Fucb1-3GlcNAcb-Sp8 | 7 | 2 | 26 |
| 200 | G-ol-Sp8 | 7 | 1 | 12 |
| 385 | Galb1-3GlcNAcb1-3GalNAca-Sp14 | 7 | 1 | 20 |
| 190 | GlcNAcb1-4GlcNAcb1-4GlcNAcb1-4GlcNAcb1-4GlcNAcb1-Sp8 | 6 | 1 | 15 |
| 407 | GalNAca1-3(Fuca1-2)Galb1-4GlcNAcb1-3GalNAca-Sp14 | 6 | 3 | 51 |
| 418 | Gala1-3Galb1-3GlcNAcb1-3GalNAc-Sp14 | 5 | 3 | 59 |
| 316 | Neu5Aca2-8Neu5Acb-Sp17 | 3 | 1 | 35 |

**Figure S4: Glycans in microarray stratified by RFU for 029-09 3A04**

| Clone | V | DH | J | CDR3 Length | Predicted Germline CDR3 Sequence | # of Members | Subject(s) with Clone |
|---|---|---|---|---|---|---|---|
| **1** | H3-7 | 1-26 | H3 | 7 | ARKVGDV | 4 | 038 |
| **2** | H3-7 | 6-13 | H4 | 10 | ARAIAAAGSY | 4 | 029-09, 089 |
| **3** | H3-7 | 6-13 | H4 | 10 | ARAIAAAASR | 42 | 029-09, 051-10, 103, 038, 102, 089 |
| **4** | H3-7 | 3-10 | H4 | 10 | AREIAGRGAY | 55 | 051-10/11, 070, 089 |
| **5** | H3-7 | 6-13 | H4 | 10 | ARAIAAADSF | 12 | 103, 102, 089 |
| **6** | H3-7 | 3-22 | H4 | 10 | ARALGSGSCV | 3 | 038 |
| **7** | H3-7 | 6-19 | H4 | 10 | ARAYAGYSSY | 2 | 029-09 |
| **8** | H3-7 | 6-13 | H4 | 10 | AKSLAAADAF | 3 | 038 |
| **9** | H3-7 | None | H4 | 7 | ARRYFDY | 27 | 011-10, 038, 102, 070 |
| **10** | H3-7 | 3-10 | H5 | 10 | CARAYGSGSS | 2 | 082 |
| **11** | H3-7 | 2-21 | H5 | 14 | AGPPPGGEIAMGGS | 2 | 011-10 |
| **12** | L1-44 | N/A | L1 | 11 | AAWDDSLNGYV | 93 | 017-10, 019-10, 034-10, 120, 103, 015, 038, 070, 089 |
| **13** | L1-44 | N/A | L2 | 11 | AAWDDSLNGLV | 13 | 038, 102, 070, 108, 089, 008 |
| **14** | L1-44 | N/A | L2 | 11 | AAWDDSLNGFI | 5 | 102 |
| **15** | L1-44 | N/A | L3 | 11 | AAWDDSLNVWV | 50 | 029-09, 051-10/11, 011-10, 120, 103, 099, 030, 038, 102, 070, 108, 089, 008 |
| **16** | L1-51 | N/A | L1 | 11 | GTWDSSLSAYV | 7 | 019-10, 085, 120, 103, 070, 008 |
| **17** | L1-51 | N/A | L2 | 11 | GTWDSSLSAMV | 16 | 029-09, 103, 038, 089, 008 |
| **18** | L1-51 | N/A | L3 | 11 | GTWDSSLKSIV | 2 | 089 |
| **19** | L1-51 | N/A | L3 | 11 | GTWDSSLSAGM | 6 | 029-09, 008-10, 082, 008 |

**Table S5: Heavy and light chains clonal information.**

| Vaccine | Manufacturer/Name | Propagation Cell Type/Host | Target |
|---|---|---|---|
| 2010 TIV | Novartis – Fluvirin | Egg | Influenza Viruses |
| 2013 QIV | GSK – Fluarix | Egg | Influenza Viruses |
| 2015 QIV | Sanofi – Fluzone | Egg | Influenza Viruses |
| 2015 QIV | Seqirus – Flucelvax | MDCK Cells - Mammalian | Influenza Viruses |
| 2015 QIV | Sanofi – Flublok | Insect cells | Influenza Viruses |
| 2016 QIV | GSK – Fluarix | Egg | Influenza Viruses |
| 2017 QIV | GSK – Fluarix | Egg | Influenza Viruses |
| 2020 QIV | GSK – Fluarix | Egg | Influenza Viruses |
| MMR | Merck – MMR-II | Chicken-derived cell line | Measles, Mumps, Rubella Viruses |
| Rabavert | GSK | Primary chicken fibroblasts | Rabies Virus |
| Ixiaro | Valneva | Vero cell line - Mammalian | Japanese Encephalitis Virus |
| Pneumovax-23 | Merck | *Streptococcus pneumoniae* | *Streptococcus pneumoniae* polysaccharides from 23 serotypes |

**Table S6: Vaccines used in study to test egg-mAb binding potential.**

| Subject ID | Egg Binding mAbs (Yes/No) | Post Vaccination Time Point (Days) |
|---|---|---|
| 029-09 | Yes | 18 |
| 045-09 | No | 21 |
| 047-09 | No | 21 |
| 051-09 | Yes | 21 |
| 008-10 | Yes | 21 |
| 011-10 | Yes | 21 |
| 014-10 | No | 21 |
| 017-10 | Yes | 21 |
| 024-10 | No | 14 |
| 051-10 | Yes | 21 |
| 120-10 | No | 21 |

**Table S7: Serum donors for samples analyzed in Figure 1H and Figure S1D-F.**
